# Crystal structures of the σ_2_ receptor template large-library docking for selective chemotypes active *in vivo*

**DOI:** 10.1101/2021.04.29.441652

**Authors:** Assaf Alon, Jiankun Lyu, Joao M. Braz, Tia A. Tummino, Veronica Craik, Matthew J. O’Meara, Chase M. Webb, Dmytro S. Radchenko, Yurii S. Moroz, Xi-Ping Huang, Yongfeng Liu, Bryan L. Roth, John J. Irwin, Allan I. Basbaum, Brian K. Shoichet, Andrew C. Kruse

## Abstract

The σ_2_ receptor is a poorly understood transmembrane receptor that has attracted intense interest in many areas of biology including cancer imaging, Alzheimer’s disease, schizophrenia, and neuropathic pain. However, little is known regarding the molecular details of the receptor, and few highly selective ligands are available. Here, we report the crystal structure of the σ_2_ receptor in complex with the clinical drug candidate roluperidone and the probe compound PB28. These structures, in turn, templated a large-scale docking screen of 490 million make-on-demand molecules. Of these, 484 compounds were synthesized and tested, prioritizing not only high-ranking docked molecules, but also those with mediocre and poor scores. Overall, 127 compounds with binding affinities superior to 1 μM were identified, all in new chemotypes, 31 of which had affinities superior to 50 nM. Intriguingly, hit rate fell smoothly and monotonically with docking score. Seeking to develop selective and biologically active probe molecules, we optimized three of the original docking hits for potency and for selectivity, achieving affinities in the 3 to 48 nM range and to up to 250-fold selectivity vs. the σ_1_ receptor. Crystal structures of the newly discovered ligands bound to the σ_2_ receptor were subsequently determined, confirming the docked poses. To investigate the contribution of the σ_2_ receptor in pain processing, and to distinguish it from the contribution of the σ_1_ receptor, two potent σ_2_-selective and one potent σ_1_/σ_2_ non-selective ligand were tested for efficacy in a mouse model of neuropathic pain. All three ligands demonstrated time-dependent decreases in mechanical hypersensitivity in the spared nerve injury model, supporting a role for the σ_2_ receptor in nociception, and a possible role for σ_1_/σ_2_ polypharmacology. This study illustrates the opportunities for rapid discovery of *in vivo* active and selective probes to study under-explored areas of biology using structurebased screens of diverse, ultra-large libraries following the elucidation of protein structures.

## Introduction

The σ receptors are integral membrane proteins widely expressed in both the central nervous system and in peripheral tissues, including the liver and kidney. The σ receptors are divided into σ_1_ and σ_2_ “subtypes” based on differences in tissue distribution and in pharmacological profile^1^, but despite their names, the two proteins are entirely unrelated in sequence. Cloned in 1996, the σ_1_ receptor has no paralog within the human genome; its closest homolog of known function is the yeast Δ8,7 sterol isomerase ERG2^2^. Pharmacological studies conducted on σ_1_ knockout mice^3^ showed that the σ_2_ is not a splice variant or other modified form of σ_1_, but rather derives from an unrelated gene. The molecular identity of the σ_2_ receptor remained unknown until recently, when we purified it from calf liver tissue^4^ and showed that it is TMEM97, an ER-resident membrane protein that regulates the sterol transporter NPC1^5,6^. TMEM97 is predicted to be a four-helix bundle protein with both amino and carboxy termini facing the cytoplasm. A member of the EXPERA family^7^, the σ_2_ receptor is distantly related to emopamil-binding protein (EBP), the mammalian Δ8,7 sterol isomerase required for cholesterol synthesis, as well as to other proteins in this family, including TM6SF2, which regulates liver lipid homeostasis^8^.

Despite relatively little being known about the role of σ_2_ in baseline physiological processes, the receptor has been implicated in multiple disease states. For example, the σ_2_ receptor is overexpressed in proliferating cells and in many tumors^9^, and labeled σ_2_ ligands have been proposed as tools for diagnosis and therapy for various cancers^10,11^. Additionally, the σ_2_ receptor was recently identified as an interaction partner of the SARS-CoV-2 viral protein, Orf9c, during cellular infection^12^. Recently it was reported that a ternary complex between the σ_2_ receptor, PGRMC1, and the LDL receptor increases the rate of LDL internalization^13^. Consistent with its high expression in the nervous system, the σ_2_ receptor has also been proposed as a target for the treatment of multiple nervous system disorders. The σ_2_ receptor ligand Elayta (CT1812) is currently in clinical trials for mild to moderate Alzheimer’s disease^14^, and another ligand, roluperidone, (MIN-101) is in clinical development for treatment of the negative symptoms of schizophrenia^15–17^. When tested in animal models, σ_2_ receptor ligands reduce alcohol-withdrawal symptoms^18,19^ and have a neuroprotective effect in brain injury^20^. Finally, recent studies have implicated σ_2_ in chronic pain^19,21,22^, with σ_2_ ligands having anti-allodynic effects in nerve-injury induced models of neuropathic pain. As this is also thought to be true of σ_1_ ligands, and because most σ_2_ ligands cross-react with the σ_1_ receptor, probe ligands selective for σ_2_ over σ_1_ would help illuminate σ_2_ biology and could be leads for novel therapeutics. However, given the relatively recent determination of the σ_2_ receptor’s molecular identity relatively little is known regarding its molecular architecture, ligand recognition, or amenability to methods like virtual screening for ligand identification^23–30^. Here, we employed a biochemical and structural approach combined with computational docking to address these issues.

### Crystallization and structure determination

The human σ_2_ receptor was expressed in *Sf9* insect cells and was extracted with lauryl maltose neopentyl glycol (LMNG) detergent and purified as described^4^. Size exclusion chromatography multi-angle light scattering (SEC-MALS) experiments showed that the receptor is a dimer in solution, and that the presence of ligands did not perturb this oligomeric state (**Supplementary Information Fig. 1a**). As the human σ_2_ receptor did not lend itself to structural studies, further experiments were performed with the bovine σ_2_ receptor, which was more tractable. Circular dichroism (CD) experiments showed that the bovine σ_2_ receptor has a 74% helical content (**Supplementary Information Fig. 1b**), in agreement with secondary structure predictions. CD thermal unfolding experiments demonstrated that the receptor is remarkably stable compared to most mammalian membrane proteins, with a midpoint of the unfolding transition (T_m_) of 54 °C (**Supplementary Information Fig. 1c**). The T_m_ increased by 1-3 °C when the receptor was incubated with various ligands.

Several rounds of construct optimization and of crystallization conditions led to high-quality diffracting crystals of the bovine σ_2_ receptor by the lipidic cubic phase method (**Supplementary Information Fig. 2a-c**). Three data sets were collected, one with receptor bound to the high-affinity non-selective ligand PB28^31^, to a resolution of 2.94 Å, another with receptor bound to the schizophrenia drug candidate roluperidone at a resolution of 2.7 Å, and yet another with the receptor bound to a ligand tentatively modeled as cholesterol at a resolution of 2.6 Å. The σ_2_ receptor has no homologs with known structure, and a BLAST search against the Protein Data Bank yielded no results. A more sensitive hidden Markov model search with HHpred^32^ identified EBP as a distant homolog, consistent with both proteins being members of the EXPERA family^7^. We used the structure of EBP^33^ to build a homology model of the σ_2_ receptor and performed a molecular replacement search that generated a marginally interpretable electron density map. Manual placement of an additional copy of the σ_2_ receptor followed by iterative manual building and reciprocal space refinement led to high-quality final structure (**Supplementary Information Table 1**).

### Overall structure of the σ_2_ receptor

The three initial σ_2_ receptor crystal structures are highly similar, with a backbone root mean square deviation (RMSD) of 0.71 Å. As anticipated from multi-angle light scattering, the structures showed that σ_2_ is an intimately associated homodimer, burying 890 Å^2^ of surface area in dimer interface that is mainly formed by transmembrane helix 3 (TM3; **Fig. 1a**). The two protomers adopt the same conformation (backbone RMSD of 0.34 Å), with each protomer showing the expected four-helix bundle fold with both the amino and carboxy termini facing the same face of the membrane, likely the cytosolic face.

**Figure 1.**
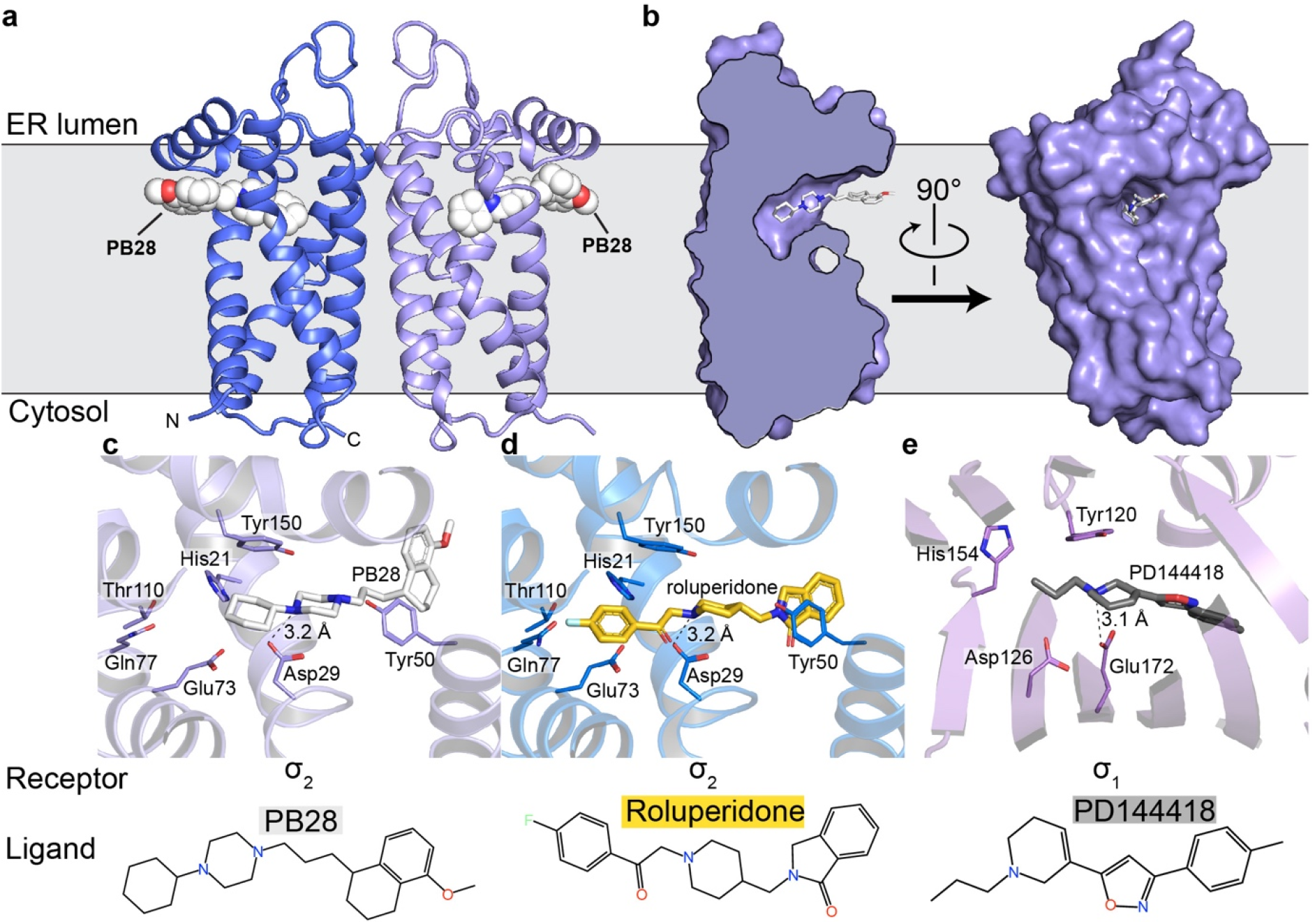
Overall structure of the σ_2_ receptor and binding site ligand recognition. **a**, Structure of the σ_2_ receptor bound to PB28. Amino- and carboxy-termini are indicated. Membrane boundaries were calculated using the PPM server^37^. **b**, Cross-section of the σ_2_ receptor binding pocket (left) and view of the entrance to the binding pocket from the membrane. **c**, View of PB28 binding pose, showing charge–charge interaction with Asp29 (black dotted line) and contacts with other binding pocket residues. **d**, Analogous structure of the roluperidone binding pose. **e**, Structure of the σ_1_ receptor bound to PD144418. Amino acids that serve similar roles and positioned in a similar orientation to amino acids in the σ_2_ receptor are indicated.

The four transmembrane helices of the protein are all kinked due to the presence of proline residues in each helix, creating a cavity slightly above the center of the membrane, nearer the ER-facing side of the membrane. Surprisingly, the ligand-binding cavity is entirely occluded from solvent by extracellular loops 1 and 2, which form a well-ordered cap over the luminal surface of the protein. Asp56, which was shown to be crucial for ligand binding^4^, is located in extracellular loop 1 and is involved in a network of hydrogen bonds likely important for proper folding (**Supplementary Information Fig. 2d)**, hence the deleterious effect of mutating this residue on binding^4^. Instead of opening to the lumen of the ER, the pocket opens laterally into the lipid bilayer (**Fig. 1b**), reminiscent of lipid-binding G protein-coupled receptors^34^. The opening to the binding pocket is lined with hydrophobic and aromatic residues. Ligands may enter through this opening in their neutral, deprotonated form, and then become protonated in the binding site, allowing formation of a salt bridge with the conserved Asp29 (**Fig. 1c-d**). A second highly conserved acidic residue, Asp73, is located 3 Å away from Asp29, suggesting that these residues are hydrogen-bonded to each another, implying that Asp73 is likely protonated.

The two σ receptors are not homologs and do not share the same fold; the σ_2_ receptor is a four-helix bundle, while the σ_1_ receptor has a β-barrel cupin fold^35^. Despite this, the binding pockets of the two receptors are remarkably similar (**Fig. 1c-e**), placing functionally similar amino acids in cognate spatial positions, which is perhaps the result of convergent evolution and explains how two very different folds can share the same pharmacology.

It is noteworthy that both σ receptors are homologs of proteins that catalyze the same step in sterol biosynthesis. The σ_1_ receptor is a homolog of ERG2, the yeast Δ8,7 sterol isomerase; the σ_2_ receptor is a homolog of EBP, the mammalian enzyme that performs the same catalytic step in the biosynthesis of cholesterol. Both EBP and ERG2 rely on two similarly placed acidic residues in their active site for catalysis, which is thought to occur by protonation of the substrate at carbon 9 (C9) followed by proton abstraction from C7, which shifts the double bond into the C8-C7 position. All necessary components for catalysis appear to be present in σ_2_ receptor, in addition to the conserved polar residues His21, Gln77, and Thr110, located at the distal end of the ligand cavity, which may aid in recognizing the hydroxyl moiety of sterols. However, the σ_2_ receptor cannot act as a sterol isomerase. It can neither function *in vivo* to rescue a strain of yeast that lacks ERG2 (**Supplementary Information Fig. 3a**) nor can it function *in vitro* to convert zymostenol to lathosterol (**Supplementary Information Fig. 3b**). The same is true for the σ_1_ receptor, which also has all the residues expected to be required for catalysis and also cannot rescue yeast that lack a sterol isomerase. It was recently reported that Δ8-9 sterols serve as signaling molecules^36^, which may hint at a possible physiological function of the σ receptors as sensors of these molecules evolved from enzymes that would modify them.

### Docking 490 million molecules against the σ_2_ receptor

A prospective docking screen against the σ_2_ receptor had two goals. The first was to discover novel chemotypes with potential σ_2_ selectivity. The second goal was to investigate whether docking scores predict binding likelihood^38^. This second goal we undertook quantitatively, with a 3-fold larger library than previously used^38^. Moreover, the σ_2_ site, with its high propensity to bind ligands, promised a higher dynamic range than the first study against the dopamine receptor. We modeled a hit rate curve as a function docking score against the σ_2_ receptor. Guided by score supplemented by manual selection among the highest-ranking docked molecules, we chose 484 make-on-demand molecules spread among 14 scoring bins covering the highest-ranking (−65 to −57.5 kcal/mol), mid-ranking (−55 to −40 kcal/mol), and low-ranking docking scores (−37.5 to −22.5 kcal/mol). We note that these ranges are true for σ_2_, but they will vary among other targets. Typically, around 40 molecules per scoring bin were selected. Overall, 412 molecules were picked automatically, whereas 72 were picked by manual visual inspection. We tested compounds at 1 μM concentration and defined as “hits” those that displaced greater than 50% [^3^H]-DTG σ_2_ binding. Based on this threshold, 127 of 484 molecules qualified as hits, equal to an overall hit rate of 26% of compounds tested, and a hit rate of over 60% at the peak among the top-ranked molecules (**Fig. 2a**). Plotting docking score vs hit-rate resulted in a curve where, with one key exception (see below), hit-rates fell monotonically with score, with a slope of −4.2%/(kcal/mol) in the inflection region. The curve dropped from a hit rate of 61% at a docking score of about −60 kcal/mol, to 0% at −40 kcal/mol, where it essentially remained for the next 4 (worse) scoring bins (**Fig. 2b**).

**Figure 2.**
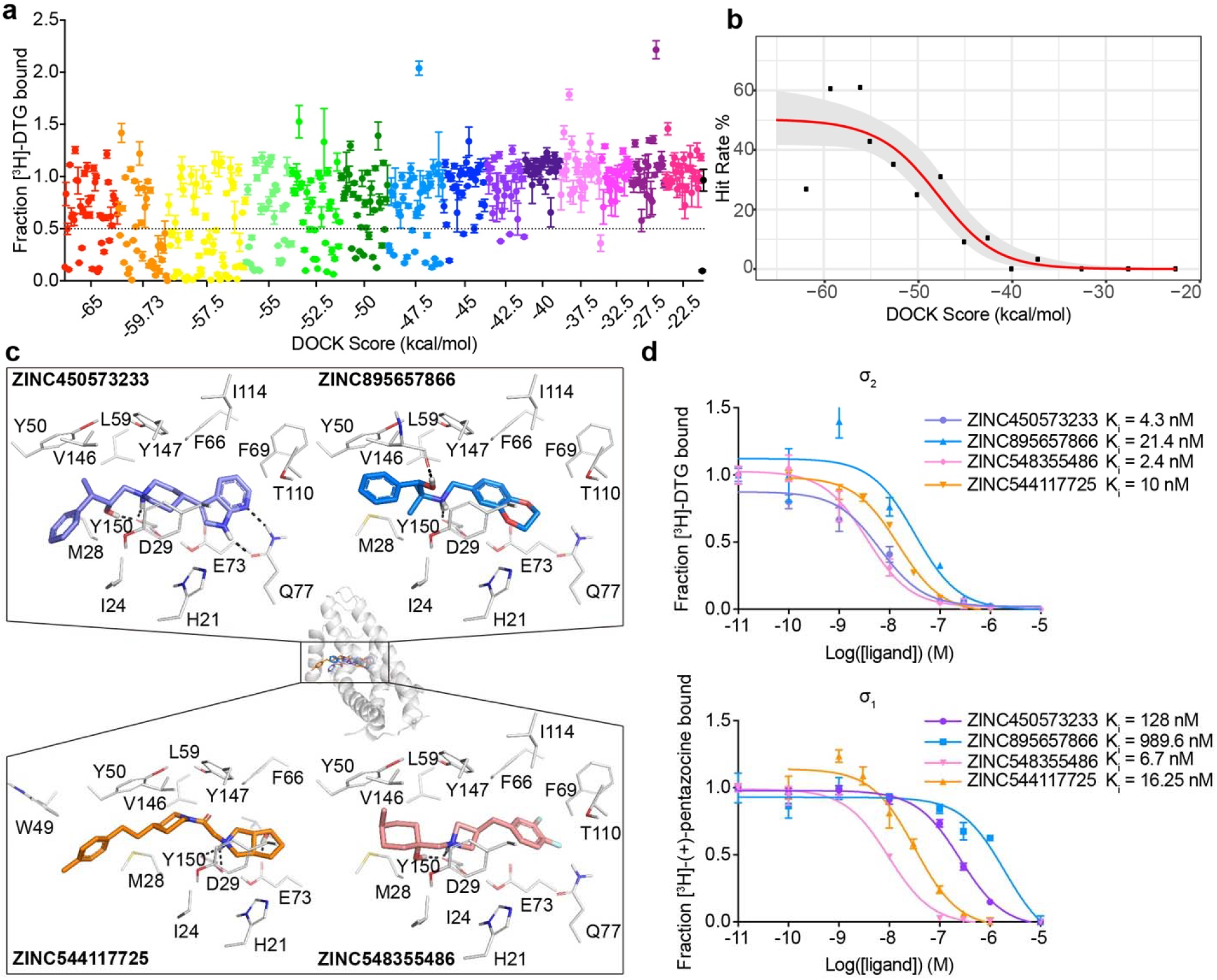
Docking 490 million molecules from ZINC libraries against the σ_2_ receptor. **a**, Displacement of the radioligand [^3^H]-DTG by each of the 484 molecules tested at 1 μM (mean ± SEM of three technical replicates). The molecules are colored by the docking score at which they were selected. Dashed line indicates 50% radioligand displacement. Dot below the dashed line represent confirmed binders, which are diminished with increasing docking score. **b,** The hit-rate of 484 experimentally tested compounds was plotted against docking energy. The hit rate at the top plateau is 50% and at the bottom plateau is 0%, and the docking score (dock_50_) and slope at the maximum (slope50) are −48 kcal mol^-1^ and −4.2% per kcal mol^-1^, respectively. **c**, Docked poses of four representative binders, each a different scaffold. **d**, Doseresponse curves of radioligand displacement assays of the four molecules in **c.** against the σ_2_ receptor (upper panel) and the σ_1_ receptor (lower panel). The data are the mean ± SEM from three technical replicates.

Intriguingly, the hit rate among the very top scoring compounds, 27%, was meaningfully lower than those of the following four bins, and much lower than the 61% hit-rate observed in the 2^nd^-best scoring bin. This dip in the hit-rate curve illuminates holes in the scoring function that can be optimized in future studies. Examination of the 41 molecules (out of 490 million docked) that rank in this very top bin showed that many of these cationic molecules have unexpectedly low desolvation penalties versus electrostatic interaction energy (**Supplementary Information Fig. 4a** and **4b,** the left column). Conversely, their van der Waals energies are undistinguishable among the first three bins (**Supplementary Information Fig. 4c**, the left column). These results implied that the desolvation penalties were underestimated for the molecules in the very top bin, especially for an electrostatics-driven site like the σ_2_ receptor site with a charged anchor residue (Asp29). A possible explanation for this drop in hit rate among the top-ranked molecules was an underestimation of ligand desolvation penalties. DOCK3.7 pre-calculates these energies based on one conformation from among hundreds of that are docked, and not necessarily the conformation that is the highest scoring against a target. Whereas different conformations of the same molecule typically have similar desolvation energies, they can differ, especially for charged molecules. Thus, a molecule can adopt a conformation that optimally complements a receptor electrostatically while only paying the desolvation cost of another, less costly conformation. Accordingly, we recalculated ligand desolvation energy using the docked conformation for all the molecules experimentally tested against σ_2_ and D4 receptors. After recalculation, the molecules in the first, top-scoring bin received much higher desolvation penalties than the following bins (**Supplementary Information Fig. 4d**). This supports the idea that the artifactually favorable scores of the very top-ranked molecules come at least partly from inappropriate desolvation penalties; this, and other holes in the scoring function, merit further investigation and optimization.

Naturally, in addition to testing docking-based prioritization, we were interested in finding new, potent, and selective σ_2_ receptor ligands. To supplement molecules prioritized by score alone, we also picked high-ranking molecules by human inspection^39^. As a comparable number of high-ranking molecules (the first 3 scoring bins) were picked manually and by docking score alone, we could investigate what human inspection added, if anything. In these top three scoring bins, covering 139 molecules (all high scoring), the hit rate by human inspection (67%) was higher than the hit rate by docking score alone (33%) (**Supplementary Information Fig. 5a and 5b**). Human-picked molecules were biased toward higher affinities, at least for those picked from the highest ranks by score: four had K_i_ values < 5 nM and twelve had K_i_ values < 50 nM. For those machine-picked only by score, two had K_i_ values < 5 nM and seven had K_i_ values < 50 nM (**Supplementary Information Fig. 5c**). While it seems clear that the human-picked molecules were more likely to be active, whether their potencies were significantly different from those picked by score alone was less clear. It’s important to emphasize that all of these molecules, human- and machine-prioritized, had highly favorable scores that put them at the very top of the rank-ordered list.

To find potent leads to selective probes for the σ_2_ receptor, we measured concentration-response curves for the primary screen-derived 14 docking hits with the best radioligand displacement at 1 μM. K_i_ values ranged from 2.4 to 68 nM. To measure selectivity, we ran competition binding assays against the σ_1_ receptor, observing K_i_ values from 1.6 nM to 1.5 μM (**Fig. 2d**, **Supplementary Information Table 2 and 3**). Several compounds showed substantial selectivity for σ_2_ over σ_1_, including ZINC450573233 and ZINC895657866, which were 30- and 46-fold selective, respectively.

We sought to improve the affinities of three potent ligands, each in a different scaffold class (**Supplementary Information Fig. 6**) from the docking screen. To do so, 20,000 analogues identified in SmallWorld (https://sw.docking.org/, NextMove Software, Cambridge UK) from a 28 billion make-on-demand library were docked into the σ_2_ site (**Methods**, **Supplementary Table 3**). Of these, 105 high-scoring analogues were sourced and tested. Encouragingly, the affinity of each scaffold was improved by 2-to 18-fold (**Supplementary Information Fig. 6** and **Supplementary Table 3**), and σ_2_ selectivity of two chemotypes improved to 47- and >250-fold (Z1665845742 and Z4857158944), respectively.

### Crystal structures of σ_2_ receptor bound to optimized analogues

To validate our docking poses we determined the crystal structure of the two high affinity analogues Z1241145220 (σ_2_ K_i_ = 3.7 nM; PDB ID: 7M95) and Z4857158944 (σ_2_ K_i_ = 4 nM; PDB ID: 7M96). The electron density maps confirmed the docking predictions, with RMSD values between the crystallized and docked poses of 0.88 and 1.4 Å, respectively (**Fig. 3a**, **Fig. 3b** and **Supplementary Information Table 1**). Newly predicted hydrogen-bond interactions with Gln77 and the backbone carbonyl of Val146, which were not seen in the roluperidone or PB28 complexes, corresponded well between docked and crystallographic poses. The higher resolution of this structure, 2.4 Å, revealed an ordered water molecule in one of the binding site sub-pockets, coordinated by residues His21, Tyr103, and Gln77, and by an azaindole nitrogen in Z1241145220 (**Fig. 3b**).

**Figure 3.**
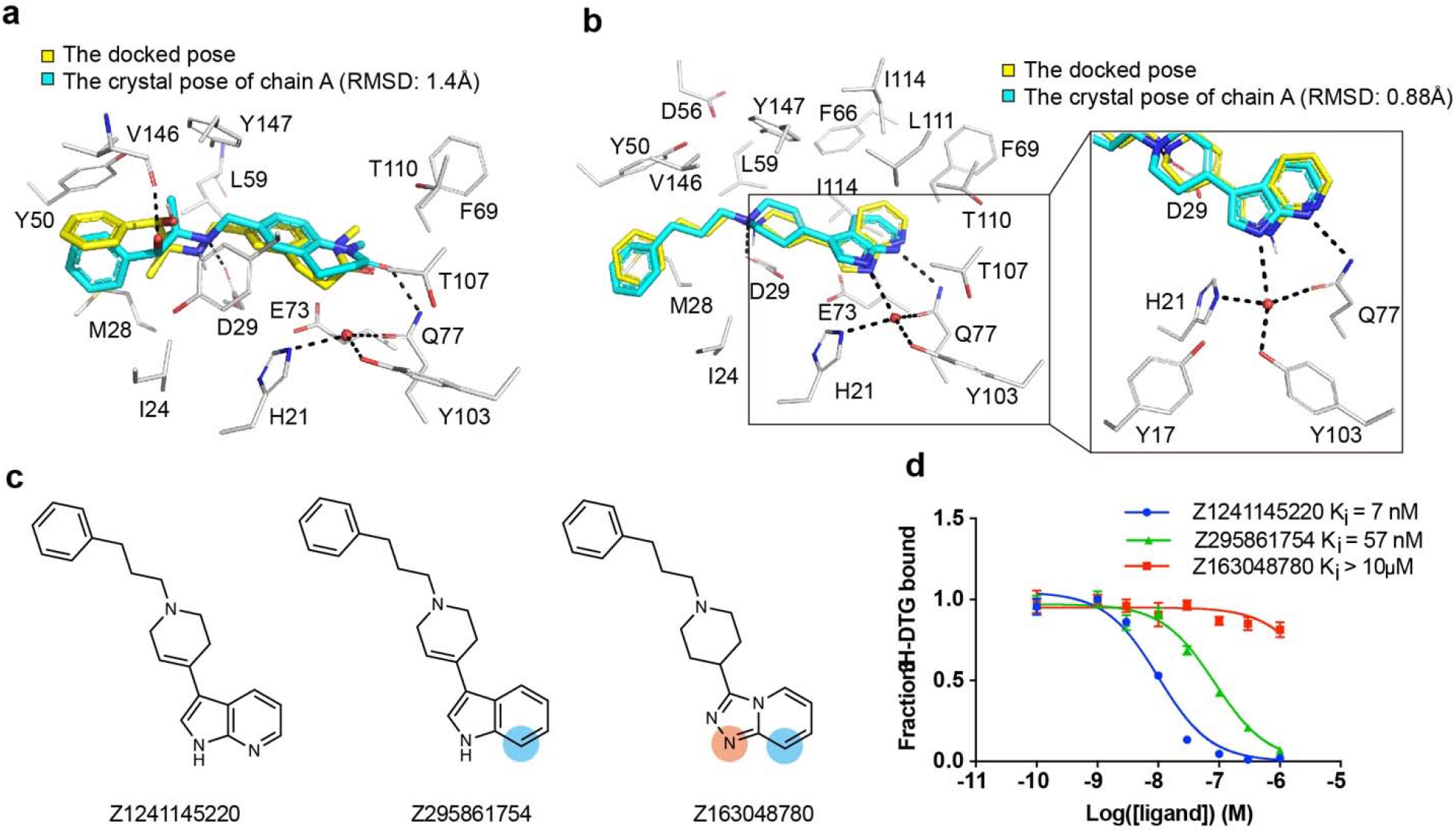
High structural fidelity between docked-predicted and crystallographically-determined poses of the new σ_2_ receptor ligands. Crystal structures of the ligands (carbons in cyan) are overlaid with their respective docking predictions (yellow). σ_2_ receptor carbon atoms are depicted in grey, oxygens in red, nitrogens in blue, sulfurs in yellow, Hydrogen bonds are shown as black dashed lines. **a**, The complex with Z4857158944 (PDB code: 7M96; RMSD = 1.4 Å). **b,** The complex with Z1241145220 (PDB code: 7M95; RMSD = 0.88 Å). **c,** Two analogues that disrupt the hydrogen bonds with Gln77 and the structural water. Differences between the analogues and the parent compound Z1241145220 are depicted with light blue and apricot circles. **d**, Competition binding curve of the three compounds show reduced affinity at σ_2_.

To investigate whether this ordered water is important for ligand recognition, we tested two analogues that were designed to disrupt the hydrogen bonds between Gln77 and the water (**Fig. 3c**). Z295861754, which is only expected to hydrogen-bond with the water but not with Gln77, showed an ~8-fold decrease in binding affinity whereas Z163048780, which is not expected to hydrogen bond with either Gln77 or the water, had a K_i_ value > 10 μM (**Fig. 3d**), indicating a crucial role of the water in the binding of Z1241145220 to the σ_2_ receptor. To generalize these findings further, and to test whether this water is a structural element of the binding site, we generated a series of σ_2_ mutants in which the coordination of this water molecule was disrupted. We measured the affinity of the radioligand probe [^3^H]-DTG to these mutants and then performed competition binding assays with Z1241145220 (**Supplementary Information Fig. 7**). These experiments showed that mutating either His21 or Gln77 reduces the affinity of Z1241145220 by an order of magnitude. Taken together, these results demonstrate that the ordered water is an integral part of the binding pocket and is required for high-affinity binding of Z1241145220, and potentially for other ligands as well. This is reminiscent of a structural water molecule required for ligand recognition at the μ-opioid receptor^40^.

### Novel σ_2_ ligands reduce mechanical hypersensitivity in a mouse model of neuropathic pain

There is strong genetic^41,42^ and pharmacological^43–45^ evidence supporting a contribution of σ_1_ to chronic pain^46^, but only recently, with the discovery of the gene encoding for σ_2_^4^, has it been fully possible to understand and distinguish the roles of σ_2_ and σ_1_ in these processes^21,22^. However, the limited availability of highly selective σ_2_ probes^22^ has hindered disentangling the contribution of σ_1_ and σ_2_ in these processes. To investigate this, we treated mice with three high-affinity σ_2_ ligands with differing degrees of selectivity: Z4857158944 (4 nM; >250-fold σ_2_/σ_1_ selectivity), Z1665845742 (5 nM; 47-fold σ_2_/σ_1_ selectivity), and Z4446724338, a 3 nM non-selective ligand (**Fig. 4a**). As a comparator, we also treated with the well-known reagent PB28, a 5 nM σ_1_/σ_2_ non-selective ligand^31^. In pharmacokinetics experiments, the three docking-derived ligands had substantial brain permeability, with brain to plasma ratios ranging from 3 to 16, and brain half-lives ranging from 1.2 to 12 hours (**Supplementary Information Table 4**). The non-selective PB28 also had high brain permeability and a relatively long half-life, though its brain C_max_ was three to eight-fold lower than that of the new docking-derived compounds. Even so, the high brain exposures of all four compounds encouraged us to examine them in a neuropathic pain model in mice.

**Figure 4.**
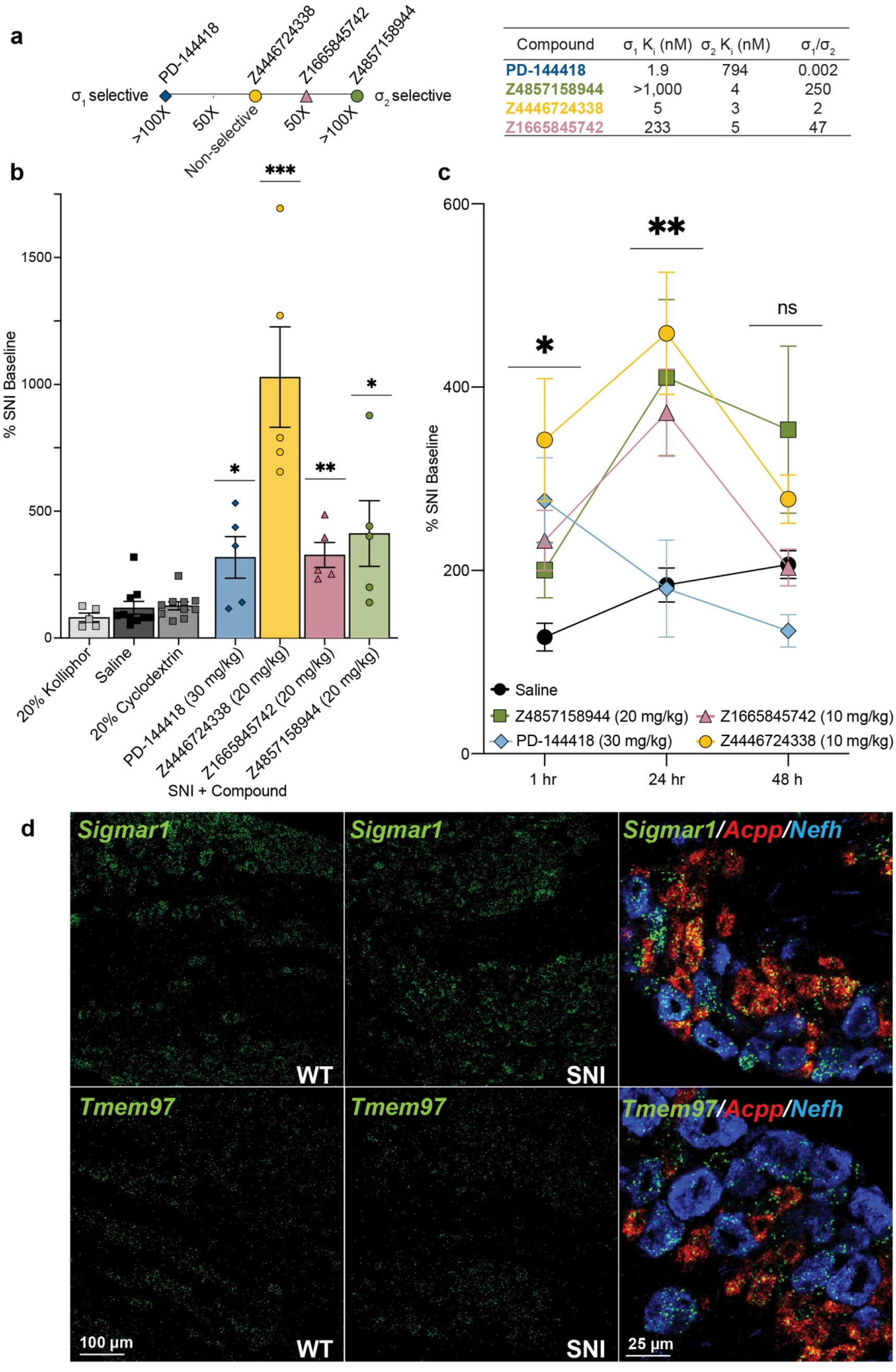
Systemic σ_1/2_ ligands are anti-allodynic in a model of neuropathic pain. **a**, Selectivity of five ligands at σ_1_ and σ_2_, based on their experimental K_i_ ratios. PD-144418 values taken from the literature^12^. **b**, Response of mice to a Von Frey filament after spared nerve injury (SNI). All four ligands reduce the SNI-induced mechanical hypersensitivity compared to their vehicle (PD-144418 vs kolliphor; Z4446724338 and Z4857158944 vs cyclodextrin; Z1665845742 vs saline; one-way ANOVAs; * p < 0.05, ** p < 0.01, *** *p* < 0.001). Data shown are mean ± SEM. Data replotted, and statistical testing results described in **Supplementary Information Fig. 8a**. **c**, The anti-allodynic effects of σ_2_, but not σ_1_, ligands increases over time, with a peak effect at 24 hours postinjection. Significance levels determined using Dunnett’s multiple comparisons Post-hoc test reflect the difference between Z4446724338 and saline for simplicity (two-way ANOVA; time x treatment interaction: *F*(8,80) = 2.25, *p* = 0.03; time: *F*(2,76) = 5.09, *p* = 0.009; treatment: *F*(4,40) = 5.40, *p* = 0.001; four treatment groups (*n* = 10) except PD-144418 (*n* = 5); ns = not significant, * p < 0.05, ** p < 0.01). Data shown are mean ± SEM. **d**, *in situ* hybridization of mouse dorsal root ganglion (DRG) sections for *Sigmar1* (σ_1_) and *Tmem97* (σ_2_) genes illustrates expression in myelinated (Nefh-positive; blue) and unmyelinated (Acpp-positive; red) subsets of sensory neurons and no change after SNI.

We tested the efficacy of these σ_1_ and σ_2_ ligands in the spared nerve injury (SNI) mouse model of neuropathic pain, in which two out of three branches of the sciatic nerve are transected^47^, resulting in significant mechanical hypersensitivity (allodynia) transmitted by the uninjured peripheral (sural) nerve. SNI mice systemically injected with either of the σ_2_-selective ligands Z1665845742 or Z4857158944 were strongly anti-allodynic when dosed in SNI mice, significantly increasing mechanical thresholds versus vehicle (**Fig. 4b**, **Supplementary Information Fig. 8a**). Intriguingly, anti-allodynic effect of Z1665845742 and Z4857158944 was comparable to that attained by a systemic injection of the σ_1_-selective ligand PD-144418. To investigate the possible synergistic effect of targeting both σ receptors we tested the nonselective ligands PB28 and Z4446724338. Systemic injection Z4446724338 dose-dependently increased the mechanical thresholds of SNI mice, 1-hour post-injection (**Fig 4b**, **Supplementary Information Fig. 8a**), with the highest dose completely reversing the SNI-induced mechanical allodynia (i.e., thresholds returned to pre-injury levels). In contrast, systemic injections of PB28, a well-established sigma receptor ligand with high affinity for both subtypes^48^, produced mixed results, with anti-allodynic effects observed only in 60% of the mice **(Supplementary Information Fig. 8a**). The much stronger anti-allodynia of Z4446724338 versus PB28 may reflect the former’s substantially higher brain permeability as measured by their respective brain C_max_ values (**Supplementary Information Table 4**). Importantly, none of the new σ_1_ and σ_2_ ligands were sedative on the rotarod test (**Supplementary Information Fig. 8b**), indicating that their anti-allodynic effect was not due to motor impairment.

On its face, substantial anti-allodynic effects of the σ_2_-selective ligands Z1665845742 and Z4857158944 suggest that this receptor is a potential target for managing neuropathic pain in the clinic. However, because σ_2_ ligands are notorious for activity at other targets, especially GPCRs^49,50^, we counter-screened all three docking-derived ligands against a panel of potential off-targets. In a TANGO screen^51^ of 320 GPCRs, the molecules did not act as agonists or inverse agonists against most targets. Some activity was observed at the 5HT1A receptor and κ-opioid receptors (**Supplementary Information Fig. 9**), but this activity did not replicate in subsequent concentration-response assays. Because a key pain target, the μ-opioid receptor, is often relatively inactive in TANGO assays, we further tested the molecules in G protein assays against this receptor; here too, no substantial activity was observed (**Supplementary Information Fig. 9**). Since the TANGO assay is limited to detecting agonism (and in some cases inverse-agonism) at GPCRs, we further screened the compounds in binding assays against a panel of 44 targets including GPCRs, ion channels, and transporters. No binding was observed for any pain-related targets (**Supplemental Information Table 5)**. Taken together, these observations suggest that the primary mechanism of action of the selective ligands is via the σ_2_ receptor (naturally, other targets, not tested here, cannot be excluded). The even stronger activity of the σ_1/2_ ligand Z4446724338 suggests that σ_1/2_ polypharmacology may further increase activity.

### σ_2_ ligands achieve peak antiallodynic effects 24 hours after dosing

In earlier studies, σ_1/2_ ligands showed peak antiallodynic effects up to 48 hours after dosing^21^. This unusual behavior was observed with ligands with mid-nanomolar potency for the σ_2_ receptor, and 9 to 14-fold selectivity vs. the σ_1_ receptor. We thought it interesting to further explore this with the low nanomolar affinity selective ligands, Z4857158944 and Z1665845742, and the joint σ_1/2_ ligand, Z4446724338. The three molecules were tested post SNI, at 1 hour, 24 hours, and 48 hours after dosing. Consistent with earlier reports, the anti-allodynic effects of the three novel σ ligands increased over time, reaching maximal effect 24 hours post-injection (**Fig. 4c**). In contrast, the anti-allodynic effect of the selective σ_1_ ligand PD-144418 was not sustained 24-or 48-hours post-injection. Furthermore, although the σ_2_-selective compounds exhibited reduced anti-allodynia efficacy at early time points compared to the mixed agonist Z4446724338, all three compounds produced similar levels of analgesia by 24 hours. We note that this long-term activity cannot be easily explained by pharmacokinetics, as the brain half-life of all three compounds suggests minimal exposure past 12 hours (**Supplementary Information Table 4**). Rather, this time course may reflect longer term signaling or regulatory effects, the exact nature of which remains a question for ongoing research^21^. Regardless of the basis, these results confirm earlier work suggesting that the antiallodynic effects of σ_2_ are prolonged, which may be useful in the management of chronic pain disorders.

### σ_1/2_ receptors are expressed on primary afferent neurons

*in situ* hybridization of dorsal root ganglia (DRG) sections, where the cell bodies of sensory neurons that transmit the “pain” message to the spinal cord reside, revealed that both σ_1_ and σ_2_ receptors are expressed in a wide variety of DRG neurons, including myelinated and unmyelinated subsets (**Fig. 4d**). We additionally found that the expression level of σ_1_ or σ_2_ did not change in DRG neurons 7 days after SNI. Although the sigma receptor has a broad distribution, we suggest that our new ligands exert their antinociceptive action via interaction with primary afferent neurons.

## Discussion

The σ_2_ receptor has been pharmacologically enigmatic for 30 years. Its implication in diverse biological processes and lack of molecular data has hindered progress in understanding its biological role. Four key observations from this study begin to illuminate these issues. **First**, high-resolution crystal structures of the σ_2_ receptor complexed with roluperidone and with PB28 reveal a ligand-binding site deeply embedded in the membrane (**Fig. 1a-b**), suggesting the possibility of a lipid as an endogenous ligand. The evolutionary connection of σ_2_ to EBP and the structure of the receptor bound to cholesterol strongly imply an ability to recognize sterols. The structures explain the simple pharmacophore of σ_2_ ligands—a positively charged amine that ion-pairs with Asp29, while flanking hydrophobic and aromatic moieties are recognized by nearby aromatic residues, such as Phe66, Phe69, His21 and Tyr50. The structures also highlight nearby polar residues, such as Gln77, and Thr110, that seem rarely exploited by classic σ_2_ ligands, but which may provide new selectivity determinants for ligand discovery (**Fig. 1c** and **1d**). **Second**, by testing 484 compounds across docking ranks from a library of 490 million molecules, a clear and quantitative relationship emerged between docking score and the likelihood of binding (**Fig. 2**). Encouragingly, crystal structures of docking-derived ligands confirmed the docking predictions with low RMSDs (**Fig. 3a** and **3b**). **Third**, from among the top-ranking docking hits were 31 novel scaffolds with potent binding affinities (K_i_ < 100 nM) (**Supplementary Information Table 2-3**). Optimization of two of these led to ligands with low nanomolar affinities and 47-fold to >250-fold selectivity for the σ_2_ over the σ_1_ receptor (**Supplementary Information Fig. 1**). **Fourth**, three new σ_2_ chemotypes, one non-selective but potent vs. σ_1_/σ_2_, and two others selective for σ_2_ over σ_1_, were tested for efficacy in a mouse model for neuropathic pain. All three showed antiallodynic effects (**Fig. 4**); the expression pattern of the receptor and the activity of the σ_2_-selective ligands confirm a contribution of this receptor in pain processing and suggest its potential relevance in pain management.

The σ_2_ and the σ_1_ receptors are promiscuous, both binding to cationic amphiphiles, and as expected, there is broad cross-reactivity between the two receptors. Although selective σ_1_ ligands, like PD-144418 and (+)-pentazocine, have been described, there are relatively few selective ligands^22,52^ for the σ_2_ receptor, which has hindered understanding its role in biology and in disease. We sought to optimize for selective ligands from among our potent docking actives. We adopted a chemical novelty approach described previously^29,53,54^ prioritizing novel scaffolds, modeled to interact with diverse groups within the σ_2_ structure. We reasoned that this would identify more potent molecules that would also be selective vs the σ_1_ receptor. This strategy has been productive previously, while the more rational approach of intentionally counter-docking vs. the off-target, here the σ_1_ receptor, remains a research area^55^. A cycle of structure-based analoging resulted in improved selectivity for two chemotypes. Increasing the linker length in ZINC450573233 by one carbon, along with two other small changes, results in Z1665845742, with selectivity improved from 30-fold to 47-fold. At the same time, replacing the 2,3-dihydro-1,4-benzodioxine group in ZINC895657866 with a 3,4-dihydro-1H-quinolin-2-one to engage with Gln77 (**Fig. 3a**), led to Z4857158944, with selectivity increased from 46-fold to >250-fold, making it among the most potent and selective σ_2_ ligands of which we are aware. We combined one of these selective molecules with a close analog that is inactive on the σ_2_ receptor, affording a “probe pair” (Z1665845742 and Z1665798906) (**Supplementary Information Fig. 10)**. Such probe pairs are useful to understand the role of receptors, such as the σ_2_ in cellular and *in vivo* studies, where the activity of the inactive member controls for off-targets that any molecule inevitably has. The approach should also facilitate disentangling the role of the σ_2_ receptor in indications for which it has been widely mooted, including cancer^9–11^, schizophrenia^17^, and Niemann Pick disease^5,6^. We make this probe pair openly available via Sigma-Millipore’s probe collection (Cat. No. TBD).

Certain caveats bear airing. While our ultimate goal was to find σ_2_-selective ligands from the docking, ligands with a spectrum of affinities and selectivities for both σ receptors emerged, reflecting the similarities of their pockets and the historical precedence in their overlapping pharmacology (**Fig. 4c-e**). Human inspection of molecules revealed an unusually high 66% hit rate, as well as competitive ligand potency, likely reflecting a site that is well-suited to ligand binding, despite its solvent-occlusion and hydrophobicity. These high hit rates, potencies, and selectivities often do not translate to other targets—this target was unusually propitious for library docking. Additionally, while the docking-predicted pose for Z4857158944 and for Z1241145220 closely resembled the subsequent crystallographic structure, important differences remain, namely the water-bridging interaction for Z1241145220. Although modeling this water was not necessary for ligand discovery, and the docked ligands fit well without it, including this water modeling may further improve future structure-based efforts against this target. Modeling waters in docking remains a research area in molecular docking^56^.

The key observations of this work should not be obscured by these caveats. The high-resolution crystal structures of σ_2_ receptors reveal the origins of its molecular recognition, and template structure-based campaigns for novel ligand discovery. This work emphasizes the value of a structure-based approach to screen vast new libraries of chemotypes unrelated to known ligands with unique properties that illuminate the biology of the σ_2_ receptor. Applications to other targets should undoubtedly be considered.

## Methods

### Protein expression and purification for crystallography

The bovine σ_2_ receptor was cloned into pVL1392 with an N-terminal human protein C epitope tag followed by a 3C protease cleavage site. The construct was truncated after residue 168 to exclude the ER localization signal for better expression and to facilitate crystallization. This receptor construct was expressed in *Sf9* insect cells (Expression Systems) using the BestBac baculovirus system (Expression Systems) according to manufacturer’s instruction. Infection was performed when cell density reached 4×10^6^ cells per milliliter. Cells were shaken at 27 °C for 60 hours before harvest by centrifugation. Cell pellets were stored at −80 °C until purification.

During all purification steps ligands (PB28, roluperidone, Z1241145220, and 7866num 1973.2) were present in all buffers at 1 μM. For the cholesterol-bound structure the protein was purified in the presence of 1 μM DTG. Cell paste was thawed and cells were disrupted by osmotic shock in 20 mM HEPES pH 8, 2 mM magnesium chloride, 1:100,000 (v:v) benzonase nuclease (Sigma Aldrich), and cOmplete EDTA-free Protease Inhibitor Cocktail (Roche). Lysed cells were centrifuged at 50,000 x g for 15 minutes. Following centrifugation, supernatant was discarded and the membrane pellets were solubilized with a glass dounce tissue homogenizer in 20 mM HEPES pH 8, 250 mM NaCl, 10% (v/v) glycerol, 1% (w/v) lauryl maltose neopentyl glycol (LMNG; Anatrace), and 0.1% (w/v) cholesterol hemisuccinate (CHS; Steraloids). Samples were stirred at 4 °C for 2 hours and then nonsolubilized material was removed by centrifugation at 50,000 x g for 30 min. Supernatant was supplemented with 2 mM calcium chloride and filtered by a glass microfiber filter (VWR). Samples were then loaded by gravity flow onto 5 ml anti-protein C antibody affinity resin. Resin was washed with 10 column volumes of 20 mM HEPES pH 8, 250 mM NaCl, 2 mM calcium chloride, 1% (v/v) glycerol, 0.1% (w/v) lauryl maltose neopentyl glycol, and 0.01% (w/v) cholesterol hemisuccinate, and then with 10 column volumes of 20 mM HEPES pH 8, 250 mM NaCl, 2 mM calcium chloride, 0.1% (v/v) glycerol, 0.01% (w/v) lauryl maltose neopentyl glycol, and 0.001% (w/v) cholesterol hemisuccinate. The receptor was eluted with buffer containing 20 mM HEPES pH 8, 250 mM NaCl, 5 mM EDTA, 0.1% (v/v) glycerol, 0.01% (w/v) lauryl maltose neopentyl glycol, 0.001% (w/v) cholesterol hemisuccinate, and 0.2 mg/ml protein C peptide in 1 ml fractions. Peak fractions were pulled and 3C protease was added (1:100 w:w) and incubated with the receptor at 4 °C overnight. Next the receptor was purified by size exclusion chromatography on a Sephadex S200 column (GE Healthcare) in 20 mM HEPES pH 8, 250 mM NaCl, 0.1% glycerol, 0.01% lauryl maltose neopentyl glycol, and 0.001% cholesterol hemisuccinate. Peak fractions were pulled, calcium chloride was added to 2 mM and the sample was reapplied on the anti-protein C resin to remove uncleaved receptor. The column was washed with 5 column volumes and flow-through and wash fractions were pulled, concentrated, and reapplied on SEC. Peak fractions were pulled, concentrated to 50 mg/ml, and aliquoted. Protein aliquots were flash frozen in liquid nitrogen and stored in −80 °C until use. Purity was evaluated by SDS-PAGE.

### Crystallography and data collection

Purified σ_2_ receptor was reconstituted into lipidic cubic phase (LCP) by mixing with a 10:1 (w:w) mix of monoolein (Hampton Research) with cholesterol (Sigma Aldrich) at a ratio of 1.5:1.0 lipid:protein by mass, using the coupled syringe reconstitution method^57^. All samples were mixed at least 100 times. The resulting phase was dispensed in 30–40 nl drops onto a hanging drop cover and overlaid with 800 nl of precipitant solution using a Gryphon LCP robot (Art Robbins Instruments). The PB28-bound crystals grew in 20–30% PEG 300, 0.1 M MES pH 6, 600 mM NaCl. The Roluperidone-bound crystals grew in 20% PEG 300, 0.1 M MES pH 6, 500 mM NaCl, 60 mM succinate. The Z1241145220-bound crystals grew in 30% PEG 300, 0.1 M MES pH 6, 210 mM ammonium phosphate. The 7866num 1973.2-bound crystals grew in 30% PEG 300, 0.1 M MES pH 6, 560 mM ammonium phosphate. The cholesterol-bound crystals grew in 25% PEG300, 0.1 M MES pH 6, 400 mM sodium citrate, and 1% 1,2,3-heptanetriol. All crystals grew in the presence of 1 μM of ligand, except for the cholesterol structure, which had no ligand present during crystal growth. Crystals were harvested using either MicroLoops LD or mesh loops (MiTeGen) and stored in liquid nitrogen until data collection. Data collection was performed at Advanced Photon Source GM/CA beamlines 23ID-B and 23ID-D. Data collection used a 10 μm beam and diffraction images were collected in 0.2° oscillations at a wavelength of 1.254858 Å for the PB28-bound crystals and a wavelength of 1.033167 Å for all other crystals. A complete data set was obtained from a single crystal in each case.

### Data reduction and refinement

Diffraction data were processed in HKL2000^58^ and in XDS^59^, and statistics are summarized in **Supplementary Information Table 1**. The PB28-bound structure was solved using molecular replacement starting with a Rosetta homology model generated using the structure of EBP (Protein Data Bank accession 6OHT). Matthews probability predicted four copies in the asymmetric unit. Initially, a single copy of this model was placed using Phaser. This model did not fit well into density and was replaced with Idealized helices that were used as a search model for a search for an additional copy. The resulting dimer was duplicated and manually placed into unmodeled density. The resulting structure was iteratively refined in Phenix^60^ and manually rebuilt in Coot^61^. Final refinement statistics are summarized in **Supplementary Information Table 1**. The PB28 structure was used as a model for molecular replacement for all other datasets. In the case of the structure modeled as cholesterol-bound, electron density for a sterol-shaped ligand was observed and tentatively modeled as cholesterol based on the high (millimolar) concentration of cholesterol in the crystallization conditions and the compatibility of cholesterol with the shape of the electron density in the binding pocket. The receptor was purified in the presence of ditolylguanidine (DTG), but no DTG was present in the precipitating solution, and electron density was clearly incompatible with bound DTG. We cannot exclude the possibility that some other compound structurally similar to cholesterol was carried through the purification and is the ligand observed in the binding pocket.

### Preparation of membranes for radioligand binding

The human σ_2_ receptor was cloned into pcDNA3.1 (Invitrogen) mammalian expression vector with an amino-terminal protein C tag followed with a 3C protease cleavage site. Mutations were introduced by Site-directed mutagenesis using HiFi HotStart DNA Polymerase (Kapa Biosystems). Expi293 cells were transfected using FectoPRO (Polyplus-transfection) according to manufacturer instruction. Cells were harvested by centrifugation and lysed by osmotic shock in a buffer containing 20 mM HEPES, pH 7.5, 2 mM MgCl2,1:100,000 (vol/vol) benzonase nuclease (Sigma Aldrich), and cOmplete Mini EDTA-free protease-inhibitor tablets (Sigma Aldrich). The lysates were homogenized with a glass Dounce tissue homogenizer and then centrifuged at 20,000g for 20 min. After centrifugation, the membranes were resuspended in 50 mM Tris, pH 8.0, divided into 100 μl aliquots, flash frozen in liquid nitrogen, and stored at −80 °C until use.

### Saturation and competition binding in Expi293 membranes

Saturation binding was performed with a method similar to that of Chu and Ruoho^62^. Briefly, membrane samples from Expi293 cells expressing wild-type or mutant σ_2_ receptor, prepared as described above, were thawed, homogenized with a glass Dounce, and diluted in 50 mM Tris, pH 8.0. Binding reactions were done in 100 μL, with 50 mM Tris pH 8.0, [^3^H]-DTG (PerkinElmer), and supplemented with 0.1% bovine serum albumin to minimize non-specific binding. To assay non-specific binding, equivalent reactions containing 10 μM haloperidol were performed in parallel. Competition assay were performed in a similar fashion with 10 nM [^3^H]-DTG and the indicated concentration of the competing ligand. Samples were shaken at 37 °C for 90 min. Afterward, the reaction was terminated by massive dilution and filtration over a glass microfiber filter with a Brandel harvester. Filters were soaked with 0.3% polyethyleneimine for at least 30 min before use. Radioactivity was measured by liquid scintillation counting. Data analysis used GraphPad Prism, with K_i_ values calculated by Cheng-Prusoff correction using the experimentally measured probe dissociation constant.

### Circular dichroism

Far-UV circular dichroism (CD) spectra (185–260 nm) were measured with a JASCO J-815 (JASCO Inc., Tokyo, Japan) using 1 mm path length cuvettes. Protein concentration was 0.5 mg/ml in 10 mM potassium phosphate pH 7.4, 250 mM potassium fluoride. Melt curves were measured at 222 nm between temperatures 20-80 °C.

### Size-exclusion chromatography with multi-angle light scattering (SEC-MALS)

The oligomeric state of σ_2_ receptor was assessed by SEC–MALS using a Wyatt Dawn Heleos II multi-angle light scattering detector and Optilab TrEX refractive index monitor with an Agilent isocratic HPLC system Infinity II 1260. Receptor was prepared as described above, but with no ligand added during purification. The ligand-free receptor was diluted to 1□mg□ml^-1^ in SEC-MALS buffer (0.01% lauryl maltose neopentyl glycol, 20 mM HEPES pH 7.5, 150□mM sodium chloride). Ligands were added to a final concentration of 1□μM and the sample was incubated with ligand for 2□h at room temperature (21□°C). Separation steps were performed in SEC-MALS buffer with a Tosoh G4SWxl column at a flow rate of 0.5□ml□min^-1^. Data analysis used the Astra software package version 6.1.4.25 (Wyatt) using the protein conjugate method with a dn/dc value of 0.21 (mL/g) for detergent and 0.185 (mL/g) for protein.

### Molecular docking

The σ_2_ receptor bound to cholesterol (PDB ID 7MFI) was used in the docking calculations. The structure was protonated at pH 7.0 by Epik and PROPKA in Maestro^63^ (2019 release). Based on the mutagenesis data^4^, E73 was modeled as a neutral residue. AMBER united atom charges were assigned to the structure. To model more realistic low protein dielectric boundary of this site, we embedded the receptor into a lipid-bilayer to capture its native environment in endoplasmic reticulum (ER) membrane, then followed by a 50 ns coarse-grained molecular dynamic (MD) simulation with a restricted receptor conformation. A more detailed protocol can be found on the DISI wiki page (http://wiki.docking.org/index.php/Membrane_Modeling). The volume of the low dielectric and the desolvation volume was extended out 2.2 Å and 1.2 Å, respectively, from the surface of protein and modelled lipid-bilayer using spheres calculated by SPHGEN. Energy grids were pre-generated using CHEMGRID for AMBER van der Waals potential^64^, QNIFFT^65^ for Poisson–Boltzmann-based electrostatic potentials, and SOLVMAP^66^ for ligand desolvation.

The resulting docking setup was evaluated for its ability to enrich known σ_2_ ligands over property-matched decoys. Decoys are unlikely to bind to the receptor because despite their similar physical properties to known ligands, they are topologically dissimilar. We extracted 10 known σ_2_ ligands from ChEMBL(https://www.ebi.ac.uk/chembl/) including PB28 and roluperidone whose crystallographic poses were report here. Five-hundred and forty-two property-matched decoys were generated by the DUDE-Z pipeline^67^.

Docking performance was evaluated based on the ability to enrich the knowns over the decoys by docking rank, using log adjusted AUC values (logAUC). The docking setup described above is able to achieve a high logAUC of 39 and to recover the crystal poses of PB28 and roluperidone with RMSD values of 0.93 and 0.77 Å, respectively. We also constructed an ‘extrema’ set (cite the protocol paper) of 61,687 molecules using the DUDE-Z web server (http://tldr.docking.org) to ensure that molecules with extreme physical properties were not enriched. The docking setup enriched close to 90% mono-cations among the top1000 ranking molecules. In order to check if the limited amounts of knowns and property-matched decoys over-trained the docking parameters, the enrichment test was run using 574 additional σ_2_ ligands from S2RSLDB^52^ (http://www.researchdsf.unict.it/S2RSLDB) against the ‘extrema’ set. The resulting high logAUC of 41 demonstrated the docking setup was still able to enrich knowns over decoys on a 112-fold larger test set, indicating the favorable docking parameters for launching an ultra-large-scale docking campaign.

Four-hundred and ninety million cations from ZINC15 (http://zinc20.docking.org), characterized by similar physical properties as σ_1/2_ known ligands (for instance, with calculated octanol-water partition coefficients (cLogP) <=5 and with 250 Da <molecular weight <=400 Da), was then docked against the σ_2_ ligand binding site using DOCK3.8. Of these, 469 million molecules were successfully docked. On average, 3,502 orientations were explored and for each orientation, 183 conformations were averagely sampled. In total, more than 314 trillion complexes were sampled and scored. The total calculation time was 177,087 hours, or 3.7 calendar days on a cluster of 2,000 cores.

The top-ranking 300,000 molecules were filtered for novelty using the ECFP4-based Tanimoto coefficient against 2,232 σ_1/2_ ligands in ChEMBL (https://www.ebi.ac.uk/chembl/) and 574 σ_2_ ligands from S2RSLDB (http://www.researchdsf.unict.it/S2RSLDB). Molecule with Tanimoto coefficients of (T_c_) ≥ 0.35 were eliminated. The remaining 196,170 molecules were clustered by ECFP4-based Tc of 0.5, resulting in 33,585 unique clusters. From the top 5,000 novel chemotypes, molecules with > 2 kcal/mol internal strains were filtered out using strain_rescore.py in Macromodel (2019 release). After filtering for novelty and diversity, the docked poses of the best-scoring members of each chemotypes were manually inspected for favorable and diversified interactions with the σ_2_ site, such as the salt bridge with Asp29, the hydrogen bond with His21/Val146 and the π-π stacking with Tyr50/Trp49. Ultimately, 86 compounds were chosen for testing, 79 of which were successfully synthesized.

### Hit-rate curve prediction

In order to guide the design of scoring bins for the hit rate curve, 1,000 docked poses were sampled every 2.5 kcal/mol from the best score of −65 kcal/mol up to −22.5 kcal/mol. The estimated hit rate was calculated by the number of sensible docked poses divided by 1,000. The criteria to define a sensible docked pose contains 1) no unsatisfied hydrogen bond donors; 2) less than 3 unsatisfied hydrogen acceptors; 3) forms a salt bridge with Asp29; 4) total torsion strain energy < 8 units; 5) maximum strain energy per torsion angle < 3 units. The first three filters were implemented based on LUNA (https://github.com/keiserlab/LUNA), which calculated all the intra- and interactions of a docked pose with the receptor, then hashed them into a binary fingerprint. The strain energy was calculated by an in-house population-based method^68^. Based on the shape of the estimated prior curve (**Supplementary Information Fig. 11**), more scoring bins are selected in the higher estimated hit-rate region: −65, −59.73 and −57.5 kcal/mol. After that, every scoring bin was 2.5 kcal/mol from each other till −37.5. The last four bins were 5 kcal/mol from each other. 13,000 molecules sampled were from these 14 scoring bins were filtered by novelty and internal torsion strain described above. The remaining 9,216 novel and none-strained molecules were cluster by the LUNA 1024-length binary fingerprint of a Tc = 0.32, resulting in 6,681 clusters. The first 40 chemotypes were attempted to be purchased from each scoring bin. After the evaluation of synthesis availability from the vendors, 491 molecules were ordered.

### Hit-rate curve fitting

To fit the Bayesian hit-rate models we used Stan^69^ (v2.21.2) via BRMS^70^ (v2.14.4), with generic parameters: iter=4000, and cores=4. Here are the model specific parameters. For both hit-picking prior and posterior Sigmoid models formula=bmrs::formula(hit ~ top * inv_logit(hill*4/top*(dock_energy - dock50)), top + hill + dock50 ~ 1, nl=TRUE), where hill is scaled by 4/top so it is the slope of the curve at the dock50 irrespective of the value of Top. For Prior Sigmoid model, prior=c(brms::prior(normal(.5, .2), lb=0, ub=1, nlpar=“top”), brms::prior(normal(−50, 10), nlpar=“dock50”), brms::prior(normal(−.1, .1), ub=-.001, nlpar=“hill”)), inits=function(){list(top=as.array(.5), dock50=as.array(−50), hill=as.array(−.1))}, family=gaussian(). Updating the Prior sigmoid model with the mean expected hit-rate for each computationally analyzed tranche yielded an estimate and 95% credible interval for the sigma parameter for the Gaussian response of 20 [15, 30]%, but did not significantly adjust the distributions for Top, Hill, or Dock50 (**Supplementary Information Fig. 12**). Therefore, to estimate the posterior sigmoid model, we transferred the per-parameter prior distributions and initial values and used the family=bernoulli(“identity”). To compare models we used the loo package to add the Pareto smoothed importance sampling leave-one-out (PSIS-LOO) and Bayesian version of the R2^71^ (loo_R2) information criteria. Figures were generated using tidybayes^72^, ggplot2^73^, and tidyverse^74^ packages in R^75^.

### Analoging within the make-on-demand library

Using 4 primary docking hits (ZINC450573233, ZINC533478938, ZINC548355486 and ZINC895657866) as queries in SmalWorld (https://sw.docking.org/) from the 28B make-on-demand library, a subset of Enamine REAL space, 20,005 analogues were selected by its default settings, then docked into the σ_2_ site for potential favorable interactions with His21, Tyr50, Gln77 and Val146.

### Make-on-demand synthesis

79 molecules that were prioritized from the initial cation-only docking screen were delivered within 7 weeks with a 93% fulfilment rate, and 412 molecules for the hit-rate curve were delivered within 4 weeks with an 82% fulfilment rate after a single synthesis attempt. Most of the make-on-demand molecules were derived from Enamine REAL database (https://enamine.net/compound-collections/real-compounds). For purity data on all compounds synthesized see **Supplementary Information Tables 6-8**.

### Yeast isomerase complementation assay

The human σ_2_ receptor, ERG2, and EBP were subcloned into the URA3 shuttle vector p416GPD. The plasmids were transformed into the Erg2-deficient *Saccharomyces cerevisiae* strain Y17700 (BY4742; MATα; ura3Δ0; leu2Δ0; his3Δ1; lys2Δ0; YMR202w::kanMX4) (Euroscarf) by the lithium acetate/single-stranded carrier DNA/polyethylene glycol method. A single colony was picked from a URA-selective plate and grown in suspension. Yeast were diluted in sterile water in a five-fold serial dilution starting from O.D. 0.1. Two microliters of the yeast dilutions were spotted on a URA-selective plate either in either the absence or the presence of sub-inhibitory concentrations of cycloheximide (50 ng/ml) and grown at 30°C for 24-48 h before imaging.

### Sterol isomerization enzymatic assay

EBP and σ_2_ were cloned into pcDNA3.1 (Invitrogen) mammalian expression vector with FLAG and protein C affinity tag, respectively. Proteins were purified as described for crystallography preparations, except no ligand was present during purification. Following size exclusion chromatography proteins were flash frozen in liquid nitrogen and kept at −80 °C until use. Zymostenol (CAS #566-97-2) and lathosterol (CAS #80-99-9) were purchased from Avanti Polar Lipids. For each sterol, a 2x solution was prepared by first dissolving DDM in isopropanol to 1% (w/v) and dissolving sterols in chloroform to a concentration of 1 mg/ml, followed by transferring 500 μM of the sterols to a new vial, evaporating under argon, and dissolving with DDM in a 1:20 (w/w) detergent to sterol ratio and a final 0.2% detergent in HEPES buffered saline (HBS; 20 mM HEPES pH 7.5, 150 mM NaCl). Proteins were diluted in HBS to 5 μM. Individual sterol standards were prepared by mixing each sterol 1:1 with HBS. A mixed sterol standard was prepared by mixing both sterols in a 1:1 ratio. For the enzymatic reactions, sterols were mixed in 1:1 ratio with the protein sample to give a final protein concentration of 2.5 μM, sterol concentration of 250 μM, and detergent concentration of 0.1%, in HBS. Reactions were incubated for 1 hour at 37 °C and then diluted 1:10 in methanol and kept at −20 °C until analysis by LC-MS. Samples were analyzed on a QE-plus mass spectrometer coupled to an Ultimate 3000 LC (Thermo fisher) in a method modified from Skubic *et al*.^76^. Five microliters were injected on a Force PFPP column coupled with an Allure PFPP column (both 2mm x 150 mm, Restek) maintained at 40°C. The mobile phases were A: methanol: isopropyl alcohol:water:formic acid (80:10:10:0.02) 5 mM ammonium formate, and B: isopropyl alcohol. The gradient was as follows: 0% B for 15 min, then 100% B in 1 second, maintained at 100% B for 5 min, followed by 5 min re-equilibration at 0% B. The flow rate was 0.15 mL min^-1^. The mass spectrometer was acquiring in t-SIM mode for the [M-H2O+H]+ ion (369.35158) with 70000 resolution, and 0.5 m/z isolation. Standard samples for each compound were run first separately to obtain the retention time of each of the two isobaric compounds.

### μOR activation assay

To measure μOR G_i/o_-mediated cAMP inhibition, 2.5 million HEK-293T cells were seeded in 10-cm plates. Eighteen to 24 hours later, upon reaching 85-90% confluency, cells were transfected using a 1:3 ratio of human μOR and a split-luciferase based cAMP biosensor (pGloSensorTM-22F; Promega). Transit 2020 (Mirus Biosciences) was used to complex the DNA at a ratio of 3□μL TransIT per μg DNA, in OptiMEM (Gibco-ThermoFisher) at a concentration of 10□ng DNA per μL OptiMEM. Twenty-four hours later, cells were harvested from the plate using Versene (PB□+□0.5□mM EDTA, pH□7.4) and plated in poly-D-lysine-coated white, clear-bottom 96-well assay plates (Corning Costar #3917) at a density of 35,000 cells per well and incubated at 37□°C with 5% CO_2_ overnight. The next day, after aspiration of the culture medium, cells were incubated for 2 hours covered, at room temperature, with 40 μL assay buffer (CO_2_-independent medium, 10% FBS) supplemented with 2% (v/v) GlosensorTM reagent (Promega). To stimulate endogenous cAMP via β adrenergic-G_s_ activation, 5x drugs were prepared in 10x isoproterenol containing assay buffer (200 nM final concentration). For naloxone competition experiments, 5x naloxone (1 μM final concentration) was also added to each well. Luminescence was immediately quantified using a BMG Clariostar microplate reader. Data were analyzed using nonlinear regression in GraphPad Prism 9.0 (Graphpad Software Inc., San Diego, CA).

### Off-target counterscreens

Screening of compounds in the PRESTO-Tango GPCRome was accomplished as previously described^51^ with several modifications. First, HTLA cells were plated in DMEM with 10% FBS and 10 U ml-1 penicillin–streptomycin. Next, the cells were transfected using an in-plate PEI method^77^. PRESTO-Tango receptor DNAs were resuspended in OptiMEM and hybridized with PEI before dilution and distribution into 384-well plates and subsequent addition to cells. After overnight incubation, drugs were added to cells without replacement of the medium. The remaining steps of the PRESTO-Tango protocol were followed as previously described. For those six receptors for which activity was reduced to less than 0.5-fold of basal levels of relative luminescence units or for the one receptor for which basal signalling was increased greater than threefold of basal levels, assays were repeated as a full dose–response assay. Activity for none of the seven could be confirmed, and we discount the apparent activity seen in the single-point assay.

Radioligand binding screen of off-targets was performed by the National Institutes of Mental Health Psychoactive Drug Screen Program (PDSP)^78^. Detailed experimental protocols are available on the NIMH PDSP website at https://pdsp.unc.edu/pdspweb/content/PDSP%20Protocols%20II%202013-03-28.pdf.

### Animals

Animal experiments were approved by the UCSF Institutional Animal Care and Use Committee and were conducted in accordance with the NIH Guide for the Care and Use of Laboratory animals. Adult (8-10 weeks old) male C56BL/6 mice (strain #664) were purchased from the Jackson Laboratory. Mice were housed in cages on a standard 12:12 hour light/dark cycle with food and water *ad libitum*.

### Compounds

All ligands were synthesized by Enamine (https://enamine.net/) and dissolved 30 minutes prior testing. PB28 and Z1665845742 were resuspended in NaCl 0.9%. Z4857158944 and Z4446724338 were resuspended in 20% cyclodextran. PD-144418 was resuspended in 20% Kolliphor.

### Behavioral analyses

For all behavioral tests, animals were first habituated for 1 hour in Plexiglas cylinders. The experimenter was always blind to treatment. All tests were conducted 30 minutes after subcutaneous injection of the compounds. Hindpaw mechanical thresholds were determined with von Frey filaments using the updown method^79^. For the ambulatory (rotarod) test, mice were first trained on an accelerating rotating rod, 3 times for 5 min, before testing with any compound.

### Spared-nerve injury (SNI) model of neuropathic pain

Under isoflurane anesthesia, two of the three branches of the sciatic nerve were ligated and transected distally, leaving the sural nerve intact. Behavior was tested 7 to 14 days after injury and *in situ* hybridization was performed one week post-injury.

### *In situ* hybridization

*In situ* hybridization was performed using fresh DRG tissue from adult mice (8-10 week old), following Advanced Cell Diagnostics’ protocol and as previously described^80^. All images were taken on an LSM 700 confocal microscope (Zeiss) and acquired with ZEN 2010 (Zeiss). Adjustment of brightness/contrast and changing of artificial colors (LUT) were done with Photoshop. The same imaging parameters and adjustments were used for all images within an experiment.

### Statistical analyses

All statistical analyses were performed with GraphPad Prism 8.0 (GraphPad Software Inc., San Diego, CA) unless otherwise noted. All data are reported as means ± SEM unless otherwise noted. Dose-response experiments were analyzed with one-way ANOVA and time-course experiments were analyzed with two-way ANOVA, and both experiments used Dunnett’s multiple comparison post-hoc test to determine differences between specific treatments and vehicle controls visualized in the figures. Rotarod experiments were analyzed using one-way ANOVA (saline, Z1665845742, and Z4857158944) or unpaired two-tailed Student’s t-test (kolliphor and Z4446724338).

Details of analyses, including number of tested animals and groups, degrees of freedom, and *p*-values can be found in the figure legends.

## Code availability

DOCK3.7 is freely available for non-commercial research http://dock.compbio.ucsf.edu/DOCK3.7/. A web-based version is available at http://blaster.docking.org/.

## Reporting Summary

Further information on research design is available in the Nature Research Reporting Summary linked to this article.

## Data availability

The coordinates and structure factors for PB28-bound σ_2_, roluperidone-bound σ_2_, Z1241145220-bound σ_2_, 7866num1973.2-bound σ_2_, and cholesterol-bound σ_2_ have been deposited in the PDB with accession codes 7M93, 7M94, 7M95, 7M96, and 7MFI respectively. The identities of the compounds docked in this study are freely available from the ZINC database (http://zinc15.docking.org) and active compounds may be purchased from Enamine. Probe pairs (two similar ligands with and without activity) of σ_2_ are available by arrangement with Millipore Sigma (Cat. No. TBD). Any other data relating to this study are available from the corresponding authors on reasonable request.

## Ethical compliance

All animal experiments were approved by the Institutional Animal Care and Use Committee at UCSF and were conducted in accordance with the NIH Guide for the Care and Use of Laboratory animals.

## Acknowledgements

Funding to support this research was provided by NIH grant R01GM119185, The Vallee Foundation, and the Sanofi iAwards program (ACK), by DARPA grant HR0011-19-2-0020 and NIH grant R35GM122481 (BKS). GM/CA@APS has been funded by the National Cancer Institute (ACB-12002) and the National Institute of General Medical Sciences (AGM-12006, P30GM138396). This research used resources of the Advanced Photon Source, a U.S. Department of Energy (DOE) Office of Science User Facility operated for the DOE Office of Science by Argonne National Laboratory under Contract No. DE-AC02-06CH11357. The Eiger 16M detector at GM/CA-XSD was funded by NIH grant S10 OD012289. We thank Dr. Kelly Arnett and the Harvard Center for Macromolecular Interactions for excellent support of biophysical experiments including circular dichroism and SEC-MALS. We would like to thank Charles Vidoudez and The Harvard Center for Mass Spectrometry for performing mass spectrometry analysis of sterols.

## Author contributions

A. A. performed cloning, mutagenesis, protein purification, SEC-MALS experiments, CD measurements, crystallization, X-ray data collection and processing, structure determination and refinement, radioligand binding, yeast complementation experiments, sterol isomerization enzymatic assay. J.L. conducted the docking, chemoinformatics analyses, docking energy analysis, and ligand picking, assisted in the latter by T.A.T. and B.K.S. J.M.B. conducted and analyzed the mouse allodynia experiments assisted by V.C., as well as the receptor expression experiments, supervised and co-analyzed by A.I.B. M.J.O. conducted the Bayesian analysis of docking scores vs. hit rates; C.M.W. tested molecules for activity against the μOR. X.P.H. and Y.L. tested compounds against the GPCR-ome and other off-targets, with supervision from B.L.R. Y.S.M. supervised the synthesis of molecules from the virtual library, J.J.I. was responsible for the building of the version of the ZINC library that was docked. A.C.K., B. K.S, and A.I.B. supervised the project. The manuscript was written by A.A., J.L., B.K.S, and A.C.K. with input from other authors.

