## Supplementary Information for "Crystal structures of the σ_2_ receptor template large-library docking for selective chemotypes active *in vivo*"

**Supplementary Information Table 1**. **Data collection and refinement statistics**

|  | PB28-bound  (Se-labeled) | Roluperidone-bound  (Native) | Z1241145220-bound  (Native) | 7866num1973.2-bound  (Native) | Cholesterol-bound  (Native) |
| --- | --- | --- | --- | --- | --- |
| **Data collection** |  |  |  |  |  |
| Space group | *P*2_1_ | *P*2_1_ | *P*2_1_2_1_2_1_ | *P*2_1_ | *P*2_1_ |
| Number of crystals | 1 | 1 | 1 | 1 | 1 |
| Cell dimensions |  |  |  |  |  |
| *a*, *b*, *c* (Å) | 70.6, 55.2, 93.0 | 69.1, 54.2, 99.7 | 55.4, 61.5, 110.4 | 70.7, 55.4, 93.0 | 70.6, 54.8, 92.7 |
| α, β, γ (°) | 90, 95.0, 90 | 90, 91.1, 90 | 90, 90, 90 | 90, 94.5, 90 | 90, 94.4, 90 |
| Wavelength (Å) | 1.255 | 1.03320 | 1.03321 | 1.033167 | 1.03320 |
| Resolution (Å) | 33.88 - 2.942  (3.047 - 2.942) | 42.61 - 2.71  (2.81 - 2.71) | 49.5 - 2.41  (2.55 - 2.41) | 40.2 - 2.41  (2.55 - 2.41) | 43.25 - 2.6  (2.64 - 2.6) |
| *R*_sym_ | 24.75 (88.16) | 26.11 (205.9) | 18.4 (177.4) | 19.67 (227.6) | 21.2 (51.0) |
| *<I>* / <σ*I>* | 5.73 (0.93) | 5.90 (0.71) | 7.50 (0.7) | 5.18 (0.56) | 3.625 (0.875) |
| Completeness (%) | 98.67 (90.76) | 99.54 (99.87) | 99.55 (99.36) | 97.9 (88.3) | 92.0 (51.9) |
| Redundancy | 4.0 (3.4) | 6.8 (6.5) | 6.2 (4.4) | 4.4 (4.5) | 3.2 (1.9) |
| CC_1/2_ | 98.7 (49.5) | 99.4 (36.6) | 99.6 (26.9) | 99.5 (28.3) | 96.2 (17.8) |
| **Refinement** |  |  |  |  |  |
| Resolution (Å) | 2.94 | 2.71 | 2.41 | 2.41 | 2.6 |
| No. reflections | 15228 | 20340 | 15165 | 27448 | 20276 |
| No. reflection used for R-free | 1524 (10%) | 2004 (9.85%) | 1063 (7%) | 1370 (5%) | 2019 (9.96%) |
| *R*_work_ / *R*_free_ | 20.39 / 24.26 | 22.18 / 25.2 | 21.36 / 24.6 | 25.0 / 28.8 | 23.64 / 26.58 |
| No. atoms |  |  |  |  |  |
| Protein | 5490 | 5472 | 2761 | 5393 | 5362 |
| lipid/ion | 231 | 250 | 148 | 231 | 255 |
| ligand | 108 | 108 | 48 | 100 | 112 |
| Water | 27 | 7 | 46 | 37 | 32 |
| *B*-factors |  |  |  |  |  |
| Protein | 49.68 | 67.89 | 50.19 | 57.24 | 44.66 |
| lipid/ion | 52.48 | 65.57 | 59.49 | 62.32 | 43.56 |
| ligand | 56.89 | 79.39 | 49.32 | 66.07 | 51.05 |
| Water | 45.66 | 62.72 | 57.08 | 57.77 | 44.08 |
| R.m.s. deviations |  |  |  |  |  |
| Bond lengths (Å) | 0.003 | 0.003 | 0.005 | 0.003 | 0.003 |
| Bond angles (°) | 0.61 | 0.61 | 1.04 | 0.58 | 0.59 |

**Supplementary Information Table 2** | **14 of the highest-affinity direct docking hits for the σ_2_ receptor.** See **Supplementary Information Table 3** for all 506 compounds tested.

| 2D drawing | ZINC ID | Rank | DOCK score (kcal/mol) | TC^†^ | K_i_ (nM) | | Selectivity  (σ_1_/σ_2_) |
| --- | --- | --- | --- | --- | --- | --- | --- |
|  |  |  |  |  | σ_2_ | σ_1_ |  |
| 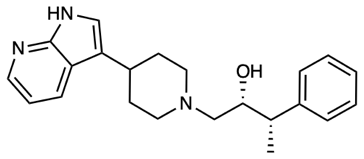 | ZINC000450573233 | 4429 | -57.25 | 0.32 | 4.3 | 128 | 30 |
| 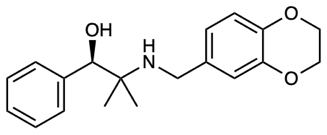 | ZINC000895657866 | 19047 | -55.35 | 0.31 | 21.4 | 989.6 | 46 |
| 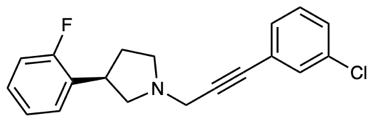 | ZINC001170548029 | 4945 | -57.11 | 0.35 | 22.6 | 727.2 | 32 |
| 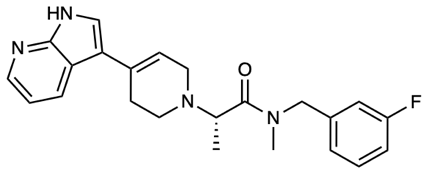 | ZINC000533478938 | 18545 | -55.38 | 0.30 | 34.5 | 1470 | 43 |
| 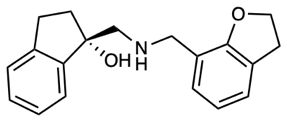 | ZINC000921927365 | 983 | -59.01 | 0.31 | 67.3 | 1186 | 18 |
| 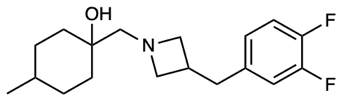 | ZINC000548355486 | 7007 | -56.68 | 0.29 | 2.4 | 4.9 | 2 |
| 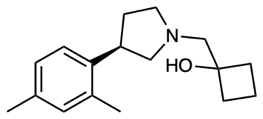 | ZINC000348332392 | 931 | -59.07 | 0.28 | 33.7 | 2.9 | 0.1 |
| 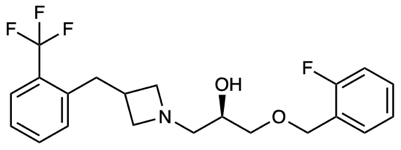 | ZINC001254761628 | 16059 | -55.58 | 0.27 | 4.7 | 53 | 11.3 |
| 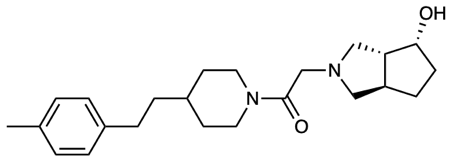 | ZINC000544117725 | 3522 | -57.52 | 0.28 | 10 | 16.25 | 1.6 |
| 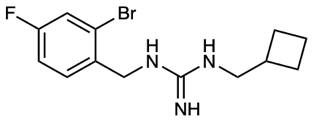 | ZINC000170908795 | 13281 | -55.84 | 0.29 | 6.7 | 32.7 | 4.9 |
| 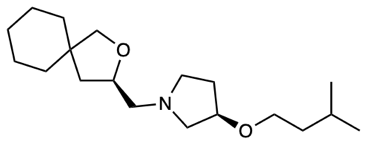 | ZINC001196519317 | 9290 | -56.3 | 0.29 | 2.4 | 13.4 | 5.6 |
| 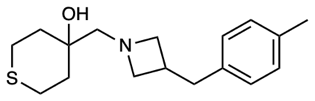 | ZINC000656714762 | 1276 | -58.68 | 0.26 | 67.8 | 4.6 | 0.1 |
| 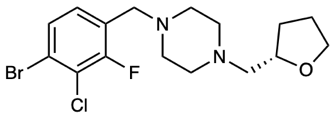 | ZINC001237901728 | 11409 | -56.03 | 0.30 | 27 | 1.6 | 0.1 |
| 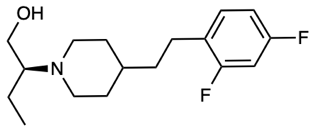 | ZINC001460312963 | 11817 | -55.99 | 0.29 | 5.2 | 1.7 | 0.3 |

^†^TC, Tanimoto coefficient to sigma ligands from ChEMBL.

**Supplementary Information Table 3** | 506 compounds tested for binding at the σ_2_ or σ_1_ receptors.

| ZINC ID | DOCK score (kcal/mol) | Global rank | TC† | % binding at 1 µM | | Ki (nM) | | selectivity (σ_1_/ σ_2_) | SMILES |
| --- | --- | --- | --- | --- | --- | --- | --- | --- | --- |
|  |  |  |  | σ_2_ | σ_1_ | σ_2_ | σ_1_ |  |  |
| ZINC000450573233 | -57.25 | 4628 | 0.32 | 3.6% | 33.6% | 4.3 | 128 | 30 | C[C@@H](c1ccccc1)[C@@H](O)CN1CCC(c2c[nH]c3ncccc23)CC1 |
| ZINC000895657866 | -55.35 | 19964 | 0.31 | 3.0% | 73.0% | 21.4 | 989.6 | 46 | CC(C)(NCc1ccc2c(c1)OCCO2)[C@H](O)c1ccccc1 |
| ZINC001170548029 | -57.11 | 5181 | 0.35 | 8.4% | 78.5% | 22.6 | 727.2 | 32 | Fc1ccccc1[C@H]1CCN(CC#Cc2cccc(Cl)c2)C1 |
| ZINC000533478938 | -55.38 | 19485 | 0.30 | 2.7% | 70.6% | 34.5 | 1470 | 43 | C[C@@H](C(=O)N(C)Cc1cccc(F)c1)N1CC=C(c2c[nH]c3ncccc23)CC1 |
| ZINC000921927365 | -59.01 | 1007 | 0.31 | 12.5% | 68.9% | 67.3 | 1186 | 18 | O[C@@]1(CNCc2cccc3c2OCC3)CCc2ccccc21 |
| ZINC000548355486 | -56.68 | 7303 | 0.29 | 0.4% | 1.0% | 2.4 | 4.9 | 2 | CC1CCC(O)(CN2CC(Cc3ccc(F)c(F)c3)C2)CC1 |
| ZINC000348332392 | -59.07 | 949 | 0.28 | 5.2% | -0.8% | 33.7 | 2.9 | 0.1 | Cc1ccc([C@H]2CCN(CC3(O)CCC3)C2)c(C)c1 |
| ZINC001254761628 | -55.58 | 16934 | 0.27 | 1.1% | 11.4% | 4.7 | 53 | 11.3 | O[C@@H](COCc1ccccc1F)CN1CC(Cc2ccccc2C(F)(F)F)C1 |
| ZINC000544117725 | -57.52 | 3655 | 0.28 | 0.0% | 5.0% | 10 | 16.25 | 1.6 | Cc1ccc(CCC2CCN(C(=O)CN3C[C@@H]4CC[C@@H](O)[C@H]4C3)CC2)cc1 |
| ZINC000170908795 | -55.84 | 13862 | 0.29 | 0.9% | 4.8% | 6.7 | 32.7 | 4.9 | N/C(=N\Cc1ccc(F)cc1Br)NCC1CCC1 |
| ZINC001196519317 | -56.30 | 9771 | 0.29 | 1.2% | 1.1% | 2.4 | 13.4 | 5.6 | CC(C)CCO[C@@H]1CCN(C[C@H]2CC3(CO2)CCCCC3)C1 |
| ZINC000656714762 | -58.68 | 1325 | 0.26 | 9.3% | -0.2% | 67.8 | 4.6 | 0.1 | Cc1ccc(CC2CN(CC3(O)CCSCC3)C2)cc1 |
| ZINC001237901728 | -56.03 | 12026 | 0.30 | 4.5% | -0.8% | 27 | 1.6 | 0.1 | Fc1c(Cl)c(Br)ccc1CN1CCN(C[C@@H]2CCCO2)CC1 |
| ZINC001460312963 | -55.99 | 12409 | 0.29 | 8.0% | -0.7% | 5.2 | 1.7 | 0.3 | CC[C@@H](CO)N1CCC(CCc2ccc(F)cc2F)CC1 |
| ZINC000188287346 | -59.56 | 600 | 0.32 | 0.9% | 8.1% | NT | NT | ND | C[C@H](Cn1ncc2ccccc2c1=O)N[C@H]1CC[C@H](Cc2ccccc2)C1 |
| ZINC000657922756 | -55.26 | 21317 | 0.25 | 1.7% | 23.2% | NT | NT | ND | C[C@@H](Cc1ccc2c(c1)CCCC2)N[C@H]1CCN(C2CCOCC2)C1=O |
| ZINC001308961074 | -56.76 | 6906 | 0.31 | 2.4% | 3.0% | NT | NT | ND | FC(F)(F)c1ccnc([C@H]2CCN(CCCc3ccc4c(c3)CCO4)C2)n1 |
| ZINC000792998860 | -56.66 | 7432 | 0.34 | 3.8% | 34.1% | NT | NT | ND | O=[N+]([O-])c1cccc([C@H](O)CN2CC(Cc3cccc(C(F)(F)F)c3)C2)c1 |
| ZINC000433044150 | -57.60 | 3425 | 0.33 | 5.0% | 4.4% | NT | NT | ND | CCc1ccc([C@@H]2CCN(Cc3cc(=O)n4c(C)cccc4n3)C2)cc1 |
| ZINC000897616680 | -57.17 | 4931 | 0.30 | 5.1% | 3.5% | NT | NT | ND | FC(F)(F)c1ccnc([C@H]2CCN(C[C@@H]3C[C@H]3c3ccccc3)C2)n1 |
| ZINC000374368469 | -55.43 | 18778 | 0.24 | 5.8% | 9.0% | NT | NT | ND | Cc1cc(C)cc(NC(=O)[C@H](C)N2CC[C@@H](C3CCOCC3)C2)c1 |
| ZINC000262856881 | -55.59 | 16724 | 0.30 | 6.4% | 5.6% | NT | NT | ND | C=C(C)CN/C(N)=N/[C@H]1C[C@H](c2cccc(F)c2)C1 |
| ZINC000430988927 | -57.10 | 5206 | 0.35 | 7.5% | 3.5% | NT | NT | ND | O=C(C[C@@H]1CCCN1CCOc1cccc([N+](=O)[O-])c1)c1cccs1 |
| ZINC000269280100 | -56.02 | 12047 | 0.27 | 11.1% | 41.1% | NT | NT | ND | N/C(=N\Cc1cccc2c1OCCCO2)NC1CCCC1 |
| ZINC000353995714 | -55.27 | 21126 | 0.30 | 13.2% | 7.3% | NT | NT | ND | Cc1cc(CN2CC[C@H]2Cc2ccccc2)ccc1-n1cncn1 |
| ZINC000657934399 | -65.05 | 5 | 0.30 | 13.3% | 9.0% | NT | NT | ND | C[C@H](Cn1ncc2ccccc2c1=O)N[C@@H](C)Cc1ccc2c(c1)CCCC2 |
| ZINC001168222793 | -55.24 | 21642 | 0.34 | 13.7% | 18.2% | NT | NT | ND | Cc1cc(C)cc(C2=CCN(CCC(=O)N3CCc4ccccc43)CC2)c1 |
| ZINC000507809396 | -55.47 | 18248 | 0.31 | 14.0% | -0.1% | NT | NT | ND | CC[C@H]1CN(Cc2ccc(-c3ccccc3)cc2F)C[C@H]1O |
| ZINC000398583627 | -55.24 | 21569 | 0.33 | 14.6% | 22.4% | NT | NT | ND | COc1ccc2cc(CN[C@](C)(CO)C(C)C)ccc2c1 |
| ZINC000549824186 | -58.09 | 2227 | 0.34 | 20.0% | 0.6% | NT | NT | ND | FC(F)(F)c1cccc([C@@]2(F)CCN(C[C@@H]3CCSC3)C2)c1 |
| ZINC000483826940 | -55.48 | 18107 | 0.24 | 22.5% | NT | NT | NT | ND | C[C@H](NC[C@H](CO)Cc1c(F)cccc1Cl)c1ccco1 |
| ZINC000886836194 | -56.02 | 12106 | 0.25 | 24.1% | NT | NT | NT | ND | Cc1cc(C(=O)CN2CC[C@@H]([C@H]3CCCO3)C2)c(C)n1C1CC1 |
| ZINC000453142034 | -55.65 | 15947 | 0.33 | 24.9% | NT | NT | NT | ND | C[C@H](CCCc1cccnc1)N[C@@H](C)Cn1cccn1 |
| ZINC000925887162 | -55.72 | 15238 | 0.32 | 27.4% | NT | NT | NT | ND | CCn1ncc2c1CCC[C@@H]2N[C@@H](C)Cc1ccccn1 |
| ZINC000933438523 | -55.90 | 13298 | 0.21 | 28.1% | NT | NT | NT | ND | Cc1csc2nc(CN[C@](C)(CO)Cc3ccc(F)cc3)cc(=O)n12 |
| ZINC000249076038 | -57.79 | 2914 | 0.29 | 30.5% | NT | NT | NT | ND | O[C@]1(CN[C@@H]2CC[C@@H](Cc3ccccc3)C2)CCSC1 |
| ZINC000621267824 | -58.94 | 1065 | 0.25 | 33.4% | NT | NT | NT | ND | CC(C)(CO)N1CCN(c2nc3c(cccc3Br)s2)CC1 |
| ZINC000934332177 | -57.44 | 3965 | 0.28 | 34.0% | NT | NT | NT | ND | O[C@H](CN1CC[C@@H](Cc2nccs2)C1)c1c(F)cccc1F |
| ZINC000567338231 | -55.65 | 15966 | 0.24 | 34.7% | NT | NT | NT | ND | Cc1csc2nc(CN[C@H]3C[C@H](OCc4ccccc4)C3(C)C)cc(=O)n12 |
| ZINC000894101819 | -58.11 | 2199 | 0.24 | 35.0% | NT | NT | NT | ND | CC[C@@H](C)N1CCN(c2nc(N)nc(C(F)(F)C(F)(F)F)n2)CC1 |
| ZINC000452107481 | -55.25 | 21450 | 0.22 | 36.9% | NT | NT | NT | ND | CC(C)N1CC[C@@H](N(C)S(=O)(=O)c2c(F)c(F)cc(F)c2F)C1 |
| ZINC000452023252 | -56.36 | 9264 | 0.28 | 37.5% | NT | NT | NT | ND | O=C([C@@H]1C[C@H]1C(F)(F)F)N(CCN1CCCCCC1)C1CCC1 |
| ZINC000801571276 | -55.91 | 13158 | 0.31 | 37.6% | NT | NT | NT | ND | O[C@H](CN[C@@H]1CCc2ccccc2OC1)COc1c(Cl)cccc1Cl |
| ZINC000448446275 | -57.11 | 5160 | 0.33 | 38.7% | NT | NT | NT | ND | CCC(CC)n1ccc(CN2CC=C(c3ccc(O)cc3)CC2)n1 |
| ZINC000662345330 | -55.86 | 13700 | 0.29 | 42.3% | NT | NT | NT | ND | CN(CC(=O)Nc1c(Cl)cccc1Cl)C[C@@H]1CC1(C)C |
| ZINC000352856249 | -56.65 | 7458 | 0.33 | 43.0% | NT | NT | NT | ND | N#Cc1ccc(N2CCN(C[C@H](O)CC3CCCC3)CC2)c(F)c1 |
| ZINC000846106280 | -56.59 | 7851 | 0.23 | 43.6% | NT | NT | NT | ND | Fc1ccc(C2OCCO2)c(Cl)c1CNC1CC2(CCC2)C1 |
| ZINC000262658947 | -57.72 | 3100 | 0.28 | 45.4% | NT | NT | NT | ND | COc1ccc(Br)cc1C/N=C1/NC[C@@H](C)N1 |
| ZINC000924470947 | -57.35 | 4268 | 0.28 | 45.5% | NT | NT | NT | ND | CC(C)(CN1C(=O)N2CCC[C@@H]3C[C@@]32C1=O)N1CCc2ccccc2C1 |
| ZINC000595632104 | -56.68 | 7309 | 0.33 | 45.7% | NT | NT | NT | ND | N#Cc1sccc1N1CCN(CCCCCn2cccn2)CC1 |
| ZINC000472356611 | -57.36 | 4219 | 0.31 | 45.9% | NT | NT | NT | ND | C[C@@H]1[C@H](Cc2ccccc2)CCN1Cc1cc(=O)n2cc(Cl)ccc2n1 |
| ZINC000574654702 | -55.62 | 16393 | 0.28 | 48.4% | NT | NT | NT | ND | CCc1nc2cc(CN[C@@H](C)CC(=O)N3CCCCCC3)ccc2n1C1CC1 |
| ZINC000656508398 | -57.20 | 4809 | 0.29 | 55.9% | NT | NT | NT | ND | OC[C@]1(CNC2=NCCCN2)CCc2ccccc21 |
| ZINC000296612417 | -56.76 | 6865 | 0.31 | 59.2% | NT | NT | NT | ND | C[C@H](COc1ccc(Cl)c(Cl)c1)N[C@@H](C)Cn1cncn1 |
| ZINC000336580930 | -55.27 | 21121 | 0.31 | 60.9% | NT | NT | NT | ND | O=C1NCCC[C@@H]1N1CC[C@@H](c2cccc(Cl)c2)C1 |
| ZINC000176995469 | -55.90 | 13220 | 0.31 | 61.4% | NT | NT | NT | ND | C[C@H](c1ccccn1)N1CC=C(c2ccc(O)cc2)CC1 |
| ZINC000571080072 | -60.05 | 410 | 0.30 | 64.0% | NT | NT | NT | ND | C[C@H](C(=O)NCc1ccc(Cl)cc1)N1CC[C@@H](c2cccnc2)C1 |
| ZINC000133991118 | -55.24 | 21534 | 0.33 | 65.4% | NT | NT | NT | ND | C[C@@H](Cn1cc(Br)cn1)NCCOc1ccc2c(c1)OCO2 |
| ZINC000473272986 | -56.23 | 10254 | 0.29 | 66.5% | NT | NT | NT | ND | COc1ccnc(N2CCN([C@H](C)CCSc3ccccc3)CC2)n1 |
| ZINC000247015101 | -57.45 | 3921 | 0.28 | 73.0% | NT | NT | NT | ND | O=C1[C@H](N2C[C@@H]3CCC[C@H]3C2)CCN1c1ccccc1Cl |
| ZINC000131571127 | -55.18 | 22496 | 0.22 | 76.6% | NT | NT | NT | ND | C[C@H](Cn1cc(Br)cn1)NCc1cn2cc(Cl)ccc2n1 |
| ZINC000407281203 | -56.29 | 9803 | 0.26 | 80.8% | NT | NT | NT | ND | CCc1cc(NC(=O)[C@@H]2CC[C@H](C3CC3)N2)ccc1C |
| ZINC000528002641 | -55.42 | 18937 | 0.23 | 84.0% | NT | NT | NT | ND | C[C@@H](N[C@H]1Cc2ccccc2NC1=O)[C@@H]1C[C@H]1c1ccc(Cl)s1 |
| ZINC000248559983 | -56.91 | 6084 | 0.31 | 87.4% | NT | NT | NT | ND | CN(C[C@@H](O)CCc1ccccc1)[C@H]1CCN(c2ccccc2F)C1=O |
| ZINC000093013680 | -55.88 | 13429 | 0.33 | 89.6% | NT | NT | NT | ND | c1sc(CN2CCC3(C2)OCCO3)cc1-c1ccccc1 |
| ZINC000769519341 | -55.28 | 21009 | 0.32 | 91.7% | NT | NT | NT | ND | C[C@H](CCN1C(=O)c2ccccc2C1=O)NC[C@@H](O)c1c(F)cccc1F |
| ZINC000911907143 | -56.14 | 11006 | 0.31 | 93.1% | NT | NT | NT | ND | Cc1ccc(NC(=O)C23CCC(CC2)N3)cc1[N+](=O)[O-] |
| ZINC000893277758 | -55.30 | 20724 | 0.27 | 95.1% | NT | NT | NT | ND | Cc1cccc2sc(N3CCN(CCO)[C@H](C)C3)nc21 |
| ZINC000892713700 | -57.10 | 5220 | 0.22 | 99.5% | NT | NT | NT | ND | CC[C@H](CO)N1CCN(c2cc(Cl)c3cnn(C)c3n2)CC1 |
| ZINC000435004139 | -55.91 | 13117 | 0.32 | 99.6% | NT | NT | NT | ND | CN(C(=O)CN1CCC(c2cnc[nH]2)CC1)C(c1ccccc1)c1ccccc1 |
| ZINC000659267272 | -56.47 | 8587 | 0.25 | 101.0% | NT | NT | NT | ND | CCOc1cc2c(cc1NC(=O)C[C@@H]1NCc3ccccc31)O[C@@H](C)C2 |
| ZINC000820594289 | -55.60 | 16659 | 0.21 | 103.8% | NT | NT | NT | ND | C[C@H]1[C@H](N(C)C)CCN1c1snc(Cl)c1C#N |
| ZINC000296435291 | -55.25 | 21422 | 0.25 | 106.5% | NT | NT | NT | ND | CS[C@H]1CCN([C@H](C)c2nc(-c3ccccc3C)no2)C1 |
| ZINC000665143541 | -55.34 | 20094 | 0.21 | 108.2% | NT | NT | NT | ND | CN(C)Cc1ccc(-c2cscc2CCO)cc1F |
| ZINC000777733869 | -57.03 | 5539 | 0.33 | 108.4% | NT | NT | NT | ND | O=C1NCCC[C@@H]1N1CC[C@@](O)(c2ccc(F)cc2)C1 |
| ZINC000369205980 | -55.59 | 16735 | 0.31 | 111.8% | NT | NT | NT | ND | N#CC1(c2ccccn2)CCN([C@@H]2Cc3ccccc3[C@@H]2O)CC1 |
| ZINC000287374567 | -55.63 | 16207 | 0.31 | 113.5% | NT | NT | NT | ND | O[C@H]1c2ccccc2C[C@H]1N1Cc2cccc(Cl)c2C1 |
| ZINC000681377109 | -55.51 | 17760 | 0.31 | 113.9% | NT | NT | NT | ND | Fc1cc(Br)cnc1N1CCN(CCc2cccs2)CC1 |
| ZINC001525937517 | -57.41 | 4069 | 0.35 | 117.0% | NT | NT | NT | ND | O=C(OC[C@@H]1CCN1Cc1ccccc1)c1nccc2occc21 |
| ZINC000129576345 | -32.64 | 57528707 | 0.31 | 14.2% | 24.2% | NT | NT | ND | COc1ccccc1O[C@H](C)CN[C@@H](C)c1ccccc1OC |
| ZINC000550829396 | -19.00 | 320047299 | 0.19 | 19.6% | 28.6% | NT | NT | ND | Cn1nccc1[C@@H]1CCC[C@@H](N[C@H]2CC3CCC2CC3)C1 |
| ZINC000182842742 | -33.47 | 50889836 | 0.30 | 36.1% | 66.7% | NT | NT | ND | Cc1ccc([N+](=O)[O-])cc1[C@H](C)N1C[C@H]2CCC[C@@H]2C1 |
| ZINC000658086473 | -42.01 | 10446238 | 0.31 | 42.3% | 54.9% | NT | NT | ND | COc1ccc([C@@H](C)N[C@H](CO)C2CCC2)cc1Br |
| ZINC000635049325 | -40.64 | 14542581 | 0.31 | 53.2% | 3.4% | NT | NT | ND | O=C(Nc1cscc1Cl)C(=O)N[C@@H]1CCN(CC2CC2)C1 |
| ZINC000389015736 | -19.08 | 318290882 | 0.30 | 54.1% | 65.5% | NT | NT | ND | C[C@H](N[C@@H]1CCCc2c3ccccc3[nH]c21)[C@H]1CCCOC1 |
| ZINC001195393353 | -29.56 | 91360310 | 0.34 | 54.5% | 24.0% | NT | NT | ND | COc1ccc2c(c1)C[C@H](CN1CCCC[C@H]1CCC(=O)OC(C)(C)C)O2 |
| ZINC000582751592 | -44.91 | 4410937 | 0.26 | 72.4% | 70.7% | NT | NT | ND | O[C@@H]1CC[C@@H]2CN(Cc3cccc4c[nH]nc43)C[C@H]12 |
| ZINC001420054689 | -34.38 | 44501635 | 0.28 | 78.5% | 54.6% | NT | NT | ND | COCC1(C(=O)N2CCC[C@@H](N(C)CC[C@H](C)F)C2)CC1 |
| ZINC000649929688 | -34.10 | 46367996 | 0.23 | 96.5% | 71.5% | NT | NT | ND | O=C(CN1CCC[C@@H]1[C@@H]1CCC[C@H]1O)N1CCOC[C@H]1C1CC1 |
| ZINC000245533477 | -21.78 | 254547099 | 0.21 | 98.5% | 70.8% | NT | NT | ND | CC[C@H](C)CNC[C@H]1CCC[C@H]1NS(C)(=O)=O |
| ZINC001087646081 | -27.94 | 116424252 | 0.32 | 99.4% | 8.8% | NT | NT | ND | CC[C@@H]1[C@@H](NC(=O)Cc2ccc(C)cc2)CCN1C[C@@H]1CCOC1 |
| ZINC000341348768 | -35.78 | 35881111 | 0.16 | 103.4% | 80.7% | NT | NT | ND | Cc1cnc(CN[C@@H](CO)CCC(C)(C)C)n1C |
| ZINC000662800454 | -32.45 | 59205541 | 0.28 | 105.6% | 86.9% | NT | NT | ND | COc1cc(NC(=O)CC2(N)CC2)cc(OC)c1Br |
| ZINC000261774189 | -23.44 | 213383997 | 0.33 | 108.3% | 53.9% | NT | NT | ND | C=CCOCC(=O)N(CCN)CCc1ccccc1 |
| ZINC000543048256 | -37.34 | 27800769 | 0.33 | 109.0% | 59.2% | NT | NT | ND | COc1cccc2ncnc(N3CCN(CC4CCOCC4)CC3)c21 |
| ZINC001078073018 | -27.42 | 125865808 | 0.25 | 111.2% | 83.4% | NT | NT | ND | C[C@@H](CCNC(=O)C1(CN(C)C)CC1)NC(=O)c1ncccc1O |
| ZINC000948091407 | -44.72 | 4707826 | 0.29 | 112.2% | 88.9% | NT | NT | ND | CCN1CC[C@@H](NC(=O)c2ccnc(-n3cncn3)c2)C[C@@H]1C |
| ZINC001655120594 | -36.27 | 33238652 | 0.27 | 112.4% | 66.8% | NT | NT | ND | C[C@H](OCCCNC(=O)CN1C[C@@H](C)OCC[C@H]1C)c1ccccc1 |
| ZINC000996610565 | -28.98 | 99710181 | 0.24 | 113.9% | 99.6% | NT | NT | ND | CC(C)NC1CCN(C(=O)Cn2cnc(C(N)=O)n2)CC1 |
| ZINC000853031922 | -38.74 | 21590638 | 0.32 | 115.3% | 33.3% | NT | NT | ND | Fc1ccc(NC(=S)NC2CCN([C@H]3CCOC3)CC2)cc1 |
| ZINC001035320653 | -11.33 | 424937722 | 0.34 | 118.8% | 21.6% | NT | NT | ND | CCc1ocnc1C(=O)NC[C@H]1CN(C[C@@H](C)CC)CCO1 |
| ZINC000369129048 | -26.35 | 146706740 | 0.21 | 119.8% | 94.9% | NT | NT | ND | Cn1cc(-c2ncc(CN(CC3CC3)C[C@H]3CN(C)CCO3)cn2)cn1 |
| ZINC000416873685 | -23.24 | 218337454 | 0.26 | 126.9% | 90.7% | NT | NT | ND | C[C@H](N)[C@@H]1CCCCN1C(=O)NCc1cn2ccccc2n1 |
| ZINC000906421824 | -17.77 | 345293233 | 0.22 | 127.3% | 86.6% | NT | NT | ND | C[C@@H]1CN(C(=O)C2CN(C)CCN(C)C2)CC(C)(C)O1 |
| ZINC000661577595 | -52.50 | 129452 | 0.33 | 6.4% | NT | 1.757 | NT | ND | C(CN1CC[C@@H](C2CCCCC2)C1)C1CCOCC1 |
| ZINC000440321606 | -61.12 | 152 | 0.27 | 11.9% | NT | 4.15 | NT | ND | CCOC(=O)[C@@H](CC)N1CCN(C2CCC(C(C)C)CC2)CC1 |
| ZINC000878406056 | -59.57 | 596 | 0.23 | 11.8% | NT | 7.656 | NT | ND | CC(C)(C)[C@@H]1CCN(C[C@@H](O)CC2(O)CCCCC2)C1 |
| ZINC001460371878 | -63.30 | 19 | 0.28 | 10.9% | NT | 7.924 | NT | ND | O[C@@H](CN1CC[C@@H](C2CCCCC2)C1)CC1(O)CCCCC1 |
| ZINC000409447221 | -55.00 | 25628 | 0.29 | 9.7% | NT | 8.847 | NT | ND | CC(C)(C)[C@@H]1CCN(C[C@@H](O)COCc2ccccc2F)C1 |
| ZINC000559413424 | -59.40 | 715 | 0.29 | 11.9% | NT | 10.49 | NT | ND | Cc1cccc(C)c1OC[C@H](O)CN1CC[C@@H](CC2CC2)C1 |
| ZINC000191344346 | -52.50 | 129135 | 0.34 | 8.7% | NT | 11.75 | NT | ND | O=C(CC1CCCC1)NC1CCN(C[C@H](O)C2CCCCC2)CC1 |
| ZINC000480785335 | -52.50 | 129321 | 0.33 | 9.8% | NT | 12.87 | NT | ND | CN(C[C@]1(O)CCN(Cc2ccccc2)C1)C(=O)c1ccc(F)c2ccccc12 |
| ZINC001376084945 | -55.00 | 25751 | 0.30 | 9.7% | NT | 13.52 | NT | ND | C[C@H](C(=O)NC[C@]1(O)CCN(CCC2CCCC2)C1)c1ccccc1 |
| ZINC000605902355 | -61.39 | 118 | 0.34 | 11.5% | NT | 13.73 | NT | ND | O[C@@H](COCc1ccccc1Cl)CN1CC[C@H](Cc2ccc(F)cc2)C1 |
| ZINC000872109019 | -55.00 | 25696 | 0.29 | 11.6% | NT | 14.71 | NT | ND | c1ccc(N2CCC(NC[C@@H]3C[C@H]4CCC[C@@H]4O3)CC2)cc1 |
| ZINC000131303503 | -55.00 | 25593 | 0.33 | 10.7% | NT | 14.74 | NT | ND | CC(C)c1ccc(NC(=O)C(=O)NC2CCC(N3CCC(C)CC3)CC2)cc1 |
| ZINC000960887654 | -47.50 | 1644724 | 0.26 | 12.9% | NT | 18.11 | NT | ND | CCc1nnsc1C(=O)N[C@@H]1[C@H]2CN(CC3CCCCCC3)C[C@H]21 |
| ZINC000301801549 | -59.57 | 595 | 0.32 | 6.9% | NT | 19.64 | NT | ND | c1ccc(CSc2cncc(N3CCC[C@H](N4CCCC4)CC3)n2)cc1 |
| ZINC000941634458 | -50.00 | 513561 | 0.27 | 12.7% | NT | 20.26 | NT | ND | C[C@@H](C(=O)N1CCN(C2CN(CC3CC(C)C3)C2)CC1)C1CCCC1 |
| ZINC000481989921 | -52.50 | 129323 | 0.32 | 11.7% | NT | 21.59 | NT | ND | Cc1cc2[nH]c(CN3CCC[C@H](N4CCCC4)CC3)cc2c(C)c1 |
| ZINC000878084395 | -61.07 | 161 | 0.27 | 27.7% | NT | NT | NT | ND | O[C@@H](CN1CC[C@@H](c2ccccc2F)C1)CC1(O)CCCCC1 |
| ZINC000849049798 | -61.21 | 140 | 0.23 | 8.7% | NT | NT | NT | ND | C[C@H]1CN/C(=N/CCCc2c(Cl)cccc2Cl)N1 |
| ZINC000076836015 | -61.05 | 163 | 0.30 | 60.5% | NT | NT | NT | ND | c1ccc2c(c1)nc(-c1cccnc1)nc2N1CCC[C@H](N2CCCC2)C1 |
| ZINC000766031584 | -61.44 | 112 | 0.33 | 63.2% | NT | NT | NT | ND | Cc1ccc2c(c1)CC[C@H]2N[C@@H](CO)Cc1ccccc1 |
| ZINC000684607185 | -62.40 | 50 | 0.34 | 17.3% | NT | NT | NT | ND | CC(C)C1CN(C[C@@H](O)COC(c2ccccc2)c2ccccc2)C1 |
| ZINC000683721842 | -61.32 | 122 | 0.30 | 99.8% | NT | NT | NT | ND | Clc1ccccc1Cn1cc(CNc2nc3ccccc3[nH]2)cn1 |
| ZINC000676461131 | -62.78 | 32 | 0.33 | 63.8% | NT | NT | NT | ND | Clc1ccccc1Cn1cc(CN2CCC[C@@H]2c2ccc[nH]2)cn1 |
| ZINC000668631550 | -62.67 | 38 | 0.27 | 58.4% | NT | NT | NT | ND | Cc1ccccc1-n1cc(CN2CCC[C@@H]([C@H]3CCCCO3)C2)nn1 |
| ZINC000662335416 | -60.95 | 180 | 0.28 | 121.5% | NT | NT | NT | ND | O=C(CN1CC2CC1(c1ccccc1)C2)NC[C@@H]1CCCO1 |
| ZINC000656499190 | -62.15 | 60 | 0.32 | 71.5% | NT | NT | NT | ND | Cc1cc(F)c(CN[C@@H]2CCN(c3ccccc3C(N)=O)C2)c(Cl)c1 |
| ZINC000617924178 | -62.67 | 37 | 0.27 | 125.6% | NT | NT | NT | ND | CCN(C(=O)c1cccc(C(F)(F)F)n1)[C@H]1CCN(CC)C1 |
| ZINC000609803478 | -63.53 | 14 | 0.32 | 49.8% | NT | NT | NT | ND | COC(=O)[C@H](NCC(C)(C)N1CCc2ccccc2C1)c1ccccc1 |
| ZINC000595790972 | -62.44 | 47 | 0.32 | 96.1% | NT | NT | NT | ND | Cc1cc([N+](=O)[O-])ccc1NC(=O)CN1CCC[C@@H]([C@@H]2CCCCO2)C1 |
| ZINC000577603040 | -60.89 | 189 | 0.34 | 61.7% | NT | NT | NT | ND | C[C@H](Cc1ccccc1Br)NCC1(O)CCC1 |
| ZINC000518003593 | -61.42 | 114 | 0.29 | 79.8% | NT | NT | NT | ND | Cc1ccccc1[C@H](OC[C@H](O)CN1Cc2ccccc2C1)c1ccccc1 |
| ZINC000512182007 | -62.09 | 63 | 0.32 | 31.6% | NT | NT | NT | ND | C[C@H]1C[C@H](N2CCCC2)CN1CC(=O)N(Cc1ccccc1)c1ccccc1 |
| ZINC000508475661 | -60.93 | 184 | 0.28 | 38.6% | NT | NT | NT | ND | Cc1cccc(CNc2nccn2Cc2ccccc2)c1C |
| ZINC000464767692 | -60.81 | 206 | 0.33 | 73.8% | NT | NT | NT | ND | CCN(CC)[C@H]1CCN(CC(=O)N[C@H](c2cccs2)c2ccccc2)C1 |
| ZINC000451319844 | -61.11 | 157 | 0.33 | 63.0% | NT | NT | NT | ND | CS[C@@H]1CCN(C[C@@H](O)COC(c2ccc(F)cc2)c2ccc(F)cc2)C1 |
| ZINC000430847713 | -61.93 | 74 | 0.32 | 67.3% | NT | NT | NT | ND | CCCN1C(=O)N(CC(C)(C)N2CCc3ccccc3C2)C(=O)[C@H]1CC |
| ZINC000429522320 | -61.11 | 156 | 0.31 | 60.1% | NT | NT | NT | ND | O[C@@H](CN[C@@H](Cc1ccccc1F)c1ccccc1)CN1CCCC1 |
| ZINC000426471156 | -61.11 | 155 | 0.31 | 36.3% | NT | NT | NT | ND | O[C@H](COC(c1ccccc1)c1ccccc1)CN1CC[C@@H]2CCC[C@H]21 |
| ZINC000421710406 | -60.85 | 197 | 0.32 | 81.8% | NT | NT | NT | ND | CC(C)Sc1ccc([N+](=O)[O-])cc1C(=O)N(C)C[C@H]1CCCN1C |
| ZINC000421710394 | -60.93 | 183 | 0.32 | 94.6% | NT | NT | NT | ND | CC(C)CSc1ccc([N+](=O)[O-])cc1C(=O)N(C)C[C@H]1CCCN1C |
| ZINC000421701270 | -61.46 | 109 | 0.33 | 76.1% | NT | NT | NT | ND | CCC[C@@H](C(=O)N(C)C[C@H]1CCCN1C)c1ccccc1 |
| ZINC000421699823 | -62.45 | 46 | 0.32 | 96.4% | NT | NT | NT | ND | COc1ccc(Oc2ccc([N+](=O)[O-])cc2C(=O)N(C)C[C@H]2CCCN2C)cc1 |
| ZINC000421295130 | -60.83 | 198 | 0.33 | 77.9% | NT | NT | NT | ND | CC(C)Sc1c(Cl)cccc1NC(=O)C(=O)N(C)C[C@H]1CCCN1C |
| ZINC000347007006 | -61.38 | 119 | 0.28 | 108.1% | NT | NT | NT | ND | COC(=O)[C@@H]1CCN([C@H](C)C(=O)NCc2ccccc2C)[C@@H]1C |
| ZINC000287581634 | -63.23 | 22 | 0.32 | 111.3% | NT | NT | NT | ND | COC(=O)C[C@H]1CCCN1C[C@@H](O)Cc1cccc(Cl)c1 |
| ZINC000263644082 | -64.51 | 6 | 0.32 | 83.4% | NT | NT | NT | ND | Cc1ccc(NC(=O)CN2CCC[C@@H](C3OCCO3)C2)cc1Cl |
| ZINC000248917461 | -61.69 | 90 | 0.28 | 103.1% | NT | NT | NT | ND | C[C@H]1CCN(CC#Cc2cccc(Cl)c2)[C@H]1CO |
| ZINC000247001079 | -63.06 | 24 | 0.30 | 61.2% | NT | NT | NT | ND | COC(=O)C[C@H]1CCCN1C[C@@H](O)Cc1ccc(C(F)(F)F)cc1 |
| ZINC000192043753 | -60.88 | 190 | 0.34 | 52.9% | NT | NT | NT | ND | O=C1Nc2ccccc2CC[C@H]1N1CC[C@@H](COCc2ccccc2)C1 |
| ZINC001474404955 | -61.56 | 101 | 0.26 | 62.3% | NT | NT | NT | ND | Cc1cc(C)c(/C=C\CN2CCC[C@@H](O)C2)cc1C |
| ZINC001362573982 | -61.32 | 123 | 0.33 | 94.8% | NT | NT | NT | ND | CN(C[C@H]1CCCN1C)C(=O)[C@]1(C)CC(c2ccccc2Cl)=NO1 |
| ZINC001206651458 | -61.46 | 111 | 0.23 | 109.0% | NT | NT | NT | ND | C[C@@H]1CN([C@H]2CCc3ccccc3NC2=O)C[C@H]1NC(=O)/C=C\C(C)(C)C |
| ZINC000894427229 | -59.55 | 613 | 0.22 | 38.7% | NT | NT | NT | ND | CCC(C)(C)N1CCN(c2nc(N)nc(C(F)(F)C(F)(F)F)n2)CC1 |
| ZINC000858362097 | -59.68 | 560 | 0.26 | 141.9% | NT | NT | NT | ND | C[C@H](Nc1nc(N)nc(N)n1)c1ccccc1C(F)(F)F |
| ZINC000830877226 | -59.47 | 670 | 0.29 | 78.8% | NT | NT | NT | ND | CS[C@@H]1CCN(C[C@@H](O)c2ccccc2C(F)(F)F)C1 |
| ZINC000819225653 | -59.65 | 568 | 0.28 | 17.6% | NT | NT | NT | ND | CC(C)(C)OC(=O)N1CCC[C@@](C)(CNC[C@@H](O)c2c(F)cccc2F)C1 |
| ZINC000074484650 | -59.43 | 694 | 0.27 | 86.9% | NT | NT | NT | ND | C[C@H](C(=O)NCc1cc(Br)cs1)N1CCCCCC1 |
| ZINC000743860012 | -59.48 | 662 | 0.32 | 23.7% | NT | NT | NT | ND | O[C@@H](CN1CC[C@@H](CSc2ccccc2)C1)c1c(F)cccc1F |
| ZINC000678272880 | -59.50 | 646 | 0.33 | 55.7% | NT | NT | NT | ND | c1ncn(-c2ccccc2CN2CCC[C@H](N3CCCC3)CC2)n1 |
| ZINC000672564445 | -59.66 | 564 | 0.25 | 69.7% | NT | NT | NT | ND | C[C@H]1[C@H](N2CCCC2)CCN1Cc1cn2cc(Br)ccc2n1 |
| ZINC000611661177 | -59.43 | 696 | 0.33 | 19.1% | NT | NT | NT | ND | COC(=O)[C@H](c1cccc(Cl)c1Cl)N1CCC[C@H](N2CCCC2)C1 |
| ZINC000550423124 | -59.69 | 553 | 0.30 | 43.5% | NT | NT | NT | ND | Cc1nc(CN2CC[C@H](CC3CC3)C2)nc2ccccc12 |
| ZINC000542825037 | -59.56 | 604 | 0.30 | 107.1% | NT | NT | NT | ND | C[C@@H](C(=O)NCc1ccccc1Cl)N1CC[C@H](n2ncc3ccccc32)C1 |
| ZINC000534109220 | -59.47 | 669 | 0.29 | 70.4% | NT | NT | NT | ND | Cc1cc(NC(=O)[C@H](C)N2CCC[C@@H]([C@H](C)O)C2)ccc1Br |
| ZINC000516964180 | -59.59 | 584 | 0.32 | 99.3% | NT | NT | NT | ND | Cc1cccc(NC(=O)N(C)C[C@H]2CCCN2C)c1C(=O)N1CCCC1 |
| ZINC000513818302 | -59.47 | 668 | 0.34 | 89.1% | NT | NT | NT | ND | Cc1ccccc1[C@@H]1CCN([C@H](C)C(=O)NCCc2ccccc2)C1 |
| ZINC000508455082 | -59.73 | 540 | 0.34 | 69.3% | NT | NT | NT | ND | Brc1ccccc1CNc1nccn1Cc1ccccc1 |
| ZINC000468894194 | -59.58 | 587 | 0.34 | 35.4% | NT | NT | NT | ND | CC(C)CO[C@H]1CCN([C@@H](C)C(=O)N(C)Cc2ccc(Br)cc2)C1 |
| ZINC000423867812 | -59.54 | 618 | 0.28 | 95.6% | NT | NT | NT | ND | C[C@H](c1ccccc1Br)N(C)C(=O)C[C@@H]1CCCN1 |
| ZINC000412048402 | -59.51 | 640 | 0.32 | 88.6% | NT | NT | NT | ND | C[C@H](C(=O)NCc1ccc(F)cc1)N1C[C@@H]2CC[C@H](O)C[C@@H]2C1 |
| ZINC000358253868 | -59.53 | 623 | 0.29 | 52.1% | NT | NT | NT | ND | Cc1ccc([C@H](NC[C@@]2(O)CCN(C)C2)c2ccccc2)cc1 |
| ZINC000186482223 | -59.46 | 674 | 0.33 | 95.3% | NT | NT | NT | ND | N#Cc1ccccc1NC(=O)CCN1CCC(O)(c2c(F)cccc2F)CC1 |
| ZINC001580410121 | -59.50 | 650 | 0.33 | 122.1% | NT | NT | NT | ND | N[C@H](Cc1ccccn1)C(=O)NCc1ccccc1C(F)(F)F |
| ZINC001320149247 | -59.45 | 686 | 0.31 | 14.7% | NT | NT | NT | ND | OC1(CN2CC(Cc3ccccc3C(F)(F)F)C2)CCC1 |
| ZINC000120571916 | -59.42 | 700 | 0.32 | 17.7% | NT | NT | NT | ND | O=C(Nc1cccc(OC(F)(F)F)c1)C(=O)N1CCC[C@@H](N2CCCC2)CC1 |
| ZINC001109713565 | -59.52 | 637 | 0.27 | 28.7% | NT | NT | NT | ND | C[C@H](C(=O)N[C@H]1C[C@H]2CC[C@@H]1N2CCOCC1CC1)c1cccs1 |
| ZINC000894429059 | -55.00 | 25702 | 0.25 | 33.6% | NT | NT | NT | ND | CCC(C)(C)N1CCN(c2cc(C(F)F)ncn2)CC1 |
| ZINC000886762228 | -55.00 | 25699 | 0.29 | 109.8% | NT | NT | NT | ND | C=CCC1(O)CCN([C@H](C)C(=O)N[C@@H](C)c2ccc(Cl)cc2)CC1 |
| ZINC000843092160 | -55.00 | 25693 | 0.26 | 16.2% | NT | NT | NT | ND | OC[C@@H](CNC[C@H](O)c1ccc(F)c(Br)c1)CC1CCCC1 |
| ZINC000825615260 | -54.99 | 25873 | 0.34 | 104.5% | NT | NT | NT | ND | O=C([C@H](O)c1cccc([N+](=O)[O-])c1)N1CC[C@H](N2CCCC2)C1 |
| ZINC000775588849 | -55.00 | 25687 | 0.34 | 39.0% | NT | NT | NT | ND | COc1ccc([C@H]2CCN(C[C@@H](O)c3ccc(F)cc3Cl)C2)cc1 |
| ZINC000645238909 | -54.99 | 25850 | 0.32 | 35.5% | NT | NT | NT | ND | Clc1ccc(O[C@H]2CCN(C3CCCC3)C2)cc1 |
| ZINC000589606863 | -55.00 | 25669 | 0.34 | 68.9% | NT | NT | NT | ND | C[C@H](CN(C)CC(=O)NCc1cccc(Cl)c1)c1ccccc1 |
| ZINC000584028046 | -55.00 | 25668 | 0.28 | 18.4% | NT | NT | NT | ND | Fc1cccc(Cl)c1CNC1CCC2(CCCO2)CC1 |
| ZINC000574285278 | -55.00 | 25666 | 0.24 | 76.2% | NT | NT | NT | ND | CCc1ccc([C@@H](C)N[C@@H]2CCCN(C(=O)CCC(F)(F)F)C2)s1 |
| ZINC000556573026 | -55.00 | 25660 | 0.26 | 16.7% | NT | NT | NT | ND | CN1CC[C@](O)(CN[C@@H](c2sccc2Br)C2CCCCC2)C1 |
| ZINC000532978237 | -55.00 | 25652 | 0.34 | 118.6% | NT | NT | NT | ND | COC(=O)c1ccccc1NC(=O)CN1C[C@@H](c2ccccc2)C[C@@H]1C |
| ZINC000531059007 | -55.00 | 25651 | 0.35 | 87.5% | NT | NT | NT | ND | CC1(C)[C@H](NC(=O)[C@]2(O)CCN(Cc3ccccc3)C2)[C@@H]2CCCO[C@@H]21 |
| ZINC000526962900 | -55.00 | 25649 | 0.25 | 60.7% | NT | NT | NT | ND | Cc1cccc(C[C@H](CO)NCc2cnc(C3CCCCC3)s2)c1 |
| ZINC000517634594 | -55.00 | 25648 | 0.28 | 98.7% | NT | NT | NT | ND | NC(=O)[C@H](Cc1ccc(Cl)cc1)NCc1cc2ccccc2s1 |
| ZINC000500192494 | -55.00 | 25646 | 0.33 | 69.3% | NT | NT | NT | ND | C[C@]1(c2ccccc2)CCCN(CC(=O)N2CCc3ccccc32)C1 |
| ZINC000466933503 | -55.00 | 25641 | 0.34 | 84.3% | NT | NT | NT | ND | C[C@@H]1CN(C2CC2)C[C@H]1NC(=O)N(Cc1cccc(Br)c1)C1CC1 |
| ZINC000453351921 | -55.00 | 25639 | 0.31 | 69.2% | NT | NT | NT | ND | COc1cccnc1CN[C@H](Cc1ccc(C)cc1)C1CC1 |
| ZINC000439311985 | -55.00 | 25636 | 0.31 | 106.4% | NT | NT | NT | ND | C[C@@H]1C[C@@H](c2ccccc2)CCN1Cc1nc(C2CC2)cs1 |
| ZINC000426526480 | -55.00 | 25634 | 0.34 | 92.0% | NT | NT | NT | ND | O=C(CN1CC[C@@H]2CCC[C@@H]21)Nc1cccc(I)c1 |
| ZINC000332414923 | -55.00 | 25618 | 0.35 | 87.2% | NT | NT | NT | ND | CC(C)CC(=O)N[C@@H](C(=O)N1CCC[C@H](N2CCCC2)C1)c1ccccc1 |
| ZINC000285725404 | -55.00 | 25614 | 0.31 | 114.6% | NT | NT | NT | ND | CCc1cccc(C)c1NC(=O)CN1C[C@@H](N2CCCC2)C[C@H]1C |
| ZINC000246894550 | -55.00 | 25607 | 0.34 | 72.6% | NT | NT | NT | ND | CCN(C[C@@H]1CCN(Cc2ccccc2)C1)C(=O)[C@H]1CC(=O)N(C2CCCC2)C1 |
| ZINC000245453841 | -55.00 | 25606 | 0.30 | 113.9% | NT | NT | NT | ND | C[C@@H](NC(=O)CNC(=O)CN1CCC[C@H]1C)c1ccc(F)cc1 |
| ZINC000228149300 | -55.00 | 25605 | 0.33 | 112.8% | NT | NT | NT | ND | Cc1ccccc1NC(=O)CN1CCC[C@@H]1C[C@H](C)O |
| ZINC000182372239 | -55.00 | 25603 | 0.27 | 34.6% | NT | NT | NT | ND | CCN(CC)[C@H]1CCN(Cc2ncc(Cl)cc2Cl)C1 |
| ZINC000176805277 | -55.00 | 25601 | 0.30 | 41.6% | NT | NT | NT | ND | COC(=O)[C@@H](NCC[C@H](C)N(C)Cc1ccccc1)c1cccc(C#N)c1 |
| ZINC001579778746 | -55.00 | 25753 | 0.26 | 19.6% | NT | NT | NT | ND | Cc1cccc2c1CCN2C(=O)[C@@H]1CC[C@H]2CCCC[C@@H]2N1 |
| ZINC001156463732 | -55.00 | 25728 | 0.33 | 32.6% | NT | NT | NT | ND | C(=C/c1ccccc1)\CNc1ccc2c(n1)CCC2 |
| ZINC000115566238 | -55.00 | 25590 | 0.35 | 18.2% | NT | NT | NT | ND | CC(C)OCCN1CCN(C2CCC(C)CC2)CC1 |
| ZINC001096194954 | -55.00 | 25718 | 0.29 | 118.2% | NT | NT | NT | ND | C[C@H](C(=O)N[C@H]1C[C@H]2CC[C@@H]1N2CCn1cccn1)c1ccccc1F |
| ZINC001095312060 | -55.00 | 25716 | 0.32 | 69.4% | NT | NT | NT | ND | Cc1ccc(CN2[C@@H]3CC[C@H]2[C@@H](NC(=O)c2cn(C)cn2)C3)c(C)c1 |
| ZINC000961091031 | -52.50 | 129608 | 0.25 | 33.0% | NT | NT | NT | ND | O=C(C1CCCC1)N1CCC[C@H]([C@@H]2CCN(C/C=C\Cl)C2)C1 |
| ZINC000932649435 | -52.50 | 129598 | 0.28 | 69.5% | NT | NT | NT | ND | CCOC(=O)[C@@H]1CCN(C[C@@H](O)COc2cccc(C)c2)C1 |
| ZINC000932000298 | -52.50 | 129596 | 0.23 | 75.9% | NT | NT | NT | ND | O=C1[C@@H](N2CC[C@@H](C3OCCO3)C2)CCN1c1cccc(Br)c1 |
| ZINC000839577413 | -52.50 | 129532 | 0.25 | 28.7% | NT | NT | NT | ND | C[C@@H]1CCC[C@H](CCN2CCC([C@H](O)c3ncc[nH]3)CC2)C1 |
| ZINC000809405032 | -52.50 | 129522 | 0.27 | 99.4% | NT | NT | NT | ND | Cc1ccc(N2CCN(C(=O)C(=O)N[C@@H]3CN(C4CC4)C[C@@H]3C)CC2)cc1 |
| ZINC000805335094 | -52.50 | 129517 | 0.32 | 133.3% | NT | NT | NT | ND | COc1ccc(C(=O)N2CC[C@H](N3CCCC3)[C@@H]2C)nc1Cl |
| ZINC000802379169 | -52.50 | 129515 | 0.27 | 89.7% | NT | NT | NT | ND | Cc1ccc(C2=NO[C@@H](CNC(=O)N3CC[C@H](N4CCCC4)C3)C2)cc1C |
| ZINC000684407814 | -52.50 | 129483 | 0.22 | 70.6% | NT | NT | NT | ND | CCN(CC)c1ncc(CN2CC(C(C)C)C2)s1 |
| ZINC000669616349 | -52.50 | 129471 | 0.28 | 56.6% | NT | NT | NT | ND | OC[C@H](N[C@H]1Cc2[nH]c3ccccc3c2C1)C1CCCCC1 |
| ZINC000064804934 | -52.50 | 129057 | 0.31 | 85.0% | NT | NT | NT | ND | CCCn1c(C)cc(/C=C(/C#N)C(=O)NCCN(C)Cc2ccccc2)c1C |
| ZINC000624110289 | -52.50 | 129425 | 0.16 | 49.3% | NT | NT | NT | ND | CC(C)CC[C@@H](CO)N[C@H](C)c1nc2c(s1)CCCC2 |
| ZINC000621202771 | -52.50 | 129422 | 0.30 | 61.4% | NT | NT | NT | ND | CCCN1CCC[C@H]1C(=O)NCc1cnccc1C(F)(F)F |
| ZINC000592403811 | -52.50 | 129403 | 0.35 | 152.7% | NT | NT | NT | ND | CC(C)Cc1ccc(C(=O)N2CCN([C@@H]3CCC[C@@H]3O)CC2)cc1 |
| ZINC000574318236 | -52.50 | 129394 | 0.32 | 100.4% | NT | NT | NT | ND | C[C@@H]1C[C@H](NC(=O)C(=O)N(C)Cc2c(Cl)cccc2Cl)CN1C1CC1 |
| ZINC000536865847 | -52.50 | 129364 | 0.27 | 36.3% | NT | NT | NT | ND | CCc1cc(N2CCC[C@@H](N[C@H](C)c3ccc(OC)c(Cl)c3)C2)ncn1 |
| ZINC000525245955 | -52.50 | 129351 | 0.27 | 93.6% | NT | NT | NT | ND | CC[C@@H]1CCC[C@H](NC(=O)[C@H](C)N2CCCCCC2)C1 |
| ZINC000520440291 | -52.50 | 129349 | 0.29 | 102.2% | NT | NT | NT | ND | COc1cc(Cl)c(C)cc1NC(=O)[C@H](C)N(C)Cc1sccc1C |
| ZINC000501775665 | -52.50 | 129338 | 0.31 | 20.4% | NT | NT | NT | ND | C[C@@H](c1ccc(F)c(F)c1)N(C)C(=O)CN1CC[C@@H](C)[C@@H](C)C1 |
| ZINC000453607577 | -52.50 | 129302 | 0.33 | 86.8% | NT | NT | NT | ND | Cc1ccc(C[C@H](CO)NCc2cccc(F)c2)cc1C |
| ZINC000446751851 | -52.50 | 129294 | 0.30 | 20.8% | NT | NT | NT | ND | O[C@@H](CN[C@@H](c1ccc(Br)cc1Cl)C1CC1)CN1CCCC1 |
| ZINC000436303801 | -52.50 | 129288 | 0.21 | 75.3% | NT | NT | NT | ND | CC(C)N1C[C@H](NC(=O)N[C@@H](C)c2ccc(Br)s2)[C@@H](C)C1 |
| ZINC000410783840 | -52.50 | 129267 | 0.28 | 103.4% | NT | NT | NT | ND | CN(Cc1cc(F)cc(F)c1)C(=O)C(=O)N1CC[C@H](N2CC=CC2)C1 |
| ZINC000376738332 | -52.50 | 129261 | 0.27 | 84.0% | NT | NT | NT | ND | CC[C@H](c1ccccc1)N1CCC(C#N)(c2ccccn2)CC1 |
| ZINC000375486080 | -52.50 | 129258 | 0.28 | 48.7% | NT | NT | NT | ND | C[C@@H](CO)N1CCCN([C@@H](C)C(=O)Nc2cccc(Cl)c2Cl)CC1 |
| ZINC000348635286 | -52.50 | 129235 | 0.29 | 104.7% | NT | NT | NT | ND | CN(C/C=C\c1ccc(F)c(F)c1)[C@@H]1COC[C@@H]1O |
| ZINC000295103757 | -52.50 | 129201 | 0.29 | 114.1% | NT | NT | NT | ND | CS[C@@H]1CCN(Cc2cnc(C)n2-c2ccccc2)C1 |
| ZINC000261552362 | -52.50 | 129156 | 0.33 | 56.1% | NT | NT | NT | ND | C[C@H]1CN/C(=N/Cc2cccc(OCc3ccccc3)c2)N1 |
| ZINC000172648663 | -52.50 | 129111 | 0.31 | 96.3% | NT | NT | NT | ND | O=C(Nc1ccc2ncsc2c1)C(=O)N1CCCN(C2CCCC2)CC1 |
| ZINC000170742095 | -52.50 | 129109 | 0.30 | 29.7% | NT | NT | NT | ND | FC(F)(F)c1ccc(-c2nc(CN3CC[C@H](N4CCCC4)C3)cs2)cc1 |
| ZINC001515357045 | -52.50 | 129783 | 0.22 | 104.3% | NT | NT | NT | ND | O=C(C[C@]1(O)CCCNC1)Nc1cn2cc(Br)ncc2n1 |
| ZINC001307701283 | -52.50 | 129771 | 0.27 | 62.5% | NT | NT | NT | ND | CC[C@H]1CCN(CC(=O)Nc2cc(Cl)ccc2C#N)[C@H]1C |
| ZINC000124982852 | -52.50 | 129080 | 0.32 | 89.7% | NT | NT | NT | ND | N#Cc1csc(CN2CCC[C@H]2Cc2ccccc2)c1 |
| ZINC001169307444 | -52.50 | 129688 | 0.33 | 21.2% | NT | NT | NT | ND | C[C@@H](C(=O)NCc1ccc(Cl)cc1)N1CCC(C(C)(C)C)CC1 |
| ZINC000875242850 | -50.00 | 513411 | 0.30 | 76.2% | NT | NT | NT | ND | C[C@H]1CCC[C@H](NC(=O)C(=O)N2CCC(N3CCN(C)CC3)CC2)CC1 |
| ZINC000080247921 | -50.00 | 511743 | 0.32 | 24.7% | NT | NT | NT | ND | Cc1noc(-c2cccnc2NC2CCC(N3CCCCC3)CC2)n1 |
| ZINC000762555244 | -50.00 | 513237 | 0.27 | 84.1% | NT | NT | NT | ND | Cc1nn(Cc2ccc(F)cc2)c(Cl)c1CNC[C@@]1(O)CCN(C)C1 |
| ZINC000073649189 | -50.00 | 511725 | 0.33 | 74.2% | NT | NT | NT | ND | CC[C@@H](C)NC(=O)[C@@H](C)N1CCC(c2c[nH]c3ccccc32)CC1 |
| ZINC000726467110 | -50.00 | 513206 | 0.30 | 79.5% | NT | NT | NT | ND | CCN1CCN(CCNC(=S)N[C@H](C)c2cccs2)CC1 |
| ZINC000685844120 | -50.00 | 513192 | 0.27 | 51.3% | NT | NT | NT | ND | Brc1ccccc1-n1cc(CN2CC[C@@H]3CCC[C@@H]32)nn1 |
| ZINC000683045155 | -50.00 | 513174 | 0.24 | 108.1% | NT | NT | NT | ND | CC[C@@]1(NCC(=O)Nc2ccc(C)cc2Br)CCOC1 |
| ZINC000657977970 | -50.00 | 513090 | 0.30 | 83.1% | NT | NT | NT | ND | C[C@H](Cn1cncn1)N[C@@H]1CCN(c2ccccc2C(F)(F)F)C1 |
| ZINC000646273308 | -50.00 | 513036 | 0.30 | 106.5% | NT | NT | NT | ND | C[C@@H](Cc1c(Cl)cccc1Cl)NC(=O)c1cn(C[C@@H]2CCCN2)nn1 |
| ZINC000631236280 | -50.00 | 512994 | 0.28 | 92.0% | NT | NT | NT | ND | COc1ccccc1C[C@@](C)(CO)NCc1ccc2c(c1)ncn2C |
| ZINC000608224751 | -50.00 | 512943 | 0.32 | 75.6% | NT | NT | NT | ND | CC(C)Cc1ccc(C(=O)Nc2ccc(OCCN3CCN(C)CC3)cc2)cc1 |
| ZINC000586751050 | -50.00 | 512900 | 0.28 | 98.2% | NT | NT | NT | ND | C[C@@H](C(=O)NC[C@H](C)N1CCCC1)c1c(Cl)cccc1Cl |
| ZINC000584233558 | -50.00 | 512897 | 0.34 | 22.7% | NT | NT | NT | ND | CC1(C)Cc2cc(C(=O)N3CCC[C@@H](N4CCCC4)CC3)ccc2O1 |
| ZINC000564110976 | -50.00 | 512837 | 0.33 | 78.2% | NT | NT | NT | ND | CC1CCC(CNC(=O)C(=O)NC2CCN([C@@H]3CCC[C@@H]3O)CC2)CC1 |
| ZINC000557493019 | -50.00 | 512815 | 0.34 | 113.3% | NT | NT | NT | ND | COCCN1CCC[C@@H]1CNC(=O)C(=O)N1CCC[C@@H](C(C)C)CC1 |
| ZINC000533411740 | -50.00 | 512762 | 0.34 | 79.7% | NT | NT | NT | ND | C[C@H](C(=O)Nc1ccc(F)cc1Cl)N1CC[C@H](N2CCCC2)C1 |
| ZINC000524392254 | -50.00 | 512733 | 0.33 | 93.9% | NT | NT | NT | ND | O=C(Nc1cnc2ccccc2c1)[C@]1(O)CCN(Cc2ccccc2)C1 |
| ZINC000504552433 | -50.00 | 512684 | 0.28 | 49.2% | NT | NT | NT | ND | COCCN1CCC(N[C@H]2COc3cc(F)c(Br)cc32)CC1 |
| ZINC000465804078 | -50.00 | 512591 | 0.33 | 67.7% | NT | NT | NT | ND | CCN(CC)[C@H]1CCN(C(=O)NCCCc2cccc(Cl)c2)C1 |
| ZINC000413145325 | -50.00 | 512422 | 0.30 | 62.9% | NT | NT | NT | ND | CC1CCN(C[C@H](O)CNc2nccc3ccc([N+](=O)[O-])cc32)CC1 |
| ZINC000359009035 | -50.00 | 512329 | 0.35 | 99.8% | NT | NT | NT | ND | CC(C)N1C[C@H](NC(=O)c2ccn(Cc3ccccc3)c2)[C@@H](C)C1 |
| ZINC000301468116 | -50.00 | 512208 | 0.29 | 88.5% | NT | NT | NT | ND | c1ccc(SC[C@@H]2CCN(c3cc(N4CCCCC4)ncn3)C2)cc1 |
| ZINC000289897491 | -50.00 | 512172 | 0.27 | 51.0% | NT | NT | NT | ND | C[C@@H]1COCC[C@@H]1NCCc1c[nH]c2cccc(Br)c12 |
| ZINC000286616311 | -50.00 | 512160 | 0.29 | 75.0% | NT | NT | NT | ND | COC(=O)c1cccc(S(=O)(=O)N2CC[C@@H](N3CCCC3)[C@@H]2C)c1 |
| ZINC000286502332 | -50.00 | 512159 | 0.33 | 139.1% | NT | NT | NT | ND | CNC(=O)CC[C@H]1CCCCN1Cc1cc(-c2ccccc2)on1 |
| ZINC000271284821 | -50.00 | 512095 | 0.28 | 28.3% | NT | NT | NT | ND | CCCN(C)C[C@H]1CCN([C@H](C)C(=O)c2c[nH]c3ncccc23)C1 |
| ZINC000265181994 | -50.00 | 512072 | 0.32 | 102.0% | NT | NT | NT | ND | COC1CCC(N(C)CC(=O)Nc2ccc3ccccc3c2)CC1 |
| ZINC000247147334 | -50.00 | 512040 | 0.27 | 103.8% | NT | NT | NT | ND | CN1CCC[C@@H]2CN(Cn3ccc(-c4ccc(Br)cc4)n3)CC[C@H]21 |
| ZINC000192810020 | -50.00 | 512009 | 0.30 | 89.0% | NT | NT | NT | ND | COc1ccc2ccccc2c1-c1cc(CN2CCN(C[C@@H](C)O)C[C@H]2C)on1 |
| ZINC000155998528 | -50.00 | 511894 | 0.35 | 30.9% | NT | NT | NT | ND | CS[C@@H]1CCC[C@H](NC(=O)NC2CCN([C@@H]3CCCC[C@@H]3O)CC2)C1 |
| ZINC000155112667 | -50.00 | 511889 | 0.31 | 33.8% | NT | NT | NT | ND | C[C@@H](CN1CCN(C)CC1)N[C@@H](C)c1ccc(-c2ccccc2)o1 |
| ZINC000153400973 | -50.00 | 511882 | 0.30 | 99.4% | NT | NT | NT | ND | O=C(CN1CC[C@]2(CCOC2)C1)N[C@@H](Cc1ccccc1)c1ccccc1 |
| ZINC001418905447 | -50.00 | 514167 | 0.26 | 129.5% | NT | NT | NT | ND | CC(C)N1CC[C@H](CCNC(=O)c2cc(-c3cccnc3)on2)C1 |
| ZINC000135482356 | -50.00 | 511863 | 0.34 | 21.7% | NT | NT | NT | ND | O=C(CCc1cc(Br)ccc1F)N1CC[C@@H](CN2CCCC2)C1 |
| ZINC001033973784 | -50.00 | 513673 | 0.26 | 43.3% | NT | NT | NT | ND | CCN(C(=O)c1ccnc(C2CC2)n1)[C@H]1CCN(CC2CCC2)C1 |
| ZINC000981787500 | -47.50 | 1644796 | 0.32 | 48.4% | NT | NT | NT | ND | O=C(C1CC2(CCC2)C1)N1CCCN(C[C@@H](O)c2ccccc2)CC1 |
| ZINC000972168981 | -47.50 | 1644771 | 0.22 | 63.4% | NT | NT | NT | ND | CN(C[C@H]1CCCCO1)[C@@H]1CCN(C(=O)c2nc(Cl)cs2)C1 |
| ZINC000866213504 | -47.50 | 1644195 | 0.32 | 204.0% | NT | NT | NT | ND | CN1CCN(C2CCN(c3nc(N)c4ccccc4n3)CC2)CC1 |
| ZINC000664479370 | -47.50 | 1643543 | 0.27 | 103.0% | NT | NT | NT | ND | O=C(N[C@H]1CCOC2(CCC2)C1)N1CCC(N2CC[C@@H](O)C2)CC1 |
| ZINC000575186982 | -47.50 | 1642833 | 0.30 | 22.9% | NT | NT | NT | ND | Cc1ccc2c(c1)[C@H](NC(=O)NC[C@@H]1CCN(C(C)C)C1)CCC2 |
| ZINC000556484619 | -47.50 | 1642692 | 0.29 | 111.9% | NT | NT | NT | ND | CC(C)OCCN1CCC(NC(=O)[C@@H]2CCc3nncn3C2)CC1 |
| ZINC000539406733 | -47.50 | 1642577 | 0.34 | 34.0% | NT | NT | NT | ND | CCCC1CCC(CN2CCN(CC(=O)N(CC)CC)CC2)CC1 |
| ZINC000529649527 | -47.50 | 1642503 | 0.33 | 84.3% | NT | NT | NT | ND | C[C@@H](C(=O)Nc1cccc(C2CCCCC2)c1)N1CCN(C)CC1 |
| ZINC000496403719 | -47.50 | 1642306 | 0.27 | 92.4% | NT | NT | NT | ND | CN(C[C@H](O)CN[C@@H]1C[C@H]2CCCCN2C1=O)C(=O)OC(C)(C)C |
| ZINC000454705882 | -47.50 | 1642082 | 0.32 | 87.6% | NT | NT | NT | ND | CN1CC[C@@H](CNC(=O)Nc2cc(F)ccc2Oc2ccccc2)C1 |
| ZINC000452100574 | -47.50 | 1642051 | 0.21 | 73.3% | NT | NT | NT | ND | CC(C)N1CC[C@H](N(C)S(=O)(=O)C[C@H]2CCCCC2(F)F)C1 |
| ZINC000450551910 | -47.50 | 1642031 | 0.33 | 16.0% | NT | NT | NT | ND | C[C@@H]1CCC[C@H](NC(=O)CN2CCN(CCC3=CCCCC3)CC2)C1 |
| ZINC000417806012 | -47.50 | 1641789 | 0.29 | 96.2% | NT | NT | NT | ND | CC(C)(C(=O)N[C@@H]1CCCc2cc(N)ccc21)[C@H](N)c1ccccc1 |
| ZINC000416493836 | -47.50 | 1641780 | 0.31 | 72.3% | NT | NT | NT | ND | Cc1cc([N+](=O)[O-])cc(C)c1S(=O)(=O)NC[C@H]1CCN(C)C1 |
| ZINC000370933962 | -47.50 | 1641610 | 0.28 | 98.9% | NT | NT | NT | ND | CN1CCC(C(=O)N2CCC[C@@H](c3nc4ccc(F)cc4o3)C2)CC1 |
| ZINC000370893072 | -47.50 | 1641608 | 0.34 | 82.1% | NT | NT | NT | ND | O=C1CCCCN1[C@@H]1CCCN(Cc2cc(-c3ccccc3)no2)C1 |
| ZINC000367601434 | -47.50 | 1641578 | 0.27 | 90.8% | NT | NT | NT | ND | Cc1ccccc1Cc1nnc(CN2C[C@@H](N3CCCC3)C[C@H]2C)o1 |
| ZINC000362611503 | -47.50 | 1641523 | 0.31 | 27.8% | NT | NT | NT | ND | CCO[C@@H](CC(=O)N1CCCN(C2CCCC2)CC1)C1=CCCC1 |
| ZINC000285662957 | -47.50 | 1640983 | 0.29 | 17.5% | NT | NT | NT | ND | COc1ccc(-c2ccccc2)cc1CNCCC1(O)CCOCC1 |
| ZINC000272631760 | -47.50 | 1640875 | 0.24 | 91.2% | NT | NT | NT | ND | CN1CCC(C)(CNC(=O)Nc2ccc(Br)nc2)CC1 |
| ZINC000272592294 | -47.50 | 1640872 | 0.27 | 79.1% | NT | NT | NT | ND | Cc1c(Br)cccc1C(=O)N1CC[C@@H]2CCN(C)C[C@H]21 |
| ZINC000265281429 | -47.50 | 1640797 | 0.34 | 97.5% | NT | NT | NT | ND | Cc1cccnc1[C@H](NC(=O)N1CCN(C2CCC2)CC1)C(C)C |
| ZINC000237909071 | -47.50 | 1640685 | 0.33 | 89.0% | NT | NT | NT | ND | N[C@@H](CC(=O)Nc1cccc(C(=O)N2CCCC2)c1)c1ccccc1 |
| ZINC000194878845 | -47.50 | 1640634 | 0.35 | 109.1% | NT | NT | NT | ND | CC(C)(C)N1CCC(NC(=O)c2ncccc2Br)CC1 |
| ZINC000159130997 | -47.50 | 1640353 | 0.30 | 107.9% | NT | NT | NT | ND | CC(C)[C@H](C(=O)NC[C@H](C[C@@H](C)O)c1ccccc1)N1CCCCC1 |
| ZINC000153451054 | -47.50 | 1640315 | 0.29 | 17.2% | NT | NT | NT | ND | Cc1cc(C(C)(C)C)ccc1OC[C@@H](O)CN1CC[C@]2(CCOC2)C1 |
| ZINC001403624658 | -47.50 | 1646408 | 0.30 | 65.3% | NT | NT | NT | ND | COc1ccc(C(=O)N(C)C[C@H](C)NCC2CCC(C)(C)CC2)nn1 |
| ZINC001399642198 | -47.50 | 1646401 | 0.22 | 21.2% | NT | NT | NT | ND | C[C@@H]1CCC[C@H](CCN(C)CCNC(=O)c2c3c(nn2C)CCC3)C1 |
| ZINC001391502525 | -47.50 | 1646389 | 0.32 | 43.6% | NT | NT | NT | ND | O=C(NCC1CN(CCC2CCCC2)C1)C1(c2cccnc2)CC1 |
| ZINC001381212829 | -47.50 | 1646382 | 0.23 | 71.1% | NT | NT | NT | ND | Cc1c(C(=O)NCC[C@H](C)NCc2nccc(C)n2)ccn1C(C)C |
| ZINC000130668661 | -47.50 | 1640271 | 0.34 | 68.6% | NT | NT | NT | ND | CN1CCN(c2ncc(CN3CCN(CC4CC4)CC3)cn2)CC1 |
| ZINC001196417557 | -47.50 | 1645787 | 0.30 | 71.9% | NT | NT | NT | ND | CCc1ccc(F)cc1C(=O)N1CCCN(CCOC(C)C)CC1 |
| ZINC001099969021 | -47.50 | 1645370 | 0.30 | 28.5% | NT | NT | NT | ND | O=C(N[C@@H]1CCN(CCC2CCCC2)C[C@@H]1O)C1(C(F)F)CCC1 |
| ZINC001077034834 | -47.50 | 1645249 | 0.28 | 16.5% | NT | NT | NT | ND | Cc1c(F)cccc1C(=O)N[C@@H]1CN(CCC2CCCCC2)C[C@H]1O |
| ZINC001061620290 | -47.50 | 1645213 | 0.25 | 91.3% | NT | NT | NT | ND | CN1CCC(C(=O)NC[C@@H]2CCN(c3ncc(Cl)cn3)C2)CC1 |
| ZINC001061317651 | -47.50 | 1645212 | 0.26 | 62.6% | NT | NT | NT | ND | CN1CCC[C@@H]1C(=O)NC[C@H]1CCCN1c1ncnc2scnc21 |
| ZINC001040612984 | -47.50 | 1645141 | 0.31 | 100.0% | NT | NT | NT | ND | COc1ccnc(CN2CCC3(CCN(C(=O)C(C)C)C3)CC2)c1 |
| ZINC001033739722 | -47.50 | 1645092 | 0.29 | 24.6% | NT | NT | NT | ND | CCN(C(=O)c1cnccn1)[C@@H]1CCN(CC2CCCCCC2)C1 |
| ZINC001032169623 | -47.50 | 1645059 | 0.28 | 92.8% | NT | NT | NT | ND | Cc1ccnc2nc(C(=O)NCC3CN(CCCC4CCC4)C3)nn21 |
| ZINC001018723708 | -47.50 | 1644923 | 0.30 | 101.3% | NT | NT | NT | ND | Cc1cc(C(=O)N2CC[C@H](NCc3cccc(Cl)c3F)C2)ncn1 |
| ZINC001004070100 | -47.50 | 1644876 | 0.31 | 85.5% | NT | NT | NT | ND | CC[C@@H](C(N)=O)N1CCC(NC(=O)[C@H]2Cc3ccc(Cl)cc32)CC1 |
| ZINC000971070823 | -45.00 | 4281957 | 0.28 | 90.9% | NT | NT | NT | ND | Cc1cc(C(=O)N2CC[C@H](N(C)Cc3ccc(C)c(C)c3)C2)on1 |
| ZINC000929842198 | -45.00 | 4281540 | 0.23 | 103.9% | NT | NT | NT | ND | O=C1CCCCN1C1CCN(C[C@H](O)C[C@]2(O)CCOC2)CC1 |
| ZINC000897504395 | -45.00 | 4281106 | 0.28 | 90.9% | NT | NT | NT | ND | C[C@@H]1CCCN1CCNC(=O)N1CCC[C@@H]1[C@@H]1CCCC[C@@]1(C)O |
| ZINC000880249690 | -45.00 | 4280844 | 0.35 | 113.4% | NT | NT | NT | ND | NC(=O)C1(N2CCCCC2)CCN(CC(=O)NCC2CCC2)CC1 |
| ZINC000774469834 | -45.00 | 4279905 | 0.31 | 83.3% | NT | NT | NT | ND | O=C(NC1CCCCC1)C(=O)NC1CCN(C[C@H]2CCCO2)CC1 |
| ZINC000679555790 | -45.00 | 4279567 | 0.28 | 103.1% | NT | NT | NT | ND | COCCOCc1cccc(NC(=O)NCC2(C)CCN(C)CC2)c1 |
| ZINC000673584800 | -45.00 | 4279488 | 0.34 | 110.1% | NT | NT | NT | ND | O=C(NCCCC1CC1)C(=O)NC1CCN([C@H]2CCC[C@@H]2O)CC1 |
| ZINC000667676804 | -45.00 | 4279359 | 0.23 | 94.4% | NT | NT | NT | ND | O=C(Nc1cnn(CC2CC2)c1)C(=O)N1CCC[C@H]([C@@H]2CCNC2)C1 |
| ZINC000655168559 | -45.00 | 4278968 | 0.32 | 97.6% | NT | NT | NT | ND | O=C(Nc1ccc2ncnn2c1)[C@]1(O)CCN(Cc2ccccc2)C1 |
| ZINC000623468108 | -45.00 | 4278334 | 0.27 | 84.9% | NT | NT | NT | ND | C[C@@H]1CCCC[C@H]1OCCNC(=O)NC[C@H]1CCCCN(C)C1 |
| ZINC000603324806 | -45.00 | 4278096 | 0.25 | 65.5% | NT | NT | NT | ND | CCc1c(C(=O)N[C@@H](C)C2CCN(C)CC2)cnn1-c1ccccc1 |
| ZINC000588863575 | -45.00 | 4277895 | 0.30 | 133.8% | NT | NT | NT | ND | O=C(Cn1cc([N+](=O)[O-])ccc1=O)N1CC[C@H](N2CC[C@@H](O)C2)C1 |
| ZINC000495656270 | -45.00 | 4276787 | 0.31 | 45.1% | NT | NT | NT | ND | CCCCNC(=O)[C@H](C)N1CC[C@](C)(CNC(=O)OC(C)(C)C)C1 |
| ZINC000453336584 | -45.00 | 4276316 | 0.28 | 100.2% | NT | NT | NT | ND | C[C@@H](CC(N)=O)NC[C@H](CO)Cc1cc(Br)ccc1F |
| ZINC000415549177 | -45.00 | 4275670 | 0.24 | 73.8% | NT | NT | NT | ND | O=C(NC[C@H]1CCCN1)Nc1cc(F)cc(F)c1Br |
| ZINC000358071102 | -45.00 | 4275056 | 0.28 | 112.6% | NT | NT | NT | ND | CCc1ccnc(CNC(=O)C(=O)N[C@@H]2C[C@H](C)N(C3CC3)C2)c1 |
| ZINC000358051108 | -45.00 | 4275054 | 0.30 | 102.3% | NT | NT | NT | ND | CN1CC[C@](O)(CNC(=O)c2cc3cc([N+](=O)[O-])ccc3s2)C1 |
| ZINC000345970626 | -45.00 | 4274806 | 0.29 | 71.7% | NT | NT | NT | ND | CC[C@H](NC(=O)C(=O)N(CC)CCCN1CCCC1)[C@@H]1CCCO1 |
| ZINC000340106708 | -45.00 | 4274656 | 0.32 | 50.0% | NT | NT | NT | ND | Cc1ccncc1N1CCN(C[C@H](O)CN2C[C@@H](C)O[C@H](C)C2)CC1 |
| ZINC000339640280 | -45.00 | 4274642 | 0.32 | 105.8% | NT | NT | NT | ND | C[C@H]1CN(CCN(C)C(=O)C(=O)NC2CCN(C)CC2)C[C@H](C)O1 |
| ZINC000334024070 | -45.00 | 4274521 | 0.29 | 110.7% | NT | NT | NT | ND | C[C@H](C(=O)NCC1CCCCC1)N1CCC[C@H](N2CCCC2=O)CC1 |
| ZINC000298791523 | -45.00 | 4274320 | 0.31 | 89.0% | NT | NT | NT | ND | O=C(CN1CCC(c2nc3cc(Cl)ccc3o2)CC1)N1CCC1 |
| ZINC000188412270 | -45.00 | 4273358 | 0.28 | 106.5% | NT | NT | NT | ND | O=C(NCCc1ccc(F)c(Br)c1)[C@H]1C[C@H](F)CN1 |
| ZINC001437918206 | -45.00 | 4286062 | 0.26 | 102.6% | NT | NT | NT | ND | Cc1cnc(CN2CC[C@@H](CCNC(=O)c3cncs3)C2)s1 |
| ZINC001419404744 | -45.00 | 4286029 | 0.34 | 19.6% | NT | NT | NT | ND | Cc1cc(C)nc(C(=O)NCCC2CCN(CCOC(C)C)CC2)c1 |
| ZINC001377717241 | -45.00 | 4285904 | 0.27 | 101.5% | NT | NT | NT | ND | CCC1(C(=O)NCC[C@@H]2CCN(Cc3nccn3C)C2)CCCC1 |
| ZINC000136880129 | -45.00 | 4272868 | 0.25 | 53.9% | NT | NT | NT | ND | O=C(NCc1ccc(F)cc1C(F)(F)F)[C@@H]1CC12CCNCC2 |
| ZINC001149399826 | -45.00 | 4284075 | 0.27 | 96.4% | NT | NT | NT | ND | CN1CCC[C@@H]1C(=O)NC[C@H]1CN(CCCC(C)(C)C)CCCO1 |
| ZINC001048683995 | -45.00 | 4283155 | 0.28 | 61.7% | NT | NT | NT | ND | Cc1ccc(CN2C[C@@H]3CN(C(=O)c4cccn4C)C[C@@H]3C2)cc1C |
| ZINC001019309159 | -45.00 | 4282459 | 0.30 | 99.6% | NT | NT | NT | ND | O=C([C@H]1Cc2ccccc21)N1CC[C@H](NCc2ccncc2Cl)C1 |
| ZINC001002178741 | -45.00 | 4282269 | 0.35 | 109.0% | NT | NT | NT | ND | Cc1cccc(F)c1C(=O)NCC1CCN(CCn2cncn2)CC1 |
| ZINC000973375109 | -42.50 | 9184026 | 0.31 | 110.9% | NT | NT | NT | ND | C[C@@H]1CCN(CC(=O)N[C@H]2C[C@H](NC(=O)c3cnnc(O)c3)C2)C1 |
| ZINC000950078544 | -42.50 | 9183476 | 0.23 | 108.0% | NT | NT | NT | ND | Cc1cc(C)nc(C(=O)N2CC[C@H]2CNC(=O)[C@@H]2CCN(C)C2)c1 |
| ZINC000903470913 | -42.50 | 9182545 | 0.30 | 92.2% | NT | NT | NT | ND | O=C(CN[C@H]1CCNC12CCC2)NCCN1CCc2ccccc21 |
| ZINC000847525041 | -42.50 | 9181198 | 0.27 | 59.3% | NT | NT | NT | ND | CN1CC[C@]2(CCN(C(=O)C(=O)Nc3cccc4nonc43)C2)C1 |
| ZINC000805503179 | -42.50 | 9180643 | 0.23 | 105.2% | NT | NT | NT | ND | CN1CCN(CCNC(=O)C(=O)NC[C@H]2CCC=CO2)CC1(C)C |
| ZINC000804054066 | -42.50 | 9180634 | 0.27 | 90.0% | NT | NT | NT | ND | O=C(NC[C@@H]1CN2CCC[C@@H]2CO1)C(=O)N[C@@H]1CC[C@H]2CCC[C@@H]2C1 |
| ZINC000738383044 | -42.50 | 9180101 | 0.25 | 117.6% | NT | NT | NT | ND | O[C@@H](CN1CCN(c2ccc(-c3nn[nH]n3)cc2F)CC1)C1CC1 |
| ZINC000595353335 | -42.50 | 9177460 | 0.32 | 97.7% | NT | NT | NT | ND | CCCN1CC[C@H](NS(=O)(=O)c2ccc(C(=O)OC)cc2F)C1 |
| ZINC000576277949 | -42.50 | 9177091 | 0.26 | 67.2% | NT | NT | NT | ND | C[C@H](NC1CC1)C(=O)NCc1ccc(C(=O)N=c2cc[nH]cc2)cc1 |
| ZINC000556797005 | -42.50 | 9176687 | 0.31 | 79.9% | NT | NT | NT | ND | CN1CCN(C(=O)C(=O)N2CCCN(C3CCCC3)CC2)CC1=O |
| ZINC000555574173 | -42.50 | 9176659 | 0.33 | 106.6% | NT | NT | NT | ND | CN1CCN(CCCCNC(=O)C(=O)N2CCSCC2)CC1 |
| ZINC000438095302 | -42.50 | 9174651 | 0.27 | 38.1% | NT | NT | NT | ND | C[C@@H]1CCC[C@@H](NC(=O)CN2CCN(CC[C@@H]3CCOC3)CC2)C1 |
| ZINC000418924570 | -42.50 | 9174087 | 0.24 | 64.9% | NT | NT | NT | ND | Cc1cccc(C)c1OC[C@H](C)NC(=O)C(=O)NC[C@@H]1CCCNC1 |
| ZINC000377772869 | -42.50 | 9173616 | 0.31 | 100.1% | NT | NT | NT | ND | CS[C@H](CO)[C@@H](C)NC(=O)N[C@@H]1CCN(C2CCN(C)CC2)C1 |
| ZINC000363920872 | -42.50 | 9173179 | 0.28 | 88.9% | NT | NT | NT | ND | CC(C)N1CCN([C@H]2CCN(C(=O)CC3(O)CCCCC3)C2)CC1 |
| ZINC000283462313 | -42.50 | 9171287 | 0.35 | 98.5% | NT | NT | NT | ND | Cc1noc(C)c1CNC(=O)C(=O)N1CCN(C2CCCC2)CC1 |
| ZINC000276781830 | -42.50 | 9171131 | 0.26 | 97.6% | NT | NT | NT | ND | CC[C@@]1(C)Oc2ccc(NC(=O)C(=O)NC[C@@H]3CCN(C)C3)cc2O1 |
| ZINC000270395954 | -42.50 | 9170979 | 0.26 | 120.7% | NT | NT | NT | ND | COCC1(O)CCN(C[C@@H](O)Cn2cnc3cc(C)c(C)cc32)CC1 |
| ZINC000182663590 | -42.50 | 9170002 | 0.25 | 98.0% | NT | NT | NT | ND | C[C@H](CCNC(=O)N1CCO[C@H]([C@@H]2CCCO2)C1)N1CCCCC1 |
| ZINC000178638284 | -42.50 | 9169879 | 0.25 | 111.1% | NT | NT | NT | ND | O=C1Nc2ccccc2C12CCC(NCC1(O)CCOCC1)CC2 |
| ZINC000152090354 | -42.50 | 9169412 | 0.29 | 44.7% | NT | NT | NT | ND | COC(=O)[C@@H](C)CS(=O)(=O)N[C@@H]1CCN(CC2CCCC2)C1 |
| ZINC001072835483 | -42.50 | 9186763 | 0.20 | 81.8% | NT | NT | NT | ND | CNC(=O)[C@@H](C)N1CCC2(CN(C(=O)c3c[nH]c4nccc-4c3)C2)C1 |
| ZINC001055479120 | -42.50 | 9186689 | 0.32 | 98.5% | NT | NT | NT | ND | CC(=O)NCCN1CCC(NC(=O)c2c(C)[nH]nc2Cl)CC1 |
| ZINC001041605786 | -42.50 | 9186302 | 0.24 | 109.7% | NT | NT | NT | ND | O=C([C@@H]1CCCN(CC(F)F)C1)N1CC[C@]2(CCN(CCO)C2)C1 |
| ZINC001021041334 | -42.50 | 9185186 | 0.30 | 86.2% | NT | NT | NT | ND | Cc1ccccc1CN[C@H]1C[C@H](NC(=O)c2cn(C)c(=O)nc2O)C1 |
| ZINC001006004942 | -42.50 | 9184781 | 0.21 | 125.6% | NT | NT | NT | ND | C[C@@H](NC(=O)c1cnnn1C)C1CN(C(=O)C23CCCN2CCC3)C1 |
| ZINC001003018745 | -42.50 | 9184684 | 0.25 | 85.7% | NT | NT | NT | ND | CCc1nnsc1C(=O)NCC1CN(C(=O)[C@H]2CCCN2C)C1 |
| ZINC001000730978 | -42.50 | 9184598 | 0.32 | 81.3% | NT | NT | NT | ND | O=C(NCC1CN(C(=O)[C@H]2CC[C@H]3CCCCN32)C1)c1cnccn1 |
| ZINC000995014137 | -40.00 | 16745329 | 0.28 | 122.9% | NT | NT | NT | ND | Cc1c(C(=O)NC2CN(C(=O)c3cnns3)C2)ccc2cncn21 |
| ZINC000991393621 | -40.00 | 16745251 | 0.27 | 116.1% | NT | NT | NT | ND | CCc1noc(C)c1C(=O)NC1CN(C(=O)[C@H]2CCCCN2C)C1 |
| ZINC000955102158 | -40.00 | 16743774 | 0.25 | 111.3% | NT | NT | NT | ND | CN(C(=O)c1cccnn1)C1CN(C(=O)[C@H]2CCC[C@H]3CCCN32)C1 |
| ZINC000937267298 | -40.00 | 16743375 | 0.24 | 111.1% | NT | NT | NT | ND | C[C@H]1CCCN1CC(=O)N(C)[C@H]1CCN(C(=O)CC(C)(C)O)C1 |
| ZINC000813849560 | -40.00 | 16739916 | 0.28 | 108.4% | NT | NT | NT | ND | C[C@H](NC(=O)c1ccc2[nH]c(=O)c(=O)[nH]c2c1)[C@H](O)c1ccccc1 |
| ZINC000679371269 | -40.00 | 16738743 | 0.22 | 114.4% | NT | NT | NT | ND | CN1CC[C@H]2CCN(C(=O)C(=O)Nc3cnc4c(c3)COCC4)[C@@H]2C1 |
| ZINC000671981913 | -40.00 | 16738552 | 0.28 | 105.7% | NT | NT | NT | ND | C[C@@H](NC(=O)C(=O)NCCCN1CCC[C@H](C(N)=O)C1)C1CCC1 |
| ZINC000627254962 | -40.00 | 16736329 | 0.28 | 113.5% | NT | NT | NT | ND | O=C([C@@H]1CCCN1c1ncccn1)N1CC[C@@H](N2CC[C@H](O)C2)C1 |
| ZINC000045463977 | -40.00 | 16722778 | 0.32 | 107.0% | NT | NT | NT | ND | CC(C)CN1CCC(NC(=O)NNC(=O)C(=O)NC2CC2)CC1 |
| ZINC000419873354 | -40.00 | 16730953 | 0.27 | 95.7% | NT | NT | NT | ND | CC[C@H]1CN(C(=O)C(=O)NCCCN2CCNCC2)CCS1 |
| ZINC000358062278 | -40.00 | 16729240 | 0.28 | 101.7% | NT | NT | NT | ND | CN1CC[C@](O)(CNC(=O)c2ccc([N+](=O)[O-])c3cccnc23)C1 |
| ZINC000297448741 | -40.00 | 16727579 | 0.26 | 104.8% | NT | NT | NT | ND | CN(CCCN(C)C(=O)C(=O)N(CCO)C1CC1)Cc1ccco1 |
| ZINC000287105318 | -40.00 | 16727190 | 0.23 | 117.3% | NT | NT | NT | ND | Cc1c(CNC(=O)C(=O)N2CC[C@@H](C3CCN(C)CC3)C2)cnn1C |
| ZINC000279187431 | -40.00 | 16726935 | 0.23 | 104.9% | NT | NT | NT | ND | COCCOc1ccc(NC(=O)C(=O)N2C[C@H]3CN(C)C[C@H]3C2)cn1 |
| ZINC000158577469 | -40.00 | 16724695 | 0.22 | 111.1% | NT | NT | NT | ND | C[C@H]1C[C@@H](NC(=O)C(=O)Nc2cnn(CC(F)F)c2)CN1C1CC1 |
| ZINC001365986475 | -40.00 | 16755723 | 0.33 | 108.9% | NT | NT | NT | ND | COC[C@@H](O)CN1CC[C@H](NC(=O)c2c(C)c(C)nn(C)c2=O)C1 |
| ZINC000133744458 | -40.00 | 16724378 | 0.26 | 65.7% | NT | NT | NT | ND | CN(C(=O)C[C@@H]1CSCCN1)[C@@H]1CCCN(c2cccnn2)C1 |
| ZINC001073859800 | -40.00 | 16749059 | 0.33 | 110.4% | NT | NT | NT | ND | CCc1cc(C(=O)NC[C@H]2CN(CCn3cncn3)CCCO2)no1 |
| ZINC001055138151 | -40.00 | 16748861 | 0.22 | 102.8% | NT | NT | NT | ND | Cc1cnc(C(=O)N2C[C@H]3CN(C(=O)CN(C)C(C)C)C[C@H]3C2)cn1 |
| ZINC001054647418 | -40.00 | 16748846 | 0.31 | 104.4% | NT | NT | NT | ND | COCCn1cc(C(=O)N2C[C@@H](C)[C@@H](NCc3ccccn3)C2)nn1 |
| ZINC001039123057 | -40.00 | 16748099 | 0.23 | 114.6% | NT | NT | NT | ND | C[C@@H](O)CN1CC[C@H](C2CCN(C(=O)CN3CCOCC3)CC2)C1 |
| ZINC001028614115 | -40.00 | 16747245 | 0.24 | 115.3% | NT | NT | NT | ND | Cc1nc(O)[nH]c1C(=O)NC[C@H]1CCN([C@H](C)C(=O)NC2CC2)C1 |
| ZINC001028403773 | -40.00 | 16747183 | 0.28 | 109.0% | NT | NT | NT | ND | Cc1cc(C(=O)NC[C@@H]2CCN(CC(=O)Nc3cnccn3)C2)on1 |
| ZINC001018665695 | -40.00 | 16746246 | 0.24 | 120.0% | NT | NT | NT | ND | Cn1ccc(N2CC[C@@H](N[C@H]3CCN(C(=O)CC4CCC4)C3)C2=O)n1 |
| ZINC001002391793 | -40.00 | 16745751 | 0.29 | 90.1% | NT | NT | NT | ND | CCN1CC[C@@H](N2CCC(NC(=O)c3cncnc3C)CC2)C1=O |
| ZINC001001445660 | -40.00 | 16745711 | 0.24 | 92.1% | NT | NT | NT | ND | O=C(NCC1CCN([C@H]2CCNC2=O)CC1)c1ccn(C(F)F)n1 |
| ZINC000997292482 | -37.50 | 27058057 | 0.35 | 87.9% | NT | NT | NT | ND | Cc1n[nH]cc1C(=O)NC1CN(C(=O)[C@H]2C[C@@H]3CCCCN3C2)C1 |
| ZINC000982403760 | -37.50 | 27057145 | 0.26 | 104.2% | NT | NT | NT | ND | CC(C)=CC(=O)N1CC[C@@H](CNC(=O)c2ccn(CCN(C)C)n2)C1 |
| ZINC000936344871 | -37.50 | 27055230 | 0.26 | 82.7% | NT | NT | NT | ND | C[C@H]1CCCN1CC(=O)N(C)[C@H]1CCN(C(=O)C2CCOCC2)C1 |
| ZINC000891830660 | -37.50 | 27053322 | 0.28 | 116.4% | NT | NT | NT | ND | CO[C@@H]1CN(C(=O)NCc2ccnc(N3CCN(C)CC3)c2)CCO1 |
| ZINC000863492725 | -37.50 | 27052173 | 0.25 | 178.7% | NT | NT | NT | ND | CCC1(N2CCOCC2)CCN(C(=O)C2=NN(C)C(=O)CC2)CC1 |
| ZINC000806327364 | -37.50 | 27050726 | 0.30 | 105.9% | NT | NT | NT | ND | CC[C@@H](CN1CCCC1)NC(=O)C(=O)NC1COC(C)(C)OC1 |
| ZINC000518759550 | -37.50 | 27042471 | 0.31 | 114.6% | NT | NT | NT | ND | Cc1ccnc2nc(C(=O)N3CCN(CCN4CCCC4)CC3)nn21 |
| ZINC000444807044 | -37.50 | 27040276 | 0.20 | 120.2% | NT | NT | NT | ND | CCO[C@@H]1C[C@](CO)(NC[C@H](O)COC2CCOCC2)C1(C)C |
| ZINC000438905691 | -37.50 | 27040072 | 0.25 | 105.4% | NT | NT | NT | ND | Cc1c(C(=O)NCC2CCCC2)cnc2c1c(=O)n(C)c(=O)n2C |
| ZINC000419445843 | -37.50 | 27039056 | 0.30 | 82.5% | NT | NT | NT | ND | CN(Cc1ccccc1)C(=O)CNC(=O)C(=O)N[C@H]1CCCNC1 |
| ZINC000378210510 | -37.50 | 27037950 | 0.33 | 95.3% | NT | NT | NT | ND | NC(=O)c1ccccc1N1CC[C@H](NC(=O)CC2CCNCC2)C1 |
| ZINC000366544782 | -37.50 | 27037378 | 0.22 | 110.5% | NT | NT | NT | ND | CN1CC[C@]2(CCN(C(=O)CN3C(=O)NC4(CCCC4)C3=O)C2)C1 |
| ZINC000349353698 | -37.50 | 27036436 | 0.33 | 142.4% | NT | NT | NT | ND | C[C@@H]1CCN2C(=O)N(CCCN3CCC[C@@H](C(N)=O)C3)C(=O)[C@H]2C1 |
| ZINC000343016571 | -37.50 | 27036040 | 0.33 | 117.8% | NT | NT | NT | ND | COCCN1CCC(NC(=O)C(=O)N[C@H]2CCOC(C)(C)C2)CC1 |
| ZINC000286782676 | -37.50 | 27034389 | 0.22 | 132.4% | NT | NT | NT | ND | CCc1nn(C)c(Cl)c1C(=O)N1CC[C@@H](N2CCNCC2)C1 |
| ZINC001416718427 | -37.50 | 27072472 | 0.26 | 80.5% | NT | NT | NT | ND | CCN1CCCC[C@H]1C(=O)NC[C@@H](O)CNC(=O)c1cnccn1 |
| ZINC001369886058 | -37.50 | 27071520 | 0.29 | 111.3% | NT | NT | NT | ND | O=C(CCn1cccn1)NCCN1CCN(CCn2cccn2)CC1 |
| ZINC001088099598 | -37.50 | 27063740 | 0.24 | 132.4% | NT | NT | NT | ND | Cc1ccoc1CC(=O)N1C[C@H]2CCN(CC(=O)N(C)C)C[C@H]2C1 |
| ZINC001074556982 | -37.50 | 27062813 | 0.22 | 89.6% | NT | NT | NT | ND | O=C(C[C@H]1C=CCC1)NCC1(O)CCN(C(=O)c2cncnc2)CC1 |
| ZINC001073886256 | -37.50 | 27062797 | 0.30 | 112.7% | NT | NT | NT | ND | CN1CC[C@@H](C(=O)NC[C@H]2CN(Cc3csnn3)CCCO2)C1 |
| ZINC001052175095 | -37.50 | 27062336 | 0.30 | 103.8% | NT | NT | NT | ND | CCN1CC[C@H](N2CCC[C@@H](NC(=O)c3ccc(O)nn3)CC2)C1=O |
| ZINC001051553588 | -37.50 | 27062271 | 0.26 | 104.8% | NT | NT | NT | ND | Cc1ncc(CNC[C@@H]2CN(C(=O)c3cn(C)cn3)CCO2)cn1 |
| ZINC001042098162 | -37.50 | 27061610 | 0.25 | 101.6% | NT | NT | NT | ND | CN1CCC[C@H]1C(=O)NCC1(O)CN(C(=O)CCc2ncccn2)C1 |
| ZINC001023604137 | -37.50 | 27059743 | 0.24 | 89.3% | NT | NT | NT | ND | C[C@@H](C(N)=O)N1CCC[C@H](CNC(=O)c2nn(C)c3c2CCCC3)C1 |
| ZINC001021109191 | -37.50 | 27059409 | 0.28 | 105.9% | NT | NT | NT | ND | Cn1cc(CN[C@H]2C[C@H](NC(=O)c3cccc(C(N)=O)n3)C2)cn1 |
| ZINC001019525589 | -37.50 | 27059164 | 0.28 | 85.0% | NT | NT | NT | ND | Cc1ccc(C(=O)NC[C@H]2CN(C(=O)[C@@H]3CCN(C)C3)CCCO2)o1 |
| ZINC000956566890 | -32.50 | 58783667 | 0.29 | 112.5% | NT | NT | NT | ND | CCC(=O)N1CCN([C@H]2CCN(C(=O)c3c[nH]c(=O)cn3)C2)CC1 |
| ZINC000861751799 | -32.50 | 58777328 | 0.23 | 106.3% | NT | NT | NT | ND | Cc1cc(N(C)C)ccc1CNC(=O)C(=O)NC[C@@H]1COCCN1 |
| ZINC000531495018 | -32.50 | 58758088 | 0.28 | 99.3% | NT | NT | NT | ND | CC(=O)N1CCN(C(=O)C(=O)Nc2cnn(-c3ccccn3)c2)CC1 |
| ZINC000422349618 | -32.50 | 58750144 | 0.27 | 129.1% | NT | NT | NT | ND | COC(=O)[C@@]1(C)CCCN(C(=O)C(=O)N[C@@H](C)Cc2ccncc2)C1 |
| ZINC000420063417 | -32.50 | 58749856 | 0.29 | 94.1% | NT | NT | NT | ND | NC[C@@H](NC(=O)C(=O)NCCN1CCSCC1)C1CCCC1 |
| ZINC000361132106 | -32.50 | 58746476 | 0.21 | 76.4% | NT | NT | NT | ND | Cc1nc2n(n1)CCN(C(=O)C(=O)N[C@@H]1CCO[C@@H](CC(C)C)C1)C2 |
| ZINC001328800283 | -32.50 | 58806168 | 0.29 | 86.4% | NT | NT | NT | ND | C[C@@H](CNC(=O)C(=O)NCc1nnn(C)n1)N(C)Cc1ccccc1 |
| ZINC001323717111 | -32.50 | 58805943 | 0.26 | 95.0% | NT | NT | NT | ND | COC(=O)N1CCC[C@H](NC(=O)C(=O)N2CC[C@@](O)(C3CC3)C2)C1 |
| ZINC001261177261 | -32.50 | 58804020 | 0.24 | 70.9% | NT | NT | NT | ND | CCNS(=O)(=O)CCNC(=O)C(=O)N[C@@H](C)C1=CCN(C)CC1 |
| ZINC001202928172 | -32.50 | 58801143 | 0.20 | 117.2% | NT | NT | NT | ND | CN(C)S(=O)(=O)N(C)CC(=O)N[C@@H]1C[C@H](NCCO)C12CCC2 |
| ZINC001199969207 | -32.50 | 58800914 | 0.27 | 124.4% | NT | NT | NT | ND | Cc1cc(C(=O)N2CCCO[C@@H](CNCC(=O)NC3CC3)C2)n(C)n1 |
| ZINC001115282185 | -32.50 | 58795907 | 0.24 | 109.4% | NT | NT | NT | ND | CN(C)S(=O)(=O)CCC(=O)N[C@@H]1[C@H]2CN(Cc3ccon3)C[C@H]21 |
| ZINC001105862991 | -32.50 | 58795331 | 0.21 | 76.1% | NT | NT | NT | ND | Cc1cc(C)nc(NC[C@@H](O)CNC(=O)[C@@H]2CCc3c[nH]nc3C2)n1 |
| ZINC001071984909 | -32.50 | 58792710 | 0.26 | 89.9% | NT | NT | NT | ND | C[C@@H]1CN(C(=O)Cc2cncnc2)C[C@H]1NC(=O)CCn1cccn1 |
| ZINC001067478391 | -32.50 | 58792564 | 0.27 | 104.3% | NT | NT | NT | ND | O=C(c1cccnn1)N1CCC(O)(CNc2cncc(Cl)n2)CC1 |
| ZINC001046685471 | -32.50 | 58791314 | 0.25 | 101.2% | NT | NT | NT | ND | Cc1nc[nH]c1C(=O)N1CC(O)(CNC(=O)CN2CC[C@H](C)C2)C1 |
| ZINC001043772717 | -32.50 | 58790938 | 0.23 | 116.1% | NT | NT | NT | ND | CN1CCCC[C@@H]1C(=O)NCC1(O)CN(C(=O)CCc2cn[nH]c2)C1 |
| ZINC001041828396 | -32.50 | 58790785 | 0.21 | 103.0% | NT | NT | NT | ND | CN1CC[C@@H](C(=O)N2CC[C@@]3(CCN(Cc4nnn(C)n4)C3)C2)C1 |
| ZINC001041431730 | -32.50 | 58790748 | 0.21 | 98.6% | NT | NT | NT | ND | O=C(CCn1ccnc1)N1CC(O)(CNC(=O)[C@H]2CC=CCC2)C1 |
| ZINC001020327691 | -32.50 | 58788316 | 0.28 | 102.3% | NT | NT | NT | ND | Cc1nc(CN[C@H]2C[C@H](NC(=O)Cn3ncn(C)c3=O)C2)cs1 |
| ZINC001016825715 | -32.50 | 58787985 | 0.29 | 106.5% | NT | NT | NT | ND | C[C@@]1(C(=O)N[C@H]2CC23CCN(CC(=O)NC2CC2)CC3)CCNC1=O |
| ZINC001015510639 | -32.50 | 58787894 | 0.28 | 87.4% | NT | NT | NT | ND | O=C(N[C@H]1CCN(CCn2cc(Cl)cn2)C1)[C@H]1CCNC1=O |
| ZINC000995125582 | -27.50 | 124384340 | 0.29 | 101.7% | NT | NT | NT | ND | CN1CC[C@H](C(=O)NC2CN(C(=O)Cc3ccc(F)cn3)C2)CC1=O |
| ZINC000989806749 | -27.50 | 124383312 | 0.25 | 117.7% | NT | NT | NT | ND | CCn1ncnc1CN1CCCN(C(=O)C[C@H]2CC(=O)NC2=O)CC1 |
| ZINC000973057271 | -27.50 | 124380178 | 0.26 | 110.9% | NT | NT | NT | ND | COCCN1C[C@H](C(=O)N[C@H]2C[C@H](NC(=O)c3cn[nH]c3)C2)CC1=O |
| ZINC000970553575 | -27.50 | 124379913 | 0.25 | 92.8% | NT | NT | NT | ND | COc1ccc(C(=O)N[C@@H]2CN(C(=O)C3(C)CCC3)C[C@@H]2O)nn1 |
| ZINC000941361883 | -27.50 | 124375758 | 0.24 | 110.7% | NT | NT | NT | ND | Cc1cc(NC(=O)C(=O)N2CC[C@@H](NC(=O)c3cncn3C)C2)no1 |
| ZINC000850493342 | -27.50 | 124360876 | 0.26 | 135.5% | NT | NT | NT | ND | CON(C)CCNC(=O)C(=O)Nc1ccc2c(c1)C(=O)N(C)C2=O |
| ZINC000748395040 | -27.50 | 124346989 | 0.22 | 108.1% | NT | NT | NT | ND | CN(C(=O)CCn1cnc2c1c(=O)n(C)c(=O)n2C)c1cn[nH]c1 |
| ZINC000659691253 | -27.50 | 124339922 | 0.23 | 82.1% | NT | NT | NT | ND | CNC[C@H](C)NC(=O)C(=O)NCc1ccc(N2CCO[C@@H](C)C2)nc1 |
| ZINC000618647685 | -27.50 | 124332212 | 0.30 | 91.7% | NT | NT | NT | ND | CN(C)S(=O)(=O)N(C)CC(=O)N[C@@H]1COc2cc(F)ccc2C1 |
| ZINC000529344158 | -27.50 | 124318081 | 0.22 | 134.8% | NT | NT | NT | ND | O=C(NC[C@@H]1CNCCO1)N1CCN(S(=O)(=O)C2CC2)CC1 |
| ZINC000442921898 | -27.50 | 124305560 | 0.24 | 111.1% | NT | NT | NT | ND | CO[C@H](C)CS(=O)(=O)N1CCN(C(=O)C(=O)N2CCCC2)CC1 |
| ZINC000408001865 | -27.50 | 124298896 | 0.22 | 110.0% | NT | NT | NT | ND | O=C(c1cncc(N2CCOCC2)c1)N1CCN2C(=O)NC[C@@H]2C1 |
| ZINC001363211634 | -27.50 | 124423462 | 0.23 | 78.6% | NT | NT | NT | ND | O=C(NCC1CN(C(=O)[C@@H]2CCc3nnc(O)n3C2)C1)C(F)(F)F |
| ZINC001350153389 | -27.50 | 124422681 | 0.22 | 93.3% | NT | NT | NT | ND | O=C(Nc1ccc2c(cc[nH]c2=O)c1)C(=O)N1C[C@H](O)[C@@H](CO)C1 |
| ZINC001341791776 | -27.50 | 124421977 | 0.27 | 221.7% | NT | NT | NT | ND | COc1cccc2c1OC[C@H](NC(=O)C(=O)N[C@@H]1CC(=O)N(C)C1)C2 |
| ZINC001119470858 | -27.50 | 124402534 | 0.34 | 106.8% | NT | NT | NT | ND | O=C(NCCN1CCNC1=O)C(=O)NC[C@H]1CCc2ccccc21 |
| ZINC001119173847 | -27.50 | 124402491 | 0.23 | 95.8% | NT | NT | NT | ND | CO[C@@H](CNC(=O)C(=O)N1C[C@@H]2C(=O)N(C)C(=O)[C@@H]2C1)CC(C)C |
| ZINC001109958062 | -27.50 | 124401435 | 0.23 | 90.2% | NT | NT | NT | ND | O=C(CN1CCNC(=O)C1)N[C@H](CNc1cc(F)ncn1)C1CC1 |
| ZINC001074702922 | -27.50 | 124396458 | 0.22 | 109.0% | NT | NT | NT | ND | NC(=O)C(=O)N1CCC(O)(CNC(=O)[C@H]2CCCCC2(F)F)CC1 |
| ZINC001065434920 | -27.50 | 124395306 | 0.25 | 57.8% | NT | NT | NT | ND | Cn1cc(C(=O)N2CCO[C@@H](CNC(=O)C=C3CCC3)C2)ncc1=O |
| ZINC001060201073 | -27.50 | 124394662 | 0.23 | 76.0% | NT | NT | NT | ND | O=C(NCC[C@@H]1CCN(C(=O)c2ccncn2)C1)c1c[nH]c(=O)cn1 |
| ZINC001058732454 | -27.50 | 124394273 | 0.22 | 93.6% | NT | NT | NT | ND | Cn1nnc2c1ncnc2N1C[C@H]2CC[C@@H](C1)N2C(=O)c1cc[nH]n1 |
| ZINC001058532763 | -27.50 | 124394246 | 0.27 | 95.3% | NT | NT | NT | ND | O=C(N[C@H]1CCN(c2cc(Cl)c(O)nn2)C1)c1c[nH]c(=O)cn1 |
| ZINC001051633080 | -27.50 | 124393286 | 0.23 | 131.9% | NT | NT | NT | ND | O=C(Cn1cnnn1)N[C@H]1CCN(c2ncnc3c2CCC3)C[C@H]1O |
| ZINC001046261579 | -27.50 | 124392461 | 0.23 | 114.1% | NT | NT | NT | ND | CCC[C@@H]1C[C@H]1C(=O)NCC1(O)CN(C(=O)C2CN(C(C)=O)C2)C1 |
| ZINC000998978244 | -22.50 | 236805874 | 0.23 | 98.3% | NT | NT | NT | ND | Cc1cnoc1C(=O)NC1CN(C(=O)[C@@H]2CCN(C)S2(=O)=O)C1 |
| ZINC000981250423 | -22.50 | 236800515 | 0.22 | 97.2% | NT | NT | NT | ND | Cn1nncc1C(=O)NC[C@H]1CC[C@@H](NC(=O)c2cnns2)C1 |
| ZINC000968717635 | -22.50 | 236797474 | 0.22 | 94.6% | NT | NT | NT | ND | CCc1c(C(=O)N[C@@H]2CN(C(=O)c3n[nH]cc3C)C[C@@H]2O)cnn1C |
| ZINC000965338365 | -22.50 | 236796664 | 0.19 | 99.1% | NT | NT | NT | ND | O=C([C@H]1CCOC1)N1CCOC2(CN(C(=O)[C@@]3(F)CCOC3)C2)C1 |
| ZINC000961583379 | -22.50 | 236795906 | 0.22 | 91.1% | NT | NT | NT | ND | Cc1c(C(=O)N[C@@H]2CN(C(=O)C3=COCCC3)C[C@@H]2O)cnn1C |
| ZINC000955144887 | -22.50 | 236794263 | 0.19 | 96.9% | NT | NT | NT | ND | CN(C(=O)c1c[nH]c(=O)cn1)C1CN(C(=O)c2n[nH]c3c2CCC3)C1 |
| ZINC000942457739 | -22.50 | 236790192 | 0.29 | 89.4% | NT | NT | NT | ND | Cn1nc(C(=O)N[C@@H]2CCN(C(=O)Cc3cscn3)C2)ccc1=O |
| ZINC000938242679 | -22.50 | 236788459 | 0.27 | 107.1% | NT | NT | NT | ND | Cc1nc(C)c(CC(=O)N2CC[C@H](NC(=O)c3cnn[nH]3)C2)c(O)n1 |
| ZINC000907721339 | -22.50 | 236782078 | 0.19 | 118.7% | NT | NT | NT | ND | CS(=O)(=O)N1CC(C(=O)N2CC(=O)N(CC(F)(F)F)C2)C1 |
| ZINC000625246444 | -22.50 | 236732519 | 0.18 | 105.2% | NT | NT | NT | ND | COCC(COC)N1CCN(S(=O)(=O)c2cnc(C)n2C)CC1 |
| ZINC000407973885 | -22.50 | 236691723 | 0.24 | 120.4% | NT | NT | NT | ND | O=C(NC1CC1)C(=O)N1CCN(S(=O)(=O)C[C@@H]2CCCO2)CC1 |
| ZINC000345525893 | -22.50 | 236681776 | 0.19 | 90.8% | NT | NT | NT | ND | C[C@@H]1OCC[C@]1(C)NC(=O)CCc1nc2c([nH]1)n(C)c(=O)nc2O |
| ZINC001413380655 | -22.50 | 236868558 | 0.23 | 117.9% | NT | NT | NT | ND | CC(=O)N[C@@H]1CCN(S(=O)(=O)c2cnn([C@@H]3CCOC3)c2)C1 |
| ZINC001363999416 | -22.50 | 236861114 | 0.19 | 104.5% | NT | NT | NT | ND | O=S(=O)(N[C@@H]1CCc2nnnn2CC1)c1cnc2n1CCCC2 |
| ZINC001326422624 | -22.50 | 236856302 | 0.22 | 99.2% | NT | NT | NT | ND | COC(=O)[C@@H]1C[C@H](O)CN(C(=O)C(=O)N[C@H](C)C2CCOCC2)C1 |
| ZINC001261948364 | -22.50 | 236850697 | 0.25 | 90.8% | NT | NT | NT | ND | CN1C(=O)[C@H]2CN(C(=O)C(=O)NC3CCC(F)(F)CC3)C[C@H]2C1=O |
| ZINC001193095852 | -22.50 | 236843572 | 0.25 | 112.2% | NT | NT | NT | ND | CCOCCNC(=O)CN1C[C@@H](O)[C@H](NC(=O)[C@]23C[C@H]2COC3)C1 |
| ZINC001125309519 | -22.50 | 236835764 | 0.22 | 88.0% | NT | NT | NT | ND | CN(C[C@@H](O)CN(C)c1ncc(F)cn1)C(=O)CCc1cnn[nH]1 |
| ZINC001063398490 | -22.50 | 236822006 | 0.26 | 145.9% | NT | NT | NT | ND | CN(C)C(=O)c1cncc(N[C@H]2CC[C@H](NC(=O)C(N)=O)CC2)n1 |
| ZINC001047126822 | -22.50 | 236818242 | 0.22 | 107.2% | NT | NT | NT | ND | CC(C)n1cc(C(=O)N2CC(O)(CNC(=O)c3ccnnc3)C2)nn1 |
| ZINC001046939982 | -22.50 | 236818218 | 0.22 | 110.5% | NT | NT | NT | ND | NC(=O)[C@@H]1CCC[C@@H](C(=O)NCC2(O)CN(C(=O)C3CCC3)C2)C1 |
| ZINC001046914841 | -22.50 | 236818208 | 0.21 | 103.7% | NT | NT | NT | ND | C[C@H](C(=O)NCC1(O)CN(C(=O)Cn2ncnn2)C1)C1CCCC1 |
| ZINC001043817104 | -22.50 | 236817302 | 0.20 | 88.5% | NT | NT | NT | ND | CN(C)c1ccc(C(=O)N2CC(O)(CNC(=O)c3cnn[nH]3)C2)nc1 |
| ZINC001043696212 | -22.50 | 236817254 | 0.25 | 84.5% | NT | NT | NT | ND | CNC(=O)NCC(=O)N1CC(O)(CNC(=O)c2ccc(F)s2)C1 |
| ZINC001042722400 | -22.50 | 236816851 | 0.20 | 125.2% | NT | NT | NT | ND | Cn1nc(C(=O)N2CC(O)(CNC(=O)C3=CCOCC3)C2)ccc1=O |
| ZINC001040614201 | -22.50 | 236816227 | 0.22 | 117.3% | NT | NT | NT | ND | COCCC(=O)N1CC[C@@](O)(CNC(=O)[C@@H]2CCN(C)C2=O)C1 |
| ZINC001025444165 | -22.50 | 236812931 | 0.24 | 95.7% | NT | NT | NT | ND | Cn1nccc1C(=O)N1C[C@@H](O)[C@H](NC(=O)c2c[nH]nc2C2CC2)C1 |
| ZINC001003297636 | -22.50 | 236807089 | 0.22 | 72.3% | NT | NT | NT | ND | O=C(Cn1cnnn1)N1CC(CNC(=O)C23CCCN2CCC3)C1 |

^†^TC, Tanimoto coefficient to sigma ligands from ChEMBL.

**Supplementary Information Table 4** | **Measured pharmacokinetic parameters for PB28, Z1665845742, Z4446724338 and Z4857158944 in male CD-1 mice by 10 mg/kg subcutaneous administration.**

| Pharmacokinetic Parameters | | | | | | | |
| --- | --- | --- | --- | --- | --- | --- | --- |
| Type | Name | T_max_, min | C_max_, ng/ml (g) | AUC_0→t_ (AUC_last_) ng*min/ml (g) | AUC_0→∞_(AUC_INF_obs_),ng*min/ml (g) | T_1/2_  (HL_Lambda_z),min | K_el_ (Lambda_z), min^-1^ |
| Plasma | Z1665845742 | 20 | 968 | 99000 | 112000 | 185 | 0.00374 |
|  | Z4446724338 | 20 | 449 | 58300 | 60500 | 47.4 | 0.0146 |
|  | Z4857158944 | 20 | 288 | 13300 | 14200 | 27.8 | 0.0249 |
|  | PB28 | 60 | 42 | 8640 | 45900 | 740 | 0.000937 |
| Brain | Z1665845742 | 20 | 3150 | 436000 | 509000 | 747 | 0.000928 |
|  | Z4446724338 | 20 | 7390 | 1140000 | 1150000 | 69.7 | 0.00995 |
|  | Z4857158944 | 20 | 2960 | 247000 | 327000 | 452 | 0.00153 |
|  | PB28 | 60 | 948 | 229000 | 240000 | 98.1 | 0.00706 |

### **Supplemental Data Table 5 | Radioligand binding affinities of a panel of potential off-targets.** Full binding curves were done on targets that at 10 μM compound concentration showed more than 50% radioligand displacement

| **Z1665845742** | | | |
| --- | --- | --- | --- |
| **Target name** | **gene** | **pK_i_ (n≥3)** | **SEM** |
| 5-hydroxytryptamine receptor 1B | HTR1B | 5.31 | 0.05 |
| 5-hydroxytryptamine receptor 1D | HTR1D | 5.72 | 0.08 |
| 5-Hydroxytryptamine receptor 2B | HTR2B | 6.03 | 0.06 |
| 5-hydroxytryptamine receptor 3A | HTR3A | 5.88 | 0.02 |
| 5-hydroxytryptamine receptor 7 | HTR7 | 5.60 | 0.08 |
| Dopamine receptor D1 | DRD1 | 5.45 | 0.13 |
| Dopamine receptor D2 | DRD2 | 6.32 | 0.03 |
| Dopamine receptor D3 | DRD3 | 6.91 | 0.05 |
| Dopamine receptor D4 | DRD4 | 6.61 | 0.04 |
| Histamine H1 receptor | HRH1 | 7.72 | 0.06 |
| Muscarinic acetylcholine receptor M5 | CHRM5 | 5.61 | 0.05 |
| α1A adrenergic receptor | ADRA1A | 6.05 | 0.05 |
| α2A adrenergic receptor | ADRA2A | 5.28 | 0.04 |
| α2B adrenergic receptor | ADRA2B | 5.26 | 0.01 |
| α2C adrenergic receptor | ADRA2C | 6.24 | 0.05 |
| Dopamine transporter | SLC6A3 | 6.00 | 0.03 |
| Noradrenaline transporter | SLC6A2 | 6.40 | 0.09 |
| Serotonin transporter | SLC6A4 | 7.26 | 0.04 |
| β2 adrenergic receptor | ADRB2 | 5.42 | 0.05 |
| **Z4446724338** | | | |
| **Target name** | **gene** | **pK_i_ (n=3)** | **SEM** |
| 5-Hydroxytryptamine receptor 2B | HTR2B | 6.42 | 0.09 |
| Dopamine receptor D3 | DRD3 | 5.65 | 0.01 |
| Dopamine receptor D4 | DRD4 | 5.28 | 0.03 |
| **Z4857158944** | | | |
| **Target name** | **gene** | **pK_i_ (n=3)** | **SEM** |
| Dopamine receptor D4 | DRD4 | 5.41 | 0.03 |
| Dopamine transporter | SLC6A3 | 5.66 | 0.04 |
| Noradrenaline transporter | SLC6A2 | 5.92 | 0.15 |
| Serotonin transporter | SLC6A4 | 5.71 | 0.07 |

### **Supplemental Data Table 6 | Purity information of 508 σ_2_ primary hits.**

| ZINC ID | Vendor ID | | Purity (%) |
| --- | --- | --- | --- |
| ZINC001460371878 | | Z4670289310 | 90 |
| ZINC001515357045 | | Z3142060526 | 98 |
| ZINC000152090354 | | Z1403332363 | 91 |
| ZINC000153400973 | | Z4855605004 | 91 |
| ZINC000153451054 | | Z1400756133 | 100 |
| ZINC000155112667 | | Z1440398442 | 100 |
| ZINC000340106708 | | Z3007447312 | 99 |
| ZINC000343016571 | | Z2049584546 | 90 |
| ZINC000345525893 | | Z1657504296 | 91 |
| ZINC000345970626 | | Z2156547523 | 99 |
| ZINC000347007006 | | Z1817676002 | 91 |
| ZINC000349353698 | | Z1869123365 | 94 |
| ZINC000358051108 | | Z1687252313 | 100 |
| ZINC000358062278 | | Z1687262970 | 100 |
| ZINC000358071102 | | Z2128309149 | 100 |
| ZINC000358253868 | | Z1687405419 | 94 |
| ZINC000359009035 | | Z1639222228 | 98 |
| ZINC000361132106 | | Z2143313061 | 100 |
| ZINC000362611503 | | Z1302947967 | 98 |
| ZINC000366544782 | | Z2208128794 | 100 |
| ZINC000367601434 | | Z2070861096 | 100 |
| ZINC000370893072 | | Z1185966042 | 100 |
| ZINC000415549177 | | Z2355792372 | 96 |
| ZINC000416493836 | | Z2358916962 | 100 |
| ZINC000421699823 | | Z2372596717 | 100 |
| ZINC000421701270 | | Z2372594890 | 100 |
| ZINC000495656270 | | Z2015801612 | 100 |
| ZINC000480785335 | | Z1958788373 | 97 |
| ZINC000438905691 | | Z1486352592 | 100 |
| ZINC000426471156 | | Z1745777108 | 100 |
| ZINC000450551910 | | Z2272953139 | 90 |
| ZINC000442921898 | | Z1998020952 | 99 |
| ZINC000520440291 | | Z125769190 | 100 |
| ZINC000423867812 | | Z2439684203 | 100 |
| ZINC000454705882 | | Z2291985817 | 100 |
| ZINC000438095302 | | Z1783454190 | 99 |
| ZINC000464767692 | | Z1289778247 | 100 |
| ZINC000500192494 | | Z1193391202 | 100 |
| ZINC000465804078 | | Z1283002963 | 99 |
| ZINC000452100574 | | Z2280166407 | 91 |
| ZINC000512182007 | | Z2070843529 | 100 |
| ZINC000501775665 | | Z1557138164 | 100 |
| ZINC000451319844 | | Z2277002329 | 98 |
| ZINC000421710394 | | Z2372599855 | 100 |
| ZINC000468894194 | | Z1414793302 | 99 |
| ZINC000439311985 | | Z1990441865 | 90 |
| ZINC000444807044 | | Z1771761660 | 91 |
| ZINC000453607577 | | Z2287450150 | 100 |
| ZINC000508455082 | | Z2068703050 | 100 |
| ZINC000430847713 | | Z1868664675 | 93 |
| ZINC000426526480 | | Z1745772472 | 100 |
| ZINC000421710406 | | Z2372600038 | 100 |
| ZINC000513818302 | | Z1011053398 | 98 |
| ZINC000516964180 | | Z4262616955 | 96 |
| ZINC000504552433 | | Z2003650908 | 100 |
| ZINC000466933503 | | Z1408680787 | 91 |
| ZINC000496403719 | | Z2407112450 | 97 |
| ZINC000481989921 | | Z1732943230 | 100 |
| ZINC000446751851 | | Z1782283561 | 100 |
| ZINC000429522320 | | Z1865690982 | 100 |
| ZINC000508475661 | | Z2068702592 | 99 |
| ZINC000422349618 | | Z2375737391 | 90 |
| ZINC000440321606 | | Z1375037087 | 90 |
| ZINC000518003593 | | Z1141963200 | 91 |
| ZINC000045463977 | | Z495734882 | 100 |
| ZINC000453351921 | | Z2286568343 | 98 |
| ZINC000453336584 | | Z2286516142 | 93 |
| ZINC000436303801 | | Z1401051328 | 100 |
| ZINC001004070100 | | Z4670319509 | 100 |
| ZINC001125309519 | | Z4670328554 | 98 |
| ZINC001073859800 | | Z4670321669 | 97 |
| ZINC001033739722 | | Z4422878790 | 100 |
| ZINC001048683995 | | Z4670289756 | 100 |
| ZINC001051553588 | | Z4670323310 | 100 |
| ZINC000972168981 | | Z4670319314 | 94 |
| ZINC001399642198 | | Z4670319495 | 98 |
| ZINC001040612984 | | Z4670289716 | 99 |
| ZINC001052175095 | | Z4670323317 | 100 |
| ZINC001033973784 | | Z4670319237 | 100 |
| ZINC001058732454 | | Z4670325694 | 100 |
| ZINC001054647418 | | Z4670321663 | 100 |
| ZINC001051633080 | | Z4670325829 | 91 |
| ZINC001391502525 | | Z4670289721 | 98 |
| ZINC001002391793 | | Z4670321604 | 95 |
| ZINC001067478391 | | Z4670289904 | 97 |
| ZINC001376084945 | | Z4670289377 | 92 |
| ZINC000971070823 | | Z4670319896 | 100 |
| ZINC001023604137 | | Z4670323256 | 97 |
| ZINC001377717241 | | Z4670320150 | 100 |
| ZINC001028614115 | | Z4670321630 | 98 |
| ZINC001115282185 | | Z4670289926 | 91 |
| ZINC001041828396 | | Z4670323530 | 91 |
| ZINC001073886256 | | Z4670323443 | 91 |
| ZINC001028403773 | | Z4670321624 | 98 |
| ZINC001206651458 | | Z4835694083 | 98 |
| ZINC001058532763 | | Z4670325690 | 96 |
| ZINC001202928172 | | Z4670325486 | 91 |
| ZINC001041605786 | | Z4670321432 | 100 |
| ZINC001015510639 | | Z4670323500 | 100 |
| ZINC000989806749 | | Z4670325791 | 91 |
| ZINC001381212829 | | Z4670289720 | 91 |
| ZINC001039123057 | | Z4670321643 | 100 |
| ZINC001032169623 | | Z4670289715 | 100 |
| ZINC000671981913 | | Z2084190151 | 100 |
| ZINC001046939982 | | Z4670328521 | 100 |
| ZINC000982403760 | | Z4670289880 | 91 |
| ZINC000937267298 | | Z4670321700 | 98 |
| ZINC000936344871 | | Z4670323383 | 99 |
| ZINC000942457739 | | Z4670289985 | 92 |
| ZINC000981250423 | | Z4670289989 | 97 |
| ZINC000774469834 | | Z2028809437 | 92 |
| ZINC000968717635 | | Z4670325953 | 94 |
| ZINC000418924570 | | Z2365003665 | 100 |
| ZINC001046914841 | | Z4670326023 | 96 |
| ZINC000961583379 | | Z4670325927 | 100 |
| ZINC000970553575 | | Z4670325632 | 100 |
| ZINC000997292482 | | Z4670323012 | 100 |
| ZINC000995125582 | | Z4670325676 | 99 |
| ZINC000950078544 | | Z4670321496 | 98 |
| ZINC000283462313 | | Z2027890301 | 98 |
| ZINC000956566890 | | Z4670325167 | 100 |
| ZINC001047126822 | | Z4670290017 | 100 |
| ZINC000973375109 | | Z4670321253 | 100 |
| ZINC001323717111 | | Z2083094175 | 96 |
| ZINC001326422624 | | Z2441305823 | 97 |
| ZINC000965338365 | | Z4670328442 | 100 |
| ZINC000955144887 | | Z4670290004 | 94 |
| ZINC000991393621 | | Z4670289823 | 100 |
| ZINC000938242679 | | Z4670325888 | 100 |
| ZINC000861751799 | | Z2713237703 | 99 |
| ZINC000995014137 | | Z4670289824 | 90 |
| ZINC000973057271 | | Z4670325660 | 96 |
| ZINC000998978244 | | Z4670328470 | 100 |
| ZINC000955102158 | | Z4670321756 | 99 |
| ZINC000941361883 | | Z4670289963 | 95 |
| ZINC000408001865 | | Z1522160855 | 100 |
| ZINC000073649189 | | Z268742176 | 100 |
| ZINC000684607185 | | Z1959404885 | 100 |
| ZINC000412048402 | | Z2330881599 | 100 |
| ZINC000743860012 | | Z1289706494 | 100 |
| ZINC000932649435 | | Z2954002723 | 93 |
| ZINC000074484650 | | Z1204057701 | 98 |
| ZINC000407973885 | | Z1003275534 | 100 |
| ZINC000748395040 | | Z1334198857 | 100 |
| ZINC000929842198 | | Z2947439720 | 97 |
| ZINC000825615260 | | Z2268071152 | 100 |
| ZINC000375486080 | | Z1817708188 | 99 |
| ZINC000819225653 | | Z2406633347 | 100 |
| ZINC000672564445 | | Z2102941193 | 100 |
| ZINC000667676804 | | Z2356289856 | 98 |
| ZINC000891830660 | | Z2968125940 | 100 |
| ZINC000683045155 | | Z1846214920 | 100 |
| ZINC000932000298 | | Z2952842982 | 100 |
| ZINC000894427229 | | Z2973188608 | 99 |
| ZINC000894429059 | | Z2973188617 | 100 |
| ZINC001350153389 | | Z2591424667 | 100 |
| ZINC000830877226 | | Z2277004157 | 93 |
| ZINC000421295130 | | Z2371762623 | 98 |
| ZINC000802379169 | | Z2021149846 | 93 |
| ZINC000664479370 | | Z2615347132 | 100 |
| ZINC000076836015 | | Z645938956 | 95 |
| ZINC000813849560 | | Z2202566421 | 100 |
| ZINC000679555790 | | Z1629918162 | 100 |
| ZINC000907721339 | | Z2907375086 | 91 |
| ZINC000897504395 | | Z2977471156 | 100 |
| ZINC000684407814 | | Z1950677214 | 100 |
| ZINC000378210510 | | Z2437464324 | 98 |
| ZINC000679371269 | | Z3080218908 | 100 |
| ZINC000726467110 | | Z1021762532 | 98 |
| ZINC000376738332 | | Z2002546099 | 100 |
| ZINC000886762228 | | Z2863431531 | 100 |
| ZINC000775588849 | | Z1731557920 | 100 |
| ZINC000377772869 | | Z1249310378 | 91 |
| ZINC000683721842 | | Z1898133468 | 100 |
| ZINC000419445843 | | Z2366046350 | 100 |
| ZINC000413145325 | | Z2347073394 | 100 |
| ZINC000804054066 | | Z2084065212 | 100 |
| ZINC000875242850 | | Z2809328002 | 100 |
| ZINC000409447221 | | Z1653378877 | 100 |
| ZINC000878084395 | | Z2833839133 | 100 |
| ZINC000678272880 | | Z1569257017 | 97 |
| ZINC000766031584 | | Z1731397861 | 100 |
| ZINC000685844120 | | Z2176977027 | 100 |
| ZINC000410783840 | | Z2320517885 | 91 |
| ZINC000661577595 | | Z2603376298 | 91 |
| ZINC000805503179 | | Z2100673253 | 100 |
| ZINC000809405032 | | Z2158241671 | 91 |
| ZINC000080247921 | | Z1357258969 | 92 |
| ZINC000676461131 | | Z1473280384 | 99 |
| ZINC000668631550 | | Z2629138429 | 98 |
| ZINC000370933962 | | Z1500052245 | 100 |
| ZINC000880249690 | | Z2842451641 | 91 |
| ZINC000419873354 | | Z2367259197 | 98 |
| ZINC000762555244 | | Z1686821209 | 95 |
| ZINC000872109019 | | Z2790627889 | 94 |
| ZINC000363920872 | | Z1414243280 | 100 |
| ZINC000669616349 | | Z1997384440 | 91 |
| ZINC000420063417 | | Z2367984694 | 91 |
| ZINC000673584800 | | Z2145850283 | 91 |
| ZINC000878406056 | | Z2835144760 | 90 |
| ZINC000417806012 | | ZINC000417806012 | 100 |
| ZINC001362573982 | | Z4670289309 | 96 |
| ZINC000270395954 | | Z1526216282 | 100 |
| ZINC000194878845 | | Z1944110869 | 100 |
| ZINC000271284821 | | Z1559388480 | 100 |
| ZINC000285725404 | | Z2070851195 | 100 |
| ZINC000286616311 | | Z2097603634 | 99 |
| ZINC001363999416 | | Z4670328397 | 100 |
| ZINC000245453841 | | Z1367354685 | 95 |
| ZINC001413380655 | | Z4670328398 | 100 |
| ZINC000532978237 | | Z1212880913 | 100 |
| ZINC000533411740 | | Z1051819646 | 100 |
| ZINC000332414923 | | Z1151758209 | 100 |
| ZINC000531059007 | | Z1315510635 | 100 |
| ZINC000191344346 | | Z1828654465 | 100 |
| ZINC000228149300 | | Z1499116079 | 100 |
| ZINC000248917461 | | Z1796863649 | 100 |
| ZINC000301801549 | | Z1510214726 | 92 |
| ZINC000272631760 | | Z1630257371 | 97 |
| ZINC000182663590 | | Z1659072002 | 100 |
| ZINC000265181994 | | Z1212877902 | 98 |
| ZINC001579778746 | | Z4670319222 | 100 |
| ZINC000301468116 | | Z1272628292 | 100 |
| ZINC000286502332 | | Z2071413788 | 100 |
| ZINC000186482223 | | Z1745468890 | 100 |
| ZINC000263644082 | | Z1021095168 | 100 |
| ZINC000276781830 | | Z1799344772 | 91 |
| ZINC000286782676 | | Z2439248019 | 100 |
| ZINC001363211634 | | Z4670325728 | 95 |
| ZINC000237909071 | | Z4079013418 | 100 |
| ZINC000246894550 | | Z1286462398 | 97 |
| ZINC000182372239 | | Z1612136617 | 100 |
| ZINC000334024070 | | Z2330904130 | 100 |
| ZINC000135482356 | | Z1528618055 | 91 |
| ZINC000192810020 | | Z2002269140 | 100 |
| ZINC000339640280 | | Z2029898084 | 91 |
| ZINC000297448741 | | Z2315500637 | 100 |
| ZINC000524392254 | | Z1203509645 | 100 |
| ZINC000526962900 | | Z1802193579 | 98 |
| ZINC000555574173 | | Z2027955462 | 100 |
| ZINC000188412270 | | Z2435429359 | 100 |
| ZINC000531495018 | | Z1386110604 | 100 |
| ZINC000542825037 | | Z1636174218 | 99 |
| ZINC000536865847 | | Z1188052306 | 100 |
| ZINC000155998528 | | Z1452748512 | 97 |
| ZINC000534109220 | | Z1535021318 | 93 |
| ZINC000192043753 | | Z1844778656 | 97 |
| ZINC000525245955 | | Z1251593378 | 91 |
| ZINC000279187431 | | Z1907290928 | 92 |
| ZINC000178638284 | | Z1440003064 | 100 |
| ZINC000265281429 | | Z1215601135 | 99 |
| ZINC000285662957 | | Z2070980036 | 100 |
| ZINC000136880129 | | Z2434891003 | 100 |
| ZINC001474404955 | | Z4670289313 | 100 |
| ZINC000170742095 | | Z1051811136 | 100 |
| ZINC000539406733 | | Z1592753681 | 100 |
| ZINC001580410121 | | Z3099816607 | 99 |
| ZINC000529649527 | | Z2717370444 | 100 |
| ZINC000518759550 | | Z221129296 | 100 |
| ZINC000298791523 | | Z649291414 | 100 |
| ZINC000287105318 | | Z2101754018 | 100 |
| ZINC000287581634 | | Z2102337335 | 98 |
| ZINC000176805277 | | Z1362216796 | 98 |
| ZINC000550423124 | | Z2219778182 | 95 |
| ZINC000158577469 | | Z1428495312 | 100 |
| ZINC000261552362 | | Z2518207693 | 100 |
| ZINC000529344158 | | Z2712489017 | 100 |
| ZINC000289897491 | | Z2171570653 | 97 |
| ZINC000272592294 | | Z2931805209 | 100 |
| ZINC000159130997 | | Z1414567598 | 100 |
| ZINC000295103757 | | Z2264936388 | 96 |
| ZINC000172648663 | | ZINC000172648663 | 95 |
| ZINC000517634594 | | Z1679871465 | 100 |
| ZINC000247001079 | | Z1492740593 | 96 |
| ZINC000247147334 | | Z1437980050 | 100 |
| ZINC001109713565 | | Z4670289335 | 100 |
| ZINC001193095852 | | Z4670328570 | 100 |
| ZINC001105862991 | | Z4670325448 | 100 |
| ZINC001095312060 | | Z4453148447 | 96 |
| ZINC001418905447 | | Z2227823699 | 100 |
| ZINC001437918206 | | Z4670320167 | 100 |
| ZINC001196417557 | | Z4670289719 | 91 |
| ZINC001002178741 | | Z4670289763 | 90 |
| ZINC001419404744 | | Z4670289759 | 100 |
| ZINC001149399826 | | Z4670320141 | 99 |
| ZINC001016825715 | | Z4670323508 | 97 |
| ZINC000348635286 | | Z4670325357 | 99 |
| ZINC001019309159 | | Z4670320189 | 99 |
| ZINC000981787500 | | Z4670289712 | 100 |
| ZINC001365986475 | | Z4670289834 | 98 |
| ZINC001018665695 | | Z4670321781 | 90 |
| ZINC000738383044 | | Z4510491787 | 97 |
| ZINC001403624658 | | Z4670319505 | 100 |
| ZINC001018723708 | | Z4670289714 | 100 |
| ZINC000662335416 | | Z2605531820 | 100 |
| ZINC001099969021 | | Z4670319526 | 95 |
| ZINC001369886058 | | Z4670289876 | 94 |
| ZINC001109958062 | | Z4670325724 | 100 |
| ZINC000961091031 | | Z2432470658 | 100 |
| ZINC001199969207 | | Z4670325476 | 97 |
| ZINC000903470913 | | Z2899013918 | 92 |
| ZINC001096194954 | | Z4670289375 | 95 |
| ZINC000064804934 | | Z804474412 | 96 |
| ZINC001072835483 | | Z4670321552 | 100 |
| ZINC001061620290 | | Z4670319443 | 100 |
| ZINC001088099598 | | Z4670323323 | 100 |
| ZINC001001445660 | | Z4670321773 | 100 |
| ZINC001074556982 | | Z4670323452 | 100 |
| ZINC001074702922 | | Z4670325844 | 100 |
| ZINC001040614201 | | Z4670328478 | 100 |
| ZINC001020327691 | | Z4670289920 | 98 |
| ZINC001055138151 | | Z4670289847 | 98 |
| ZINC000960887654 | | Z4670289710 | 96 |
| ZINC001046261579 | | Z4670325687 | 100 |
| ZINC001071984909 | | Z4670323682 | 96 |
| ZINC001041431730 | | Z4670289899 | 100 |
| ZINC001061317651 | | Z4670289717 | 97 |
| ZINC001025444165 | | Z4670326010 | 94 |
| ZINC001000730978 | | Z4670321513 | 100 |
| ZINC001416718427 | | Z4670323372 | 96 |
| ZINC001077034834 | | Z4670319519 | 98 |
| ZINC001003018745 | | Z4670289792 | 97 |
| ZINC001065434920 | | Z4670325839 | 100 |
| ZINC001042098162 | | Z4670289865 | 100 |
| ZINC001019525589 | | Z4670323028 | 100 |
| ZINC001060201073 | | Z4670325704 | 93 |
| ZINC001046685471 | | Z4670325416 | 91 |
| ZINC001055479120 | | Z4670321532 | 100 |
| ZINC001021041334 | | Z4670321424 | 100 |
| ZINC001043817104 | | Z4670326017 | 97 |
| ZINC001003297636 | | Z4670289993 | 91 |
| ZINC001043696212 | | Z4670290013 | 93 |
| ZINC001021109191 | | Z4670289863 | 100 |
| ZINC000941634458 | | Z4670319230 | 100 |
| ZINC001006004942 | | Z4670289793 | 96 |
| ZINC001063398490 | | Z4670290018 | 100 |
| ZINC001042722400 | | Z4670290011 | 92 |
| ZINC001043772717 | | Z4670289921 | 91 |
| ZINC000850493342 | | Z2591590149 | 96 |
| ZINC000605902355 | | Z1236175503 | 100 |
| ZINC000617924178 | | Z2644404614 | 96 |
| ZINC000625246444 | | Z2703748509 | 100 |
| ZINC000608224751 | | Z1275896696 | 95 |
| ZINC000645238909 | | Z2219723727 | 100 |
| ZINC000584233558 | | Z2453568167 | 95 |
| ZINC000805335094 | | Z2099860887 | 94 |
| ZINC000863492725 | | Z2722031649 | 100 |
| ZINC000843092160 | | Z2444981000 | 98 |
| ZINC000556573026 | | Z2047345337 | 100 |
| ZINC000586751050 | | Z1005477330 | 100 |
| ZINC000657977970 | | Z2591377436 | 100 |
| ZINC000595790972 | | Z2604189896 | 100 |
| ZINC000621202771 | | Z4106421794 | 100 |
| ZINC000847525041 | | Z2497215184 | 100 |
| ZINC001307701283 | | Z1732470338 | 100 |
| ZINC000839577413 | | Z2329851488 | 100 |
| ZINC000577603040 | | Z2406352462 | 96 |
| ZINC000559413424 | | Z2219774495 | 100 |
| ZINC000849049798 | | Z2518536383 | 100 |
| ZINC000574285278 | | Z2105779952 | 100 |
| ZINC000130668661 | | Z1564946356 | 91 |
| ZINC000659691253 | | Z2596554540 | 100 |
| ZINC000576277949 | | Z2439481656 | 97 |
| ZINC001261177261 | | Z2098910446 | 97 |
| ZINC000131303503 | | Z1500985969 | 100 |
| ZINC001119173847 | | Z2811678854 | 100 |
| ZINC001261948364 | | Z2808518804 | 100 |
| ZINC001320149247 | | Z1675645742 | 100 |
| ZINC000631236280 | | Z2770318131 | 100 |
| ZINC000556797005 | | Z2317426730 | 100 |
| ZINC001169307444 | | Z4670289382 | 92 |
| ZINC000589606863 | | Z1070654284 | 100 |
| ZINC000584028046 | | Z1823017442 | 91 |
| ZINC001119470858 | | Z2880613867 | 100 |
| ZINC000575186982 | | Z1545287634 | 99 |
| ZINC000858362097 | | Z2698839159 | 100 |
| ZINC000806327364 | | Z2107898889 | 100 |
| ZINC000624110289 | | Z2699497969 | 100 |
| ZINC000124982852 | | Z1562892537 | 98 |
| ZINC000646273308 | | Z2439036917 | 100 |
| ZINC001156463732 | | Z3333965222 | 100 |
| ZINC000656499190 | | Z2312321347 | 98 |
| ZINC000556484619 | | Z2222605980 | 91 |
| ZINC000557493019 | | Z2089300282 | 100 |
| ZINC000611661177 | | Z1640841632 | 100 |
| ZINC000592403811 | | Z3209795395 | 100 |
| ZINC000655168559 | | Z2500262355 | 97 |
| ZINC000588863575 | | Z2143998367 | 100 |
| ZINC000623468108 | | Z2697650278 | 91 |
| ZINC001328800283 | | Z2104441192 | 100 |
| ZINC000133744458 | | Z2437898285 | 97 |
| ZINC000609803478 | | Z1225724640 | 100 |
| ZINC000115566238 | | Z1212900047 | 91 |
| ZINC000603324806 | | Z1191804904 | 100 |
| ZINC000866213504 | | Z2771892019 | 91 |
| ZINC000595353335 | | Z2599443822 | 100 |
| ZINC000120571916 | | Z1245068947 | 100 |
| ZINC000618647685 | | Z2645787888 | 100 |
| ZINC001341791776 | | Z2083030491 | 99 |
| ZINC000627254962 | | Z2712887551 | 91 |
| ZINC000574318236 | | Z2157724660 | 100 |
| ZINC000564110976 | | Z2145852422 | 99 |
| ZINC000269280100 | | ZINC000269280100 | 100 |
| ZINC000435004139 | | ZINC000435004139 | 100 |
| ZINC000595632104 | | ZINC000595632104 | 100 |
| ZINC001237901728 | | ZINC001237901728 | 100 |
| ZINC000662345330 | | ZINC000662345330 | 100 |
| ZINC000247015101 | | ZINC000247015101 | 100 |
| ZINC000262856881 | | ZINC000262856881 | 100 |
| ZINC000262658947 | | ZINC000262658947 | 100 |
| ZINC000170908795 | | ZINC000170908795 | 90.37 |
| ZINC000472356611 | | ZINC000472356611 | 96.02 |
| ZINC000886836194 | | ZINC000886836194 | 91 |
| ZINC000846106280 | | ZINC000846106280 | 97.33 |
| ZINC000792998860 | | ZINC000792998860 | 98.93 |
| ZINC000287374567 | | ZINC000287374567 | 90.14 |
| ZINC001460312963 | | ZINC001460312963 | 97 |
| ZINC000249076038 | | ZINC000249076038 | 100 |
| ZINC000369205980 | | ZINC000369205980 | 97.82 |
| ZINC000533478938 | | ZINC000533478938 | 95.2 |
| ZINC000681377109 | | ZINC000681377109 | 100 |
| ZINC000820594289 | | ZINC000820594289 | 100 |
| ZINC000374368469 | | ZINC000374368469 | 100 |
| ZINC000407281203 | | ZINC000407281203 | 91 |
| ZINC000777733869 | | ZINC000777733869 | 91 |
| ZINC001168222793 | | ZINC001168222793 | 99 |
| ZINC000433044150 | | ZINC000433044150 | 100 |
| ZINC000473272986 | | ZINC000473272986 | 100 |
| ZINC000659267272 | | ZINC000659267272 | 100 |
| ZINC000352856249 | | ZINC000352856249 | 96.64 |
| ZINC000893277758 | | ZINC000893277758 | 100 |
| ZINC000656714762 | | ZINC000656714762 | 98.18 |
| ZINC000801571276 | | ZINC000801571276 | 100 |
| ZINC000665143541 | | ZINC000665143541 | 100 |
| ZINC000133991118 | | ZINC000133991118 | 97.7 |
| ZINC001525937517 | | ZINC001525937517 | 93 |
| ZINC000894101819 | | ZINC000894101819 | 100 |
| ZINC000248559983 | | ZINC000248559983 | 100 |
| ZINC000934332177 | | ZINC000934332177 | 91.96 |
| ZINC000450573233 | | ZINC000450573233 | 98 |
| ZINC000348332392 | | ZINC000348332392 | 100 |
| ZINC001254761628 | | ZINC001254761628 | 95 |
| ZINC000933438523 | | ZINC000933438523 | 100 |
| ZINC000571080072 | | ZINC000571080072 | 100 |
| ZINC000398583627 | | ZINC000398583627 | 100 |
| ZINC000921927365 | | ZINC000921927365 | 100 |
| ZINC000544117725 | | ZINC000544117725 | 93.89 |
| ZINC000336580930 | | ZINC000336580930 | 99 |
| ZINC000892713700 | | ZINC000892713700 | 100 |
| ZINC000656508398 | | ZINC000656508398 | 95.35 |
| ZINC000567338231 | | ZINC000567338231 | 93.64 |
| ZINC000452107481 | | ZINC000452107481 | 100 |
| ZINC000911907143 | | ZINC000911907143 | 100 |
| ZINC000621267824 | | ZINC000621267824 | 100 |
| ZINC000448446275 | | ZINC000448446275 | 100 |
| ZINC001170548029 | | ZINC001170548029 | 95 |
| ZINC000483826940 | | ZINC000483826940 | 98.29 |
| ZINC000296435291 | | ZINC000296435291 | 100 |
| ZINC000176995469 | | ZINC000176995469 | 90 |
| ZINC000452023252 | | ZINC000452023252 | 100 |
| ZINC000897616680 | | ZINC000897616680 | 93.72 |
| ZINC001196519317 | | ZINC001196519317 | 91 |
| ZINC000131571127 | | ZINC000131571127 | 97.17 |
| ZINC000296612417 | | ZINC000296612417 | 96 |
| ZINC001308961074 | | ZINC001308961074 | 99 |
| ZINC000895657866 | | ZINC000895657866 | 97.68 |
| ZINC000657922756 | | ZINC000657922756 | 98.54 |
| ZINC000657934399 | | ZINC000657934399 | 100 |
| ZINC000549824186 | | ZINC000549824186 | 100 |
| ZINC000769519341 | | ZINC000769519341 | 93.72 |
| ZINC000430988927 | | ZINC000430988927 | 95.71 |
| ZINC000507809396 | | ZINC000507809396 | 100 |
| ZINC000353995714 | | ZINC000353995714 | 100 |
| ZINC000093013680 | | ZINC000093013680 | 100 |
| ZINC000188287346 | | ZINC000188287346 | 96.98 |
| ZINC000453142034 | | ZINC000453142034 | 100 |
| ZINC000924470947 | | ZINC000924470947 | 95.77 |
| ZINC000528002641 | | ZINC000528002641 | 100 |
| ZINC000925887162 | | ZINC000925887162 | 97.17 |
| ZINC000548355486 | | ZINC000548355486 | 92.21 |
| ZINC000574654702 | | ZINC000574654702 | 93.79 |
| ZINC001035320653 | | Z4426348660 | 100 |
| ZINC000996610565 | | Z4426079301 | 100 |
| ZINC001087646081 | | Z4426350008 | 100 |
| ZINC000906421824 | | Z4096780736 | 100 |
| ZINC000948091407 | | Z1212563833 | 100 |
| ZINC000261774189 | | Z2441223595 | 93 |
| ZINC000543048256 | | Z1646405102 | 100 |
| ZINC000182842742 | | Z4426080539 | 100 |
| ZINC001655120594 | | Z3316921677 | 100 |
| ZINC000853031922 | | Z2609960404 | 100 |
| ZINC000649929688 | | Z2277411173 | 99 |
| ZINC000341348768 | | Z2043627283 | 97 |
| ZINC000369129048 | | Z1950347921 | 100 |
| ZINC000582751592 | | Z2878122556 | 100 |
| ZINC000245533477 | | Z1579249600 | 91 |
| ZINC001420054689 | | Z4426350795 | 100 |
| ZINC000416873685 | | Z2360574056 | 95 |
| ZINC000129576345 | | Z1441728492 | 99 |
| ZINC000550829396 | | Z1930703559 | 90 |
| ZINC000389015736 | | Z2148995533 | 90 |
| ZINC000658086473 | | Z2591574640 | 90 |
| ZINC000662800454 | | Z2606802689 | 91 |
| ZINC001078073018 | | Z4426079310 | 100 |
| ZINC000635049325 | | Z2795651844 | 93 |
| ZINC001195393353 | | Z4426350085 | 91 |

### **Supplemental Data Table 7 | Purity information of 11 σ_2_ analogues.**

| ID | | Vendor ID | Purity (%) | |
| --- | --- | --- | --- | --- |
| Z1262980323 | Z1262980323 | | | 100 |
| Z4493924994 | Z4493924994 | | | 100 |
| Z319033458 | Z319033458 | | | 100 |
| Z1484841618 | Z1484841618 | | | 97.32 |
| Z4493948739 | Z4493948739 | | | 100 |
| Z106486854 | Z106486854 | | | 100 |
| Z2272148288 | Z2272148288 | | | 98.57 |
| Z1241145390 | Z1241145390 | | | 100 |
| Z2002520638 | Z2002520638 | | | 100 |
| Z1262978490 | Z1262978490 | | | 100 |
| Z1491031116 | Z1491031116 | | | 100 |
| Z4493924961 | Z4493924961 | | | 98.8 |
| Z1241145825 | Z1241145825 | | | 92.05 |
| Z1348522601 | Z1348522601 | | | 92.8 |
| Z4493929232 | Z4493929232 | | | 95.59 |
| Z4493948717 | Z4493948717 | | | 95.41 |
| Z1262978782 | Z1262978782 | | | 100 |
| Z2272647011 | Z2272647011 | | | 100 |
| Z1262979110 | Z1262979110 | | | 100 |
| Z1241146719 | Z1241146719 | | | 94.76 |
| Z1241144480 | Z1241144480 | | | 90.5 |
| Z1230400045 | Z1230400045 | | | 100 |
| Z2102236109 | Z2102236109 | | | 100 |
| Z4493924963 | Z4493924963 | | | 95.8 |
| Z1348458155 | Z1348458155 | | | 90.36 |
| Z1241145220 | Z1241145220 | | | 91.07 |
| Z1262980692 | Z1262980692 | | | 96.14 |
| Z1230400157 | Z1230400157 | | | 100 |
| Z1540927861 | Z1540927861 | | | 100 |
| Z1241144621 | Z1241144621 | | | 98.57 |
| Z1241143871 | Z1241143871 | | | 100 |
| Z1241143959 | Z1241143959 | | | 93.16 |
| Z1241144449 | Z1241144449 | | | 97.67 |
| Z4493929224 | Z4493929224 | | | 91 |
| Z1230400226 | Z1230400226 | | | 95.25 |
| Z4493929235 | Z4493929235 | | | 100 |
| Z4493924960 | Z4493924960 | | | 100 |
| Z1241147220 | Z1241147220 | | | 91 |
| Z2102296558 | Z2102296558 | | | 100 |
| Z1241145790 | Z1241145790 | | | 93.34 |
| Z1612280219 | Z1612280219 | | | 100 |
| Z4493929234 | Z4493929234 | | | 90.23 |
| Z1492827933 | Z1492827933 | | | 99 |
| Z1241147223 | Z1241147223 | | | 99 |
| Z1241144947 | Z1241144947 | | | 98 |
| Z1905110467 | Z1905110467 | | | 91 |
| Z1665385680 | Z1665385680 | | | 91.2 |
| Z2947867435 | Z2947867435 | | | 91 |
| Z1241143574 | Z1241143574 | | | 96.83 |
| Z1567294634 | Z1567294634 | | | 92.57 |
| Z1562881018 | Z1562881018 | | | 91.05 |
| Z1262980068 | Z1262980068 | | | 100 |
| Z2272980237 | Z2272980237 | | | 95 |
| Z1665845742 | Z1665845742 | | | 93.4 |
| Z1262979330 | Z1262979330 | | | 90.74 |
| Z1262980403 | Z1262980403 | | | 97.17 |
| Z4493924957 | Z4493924957 | | | 97.12 |
| Z2271852773 | Z2271852773 | | | 90.35 |
| Z4493948738 | Z4493948738 | | | 93.32 |
| 5486num2758.1 | 5486num2758.1 | | | 98.01 |
| 5486num2766 | 5486num2766 | | | 100 |
| 5486num2768.2 | 5486num2768.2 | | | 100 |
| 5486num2765 | 5486num2765 | | | 100 |
| Z2146804677 | Z2146804677 | | | 100 |
| Z1601026740 | Z1601026740 | | | 100 |
| Z1653918014 | Z1653918014 | | | 100 |
| Z31347727 | Z31347727 | | | 100 |
| 5486num2770 | 5486num2770 | | | 99 |
| 7866num2119.2 | 7866num2119.2 | | | 92.46 |
| 7866num1944.2 | 7866num1944.2 | | | 95 |
| 7866num3311.1 | 7866num3311.1 | | | 100 |
| 5486num2704.1 | 5486num2704.1 | | | 93.72 |
| 7866num3320.1 | 7866num3320.1 | | | 96.04 |
| 5486num3075.1 | 5486num3075.1 | | | 95.13 |
| 7866num1973.2 | Z4857158944 | | | 100 |
| 5486num2642 | 5486num2642 | | | 94.74 |
| 5486num913 | 5486num913 | | | 91.48 |
| Z1665798906 | Z1665798906 | | | 97 |
| Z163048780 | Z163048780 | | | 100 |
| Z295861754 | Z295861754 | | | 92 |
| 5486num495 | Z4510627693 | | | 94.99 |
| 5486num483 | Z4510627677 | | | 93.74 |
| 5486num940 | Z4510627684 | | | 97.71 |
| 5486num943 | Z4510627675 | | | 95.12 |
| 5486num60 | Z4510627674 | | | 90.16 |
| 5486num914 | Z4510627673 | | | 95.89 |
| Z4510627683 | Z4510627683 | | | 96.83 |
| 7866num575.2 | Z4510628118 | | | 93.06 |
| 5486num484 | Z4510627678 | | | 100 |
| 5486num302.2 | Z1504291172 | | | 95.2 |
| 5486num493 | Z4510627692 | | | 100 |
| 5486num2735 | Z4510627682 | | | 93.46 |
| 5486num570.2 | Z4510628128 | | | 95 |
| 7866num567.2 | Z4510628193 | | | 92.63 |
| 5486num2751 | Z4510627688 | | | 93.38 |
| 5486num1704.1 | Z4510628228 | | | 90 |
| 5486num2756 | Z4510627687 | | | 90.31 |
| Z4510627652 | Z4510627652 | | | 96.31 |
| 7866num560.2 | Z4510628119 | | | 98.44 |
| 7866num558.2 | Z4510628191 | | | 100 |
| 5486num911 | Z4510627672 | | | 97.93 |
| 7866num2155.1 | Z4510628285 | | | 100 |
| 5486num2260 | Z4510627694 | | | 100 |
| 5486num156 | Z4510627671 | | | 91.01 |
| Z4510627653 | Z4510627653 | | | 95.31 |
| 7866num672.2 | Z4510628117 | | | 98.05 |
| Z4430418526 | Z4430418526 | | | 100 |
| 7866num2.2 | Z4510628326 | | | 100 |
| Z4430417979 | Z4430417979 | | | 100 |
| 7866num562.2 | Z4510628137 | | | 96.51 |

### **Supplemental Data Table 8 | Purity information of 3 potent σ_2_ ligands for the animal model of neuropathic pain.**

| ID | Vendor ID | Purity (%) |
| --- | --- | --- |
| Z4765809799 | Z4765809799 | 95 |
| Z4857158944 | Z4857158944 | 98 |
| Z4634276811 | Z4634276811 | 95 |

### **Figures**

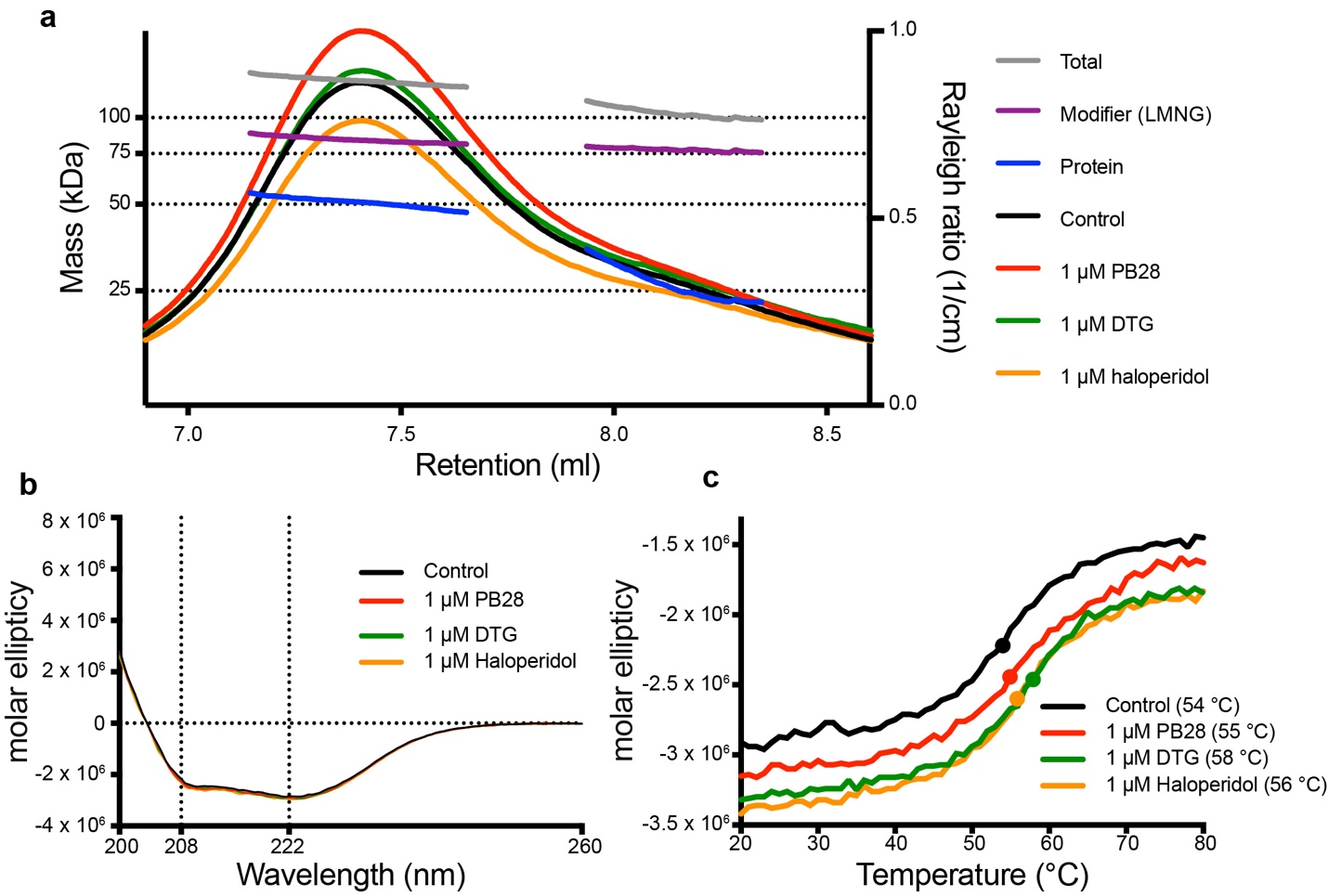

**Supplementary Information Figure 1 | Oligomeric state and stability of the σ_2_ receptor. a**, Size-exclusion chromatography with multi-angle light scattering of the human σ_2_ receptor. The σ_2_ receptor was run either w/o ligand or with 1 µM of the indicated ligand. Lines indicate calculated total mass (gray), detergent micelle (blue), and protein (purple). **b**, Circular dichroism analysis of the bovine σ_2_ receptor alone (black) or with the indicated ligand. **c**, Circular dichroism melting curves of the bovine σ_2_ receptor. Temperature was raised from 20 ºC to 95 ºC and molar ellipticity was measured at 222 nm. Protein was incubated either with or without indicated ligand at 1 µM. Melting temperatures for each measurement are indicated.

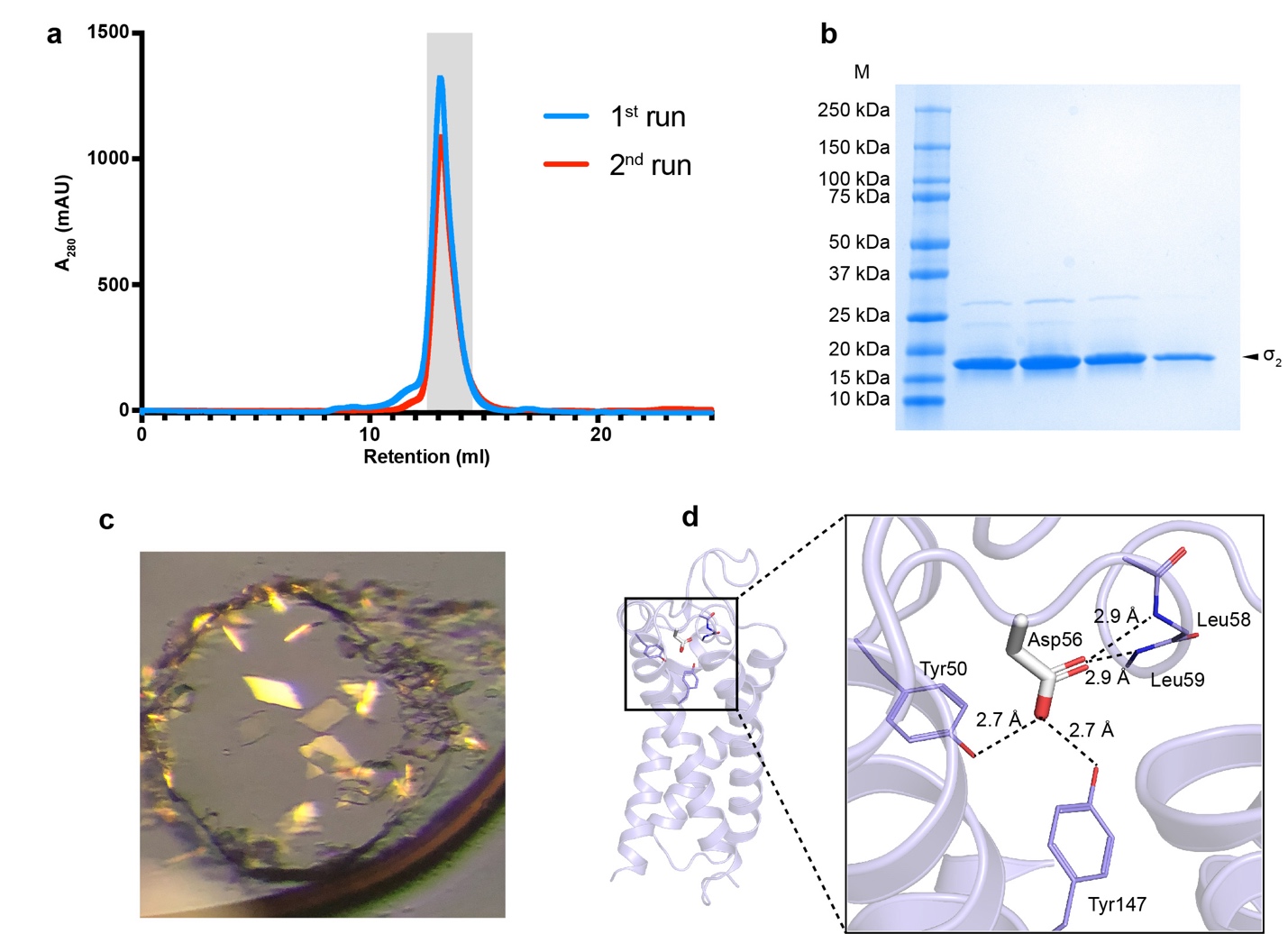

**Supplementary Information Figure 2 | Purification and crystallization of the σ_2_ receptor. a**, Size-exclusion chromatography (SEC) of the σ_2_ receptor. Blue trace is after proteolytic tag removal. Red trace is protein applied on size exclusion after reapplying the tag-free protein on affinity resin to remove proteins with intact tags. **b**, Analysis of receptor purity after the second SEC using SDS-PAGE. Gray rectangle in **a** represents fractions chosen for analysis. **c**, Crystals of bovine σ_2_ receptor in the lipidic cubic phase. **d**, Residue Asp56 is important for receptor structure but not for ligand binding.

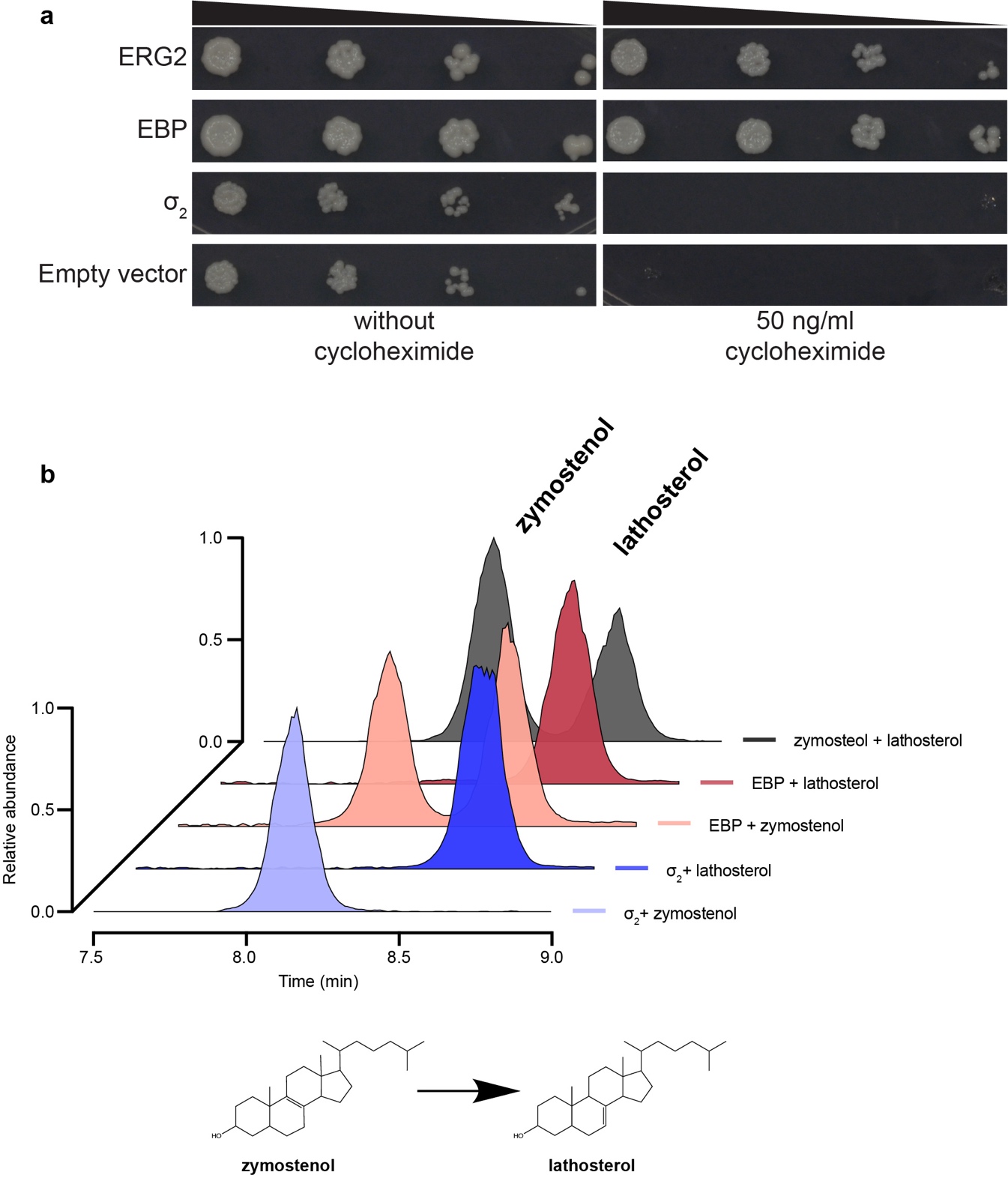

**Supplementary Information Figure 3 | The σ_2_ receptor does not function as a Δ8-9 sterol isomerase. a**, Yeast complementation assay. ERG2 and EBP can act as sterol isomerases and rescue the growth of ΔERG2 *Saccharomyces cerevisiae* while the σ_2_ receptor cannot. Yeast were grown either in permissive condition of no cycloheximide or in the restrictive conditions of 50 ng/ml cycloheximide, which requires functional Δ8-9 sterol isomerase activity for viability. **b**, EBP can catalyze to conversion of zymostenol to lathosterol while σ_2_ cannot. Standards are in dark gray. EBP converts zymostenol to lathosterol (apricot) but does not convert lathosterol to zymostenol (dark red). The σ_2_ receptor does not convert lathosterol to zymostenol (dark blue) or zymostenol to lathosterol (light purple).

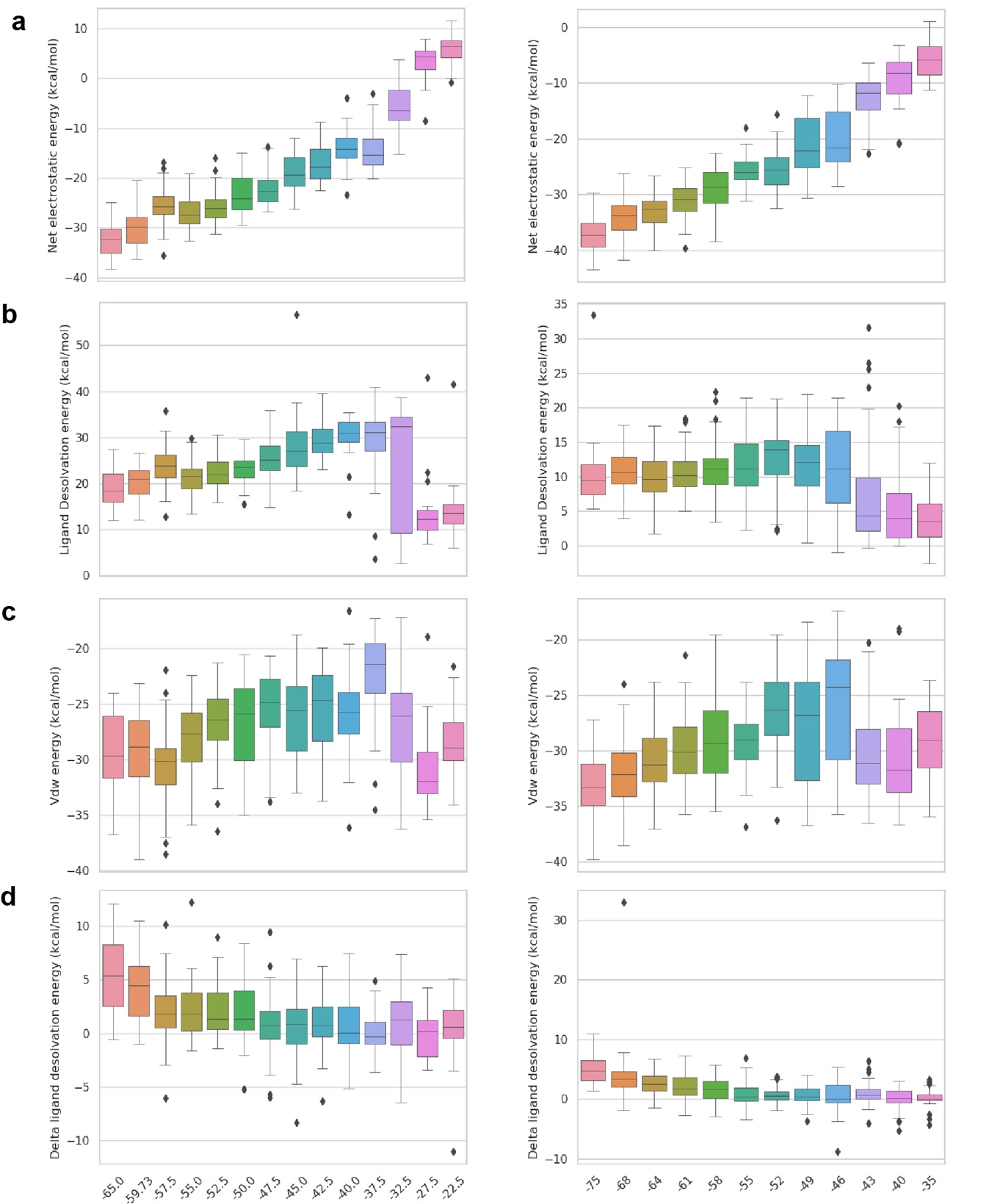

**Supplementary Information Figure 4 | The distribution of docking scores of tested molecules for hit rate curves against σ_2_ (the left column) and D_4_ (the right column) receptors.** All tested molecules are grouped based on docking score bins. The distributions are shown in box plots for **a**, net electrostatic energy, **b,** ligand desolvation energy, **c,** van der Waals (vdW) energy and **d,** delta ligand desolvation energy after recalculating atomic desolvation energy based on the docked pose.

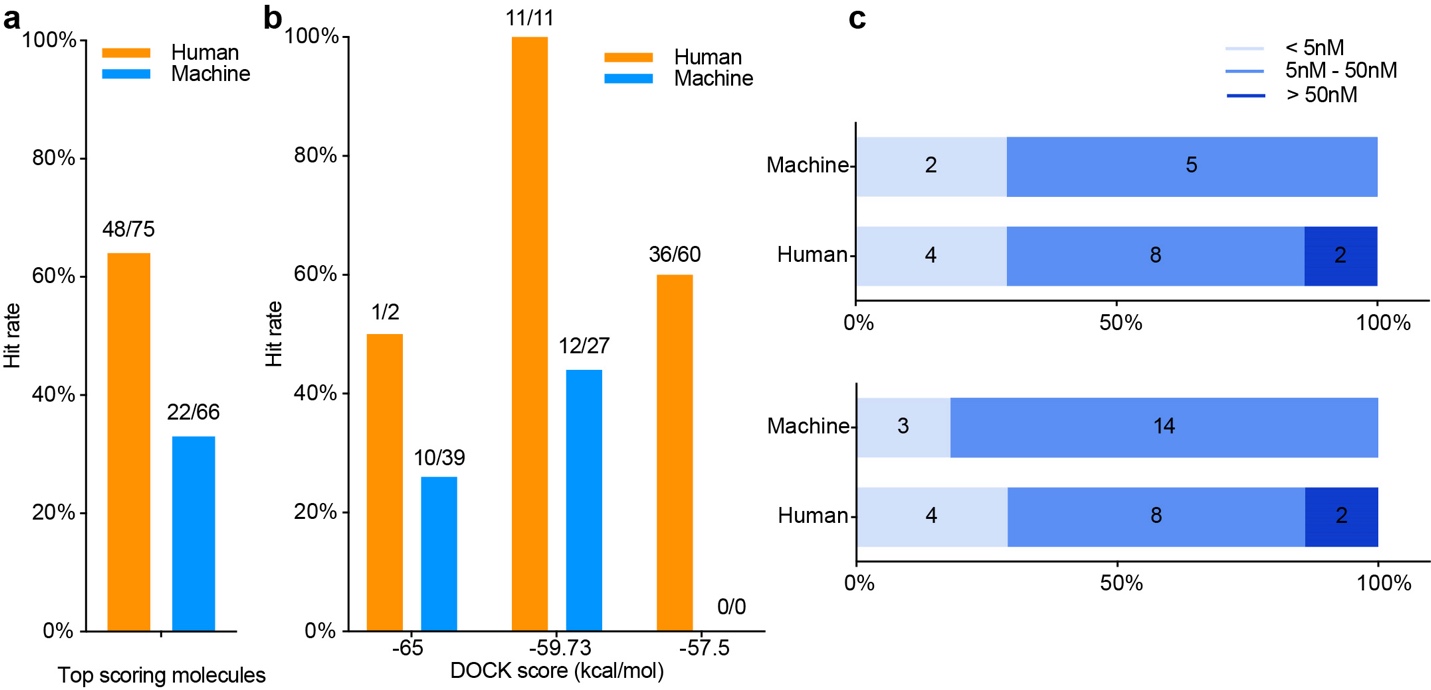

**Supplementary Information Figure 5 | Comparison of hit rates and affinities achieved by combined docking score and human inspection and these achieved by docking score alone. a**, The overall hit rates for selecting compounds from the first 3 scoring bins by each strategy: human prioritization and docking score (orange), or docking score alone (blue). Hit rate is the ratio of active compounds/tested compounds; the raw numbers appear at the top of each bar. **b**, the hit rates for selecting compounds at different scoring ranges by each strategy: human prioritization and docking score (orange), or docking score alone (blue). **c**, the upper panel, distribution of the binding affinity level among the hits from **a**. We measured concentration-response curves for 14 docking hits from human prioritization and docking score, and 7 hits from the docking score alone. These are divided into three affinity ranges: <5 nM (pale blue); 5 nM–50 nM (blue); >50 nM (dark blue); the bottom panel, distribution of the binding affinity level among the hits from all different scoring ranges. We measured concentration-response curves for 14 docking hits from human prioritization and docking score, and 17 hits from the docking score alone.

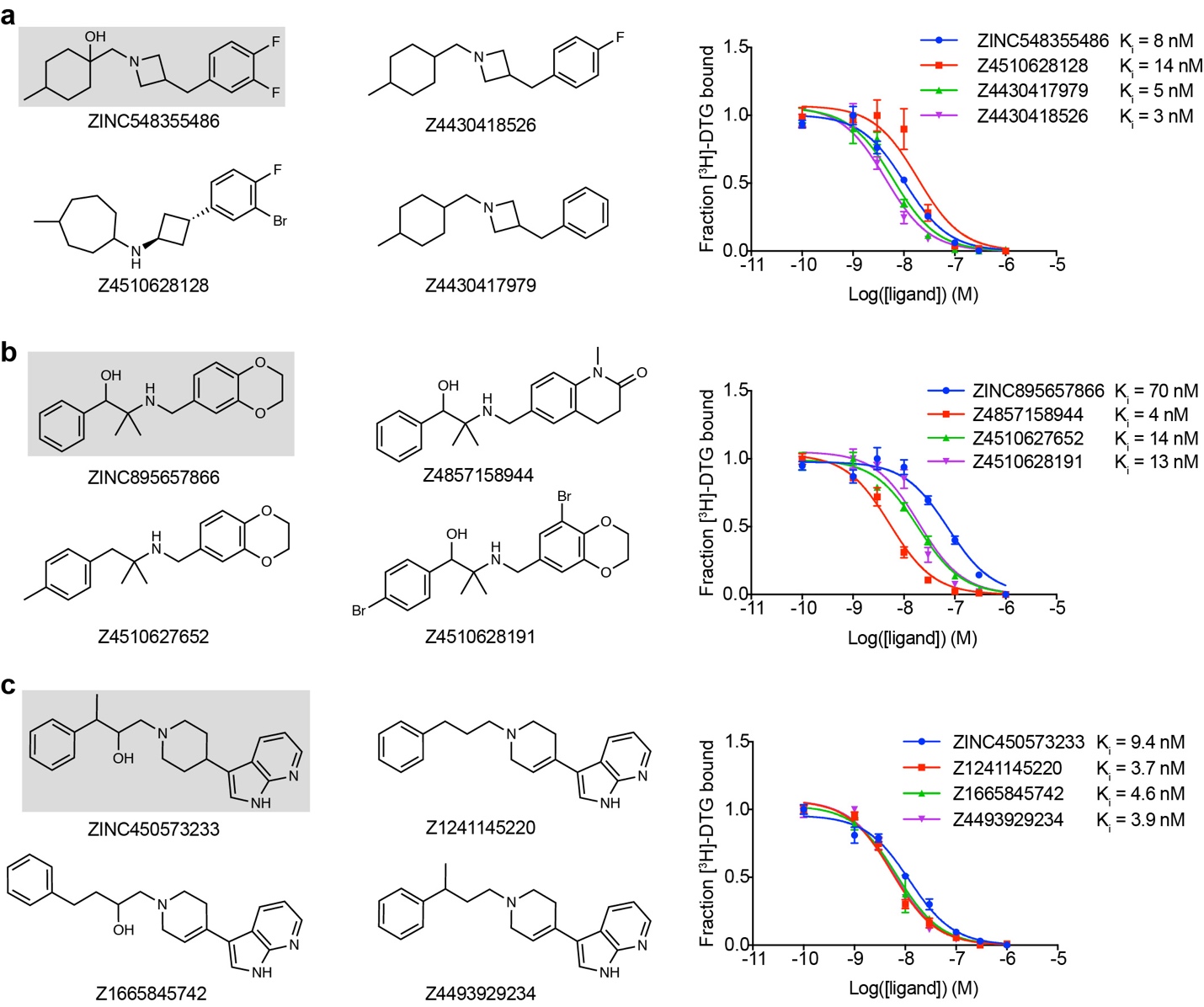

**Supplementary Information Figure 6 | Initial hits and selected analogs of σ_2_ receptor ligands.** **a**, Parent compound ZINC548355486 indicated by gray background and its three potent analogues (2D drawings on the left panel and binding curves on the right panel). **b**, Parent compound ZINC895657866 indicated by gray background and its three potent analogues (2D drawings on the left panel and binding curves on the right panel). **c**, Parent compound ZINC450573233 indicated by gray background and its three potent analogues (2D drawings on the left panel and binding curves on the right panel).

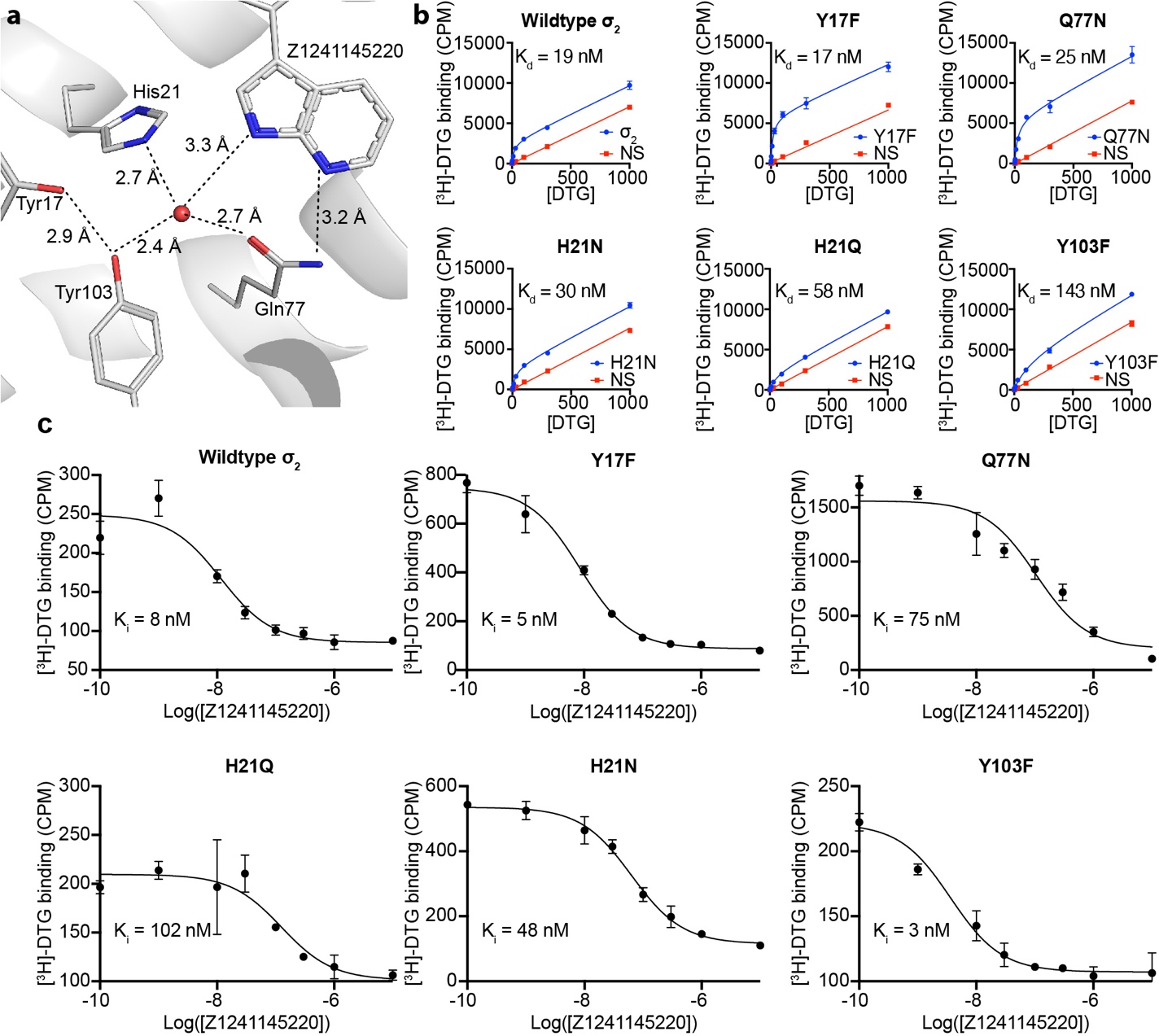

**Supplementary Information Figure 7 | The binding site of the σ_2_ receptor contains a structural water. a**, Water coordination at the binding site of the σ_2_ receptor. Water molecule is depicted as a red sphere. Hydrogen bonds are indicated by dashed line. **b**, Measurement of the dissociation constant (K_d_) of [^3^H]-DTG for the various mutants of σ_2_ receptor meant to disrupt water coordination. Residues proximal to the structural water were chosen for mutation. Residues were mutated to the indicated amino acid. **c**, Inhibition constants (K_i_) measurements of Z1241145220 in various mutants of σ_2_.

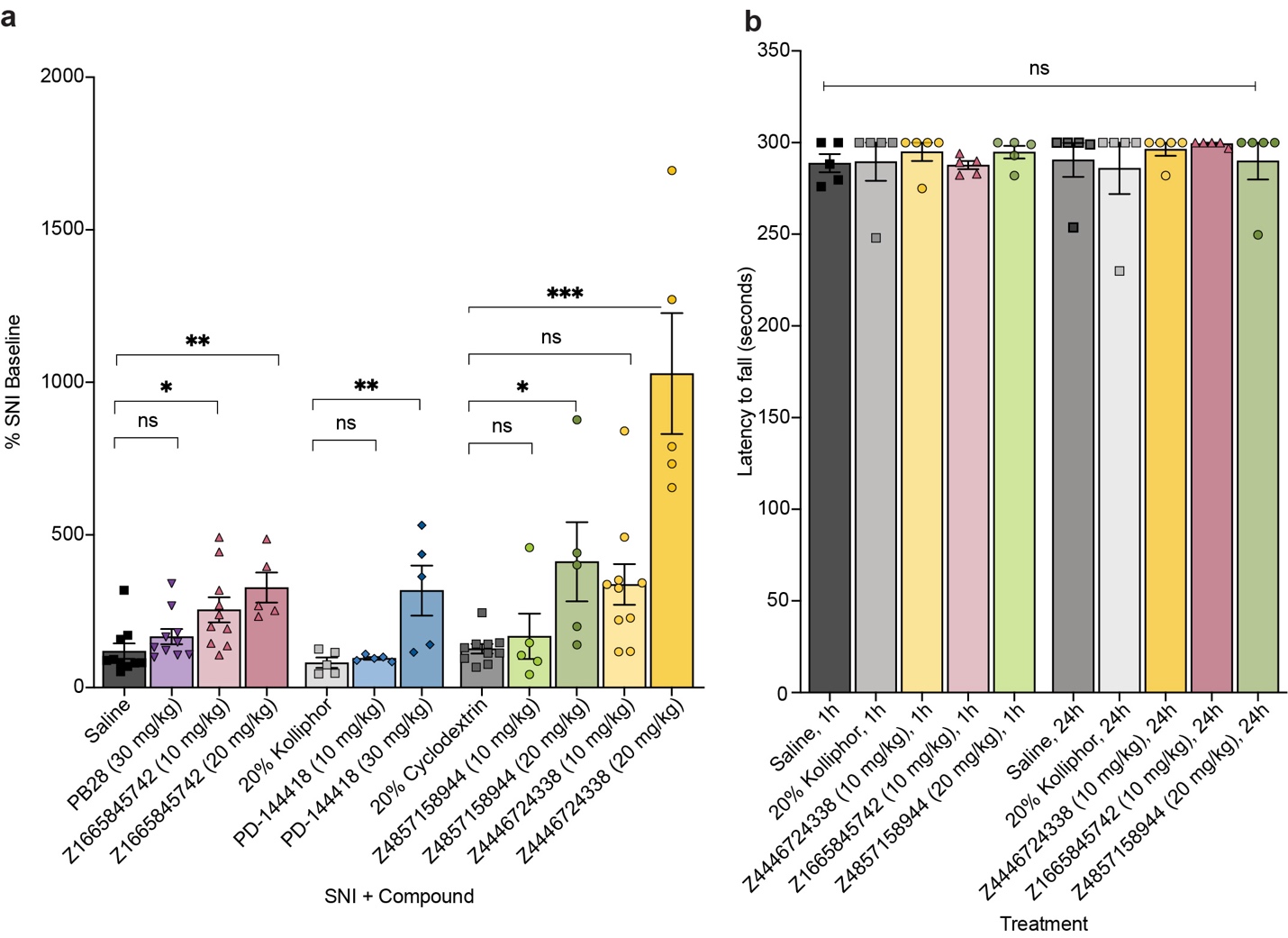

**Supplementary Information Figure 8 | Effect of systemic σ receptor ligands on motor behavior. a,** Response of mice to a Von Frey filament after spared nerve injury (SNI). All five ligands are compared to their respective vehicles (PD-144418 10 mg/kg (*n* = 5) and 30 mg/kg (*n* = 5) vs. kolliphor (*n* = 5), one-way ANOVA, *F*(2, 12) = 7.49, *p* = 0.008; Z4446724338 10 mg/kg (*n* = 10) and 20 mg/kg (*n* = 5) vs cyclodextrin (*n* = 10), one-way ANOVA, *F*(2, 22) = 25.12, *p* < 0.001; Z4857158944 10 mg/kg (*n* = 5) and 20 mg/kg (*n* = 5) vs cyclodextrin (*n* = 10), one-way ANOVA, *F*(2, 17) = 5.10, *p* = 0.02; Z1665845742 10 mg/kg (*n* = 10) and 20 mg/kg (*n* = 5) and PB28 30 mg/kg (*n* = 10) vs saline (*n* = 10), one-way ANOVA, *F*(3, 31) = 6.18, *p =* 0.002; asterisks define individual group differences to respective vehicle control using Dunnett’s multiple comparisons Post-hoc test; ns = not significant, * *p* < 0.05, ** *p* < 0.01, *** *p* < 0.001). Data shown are mean ± SEM. Data for higher doses and vehicles is replotted from **Figure 4**. **b,** No sedation or motor impairment on the rotarod was observed after drug treatments compared to vehicle at 1 hour (Z1665845742 10 mg/kg (*n* = 5) and Z4857158944 20 mg/kg (*n* = 5) vs saline (*n* = 5), one-way ANOVA, *F*(2, 12) = 1.04, *p* = 0.38; Z4446724338 10 mg/kg (*n* = 5) vs kolliphor (*n* = 5), unpaired two-tailed Student’s *t*-test, *t*(8) = 0.47, *p* = 0.65) or 24 hours post-injection (Z1665845742 10 mg/kg (*n* = 5) and Z4857158944 20 mg/kg (*n* = 5) vs saline (*n* = 5), one-way ANOVA, *F*(2, 12) = 0.45, *p* = 0.65; Z4446724338 10 mg/kg (*n* = 5) vs kolliphor (*n* = 5), unpaired two-tailed Student’s *t*-test, *t*(8) = 0.72, *p* = 0.49); ns = not significant. Data shown are means ± SEM.

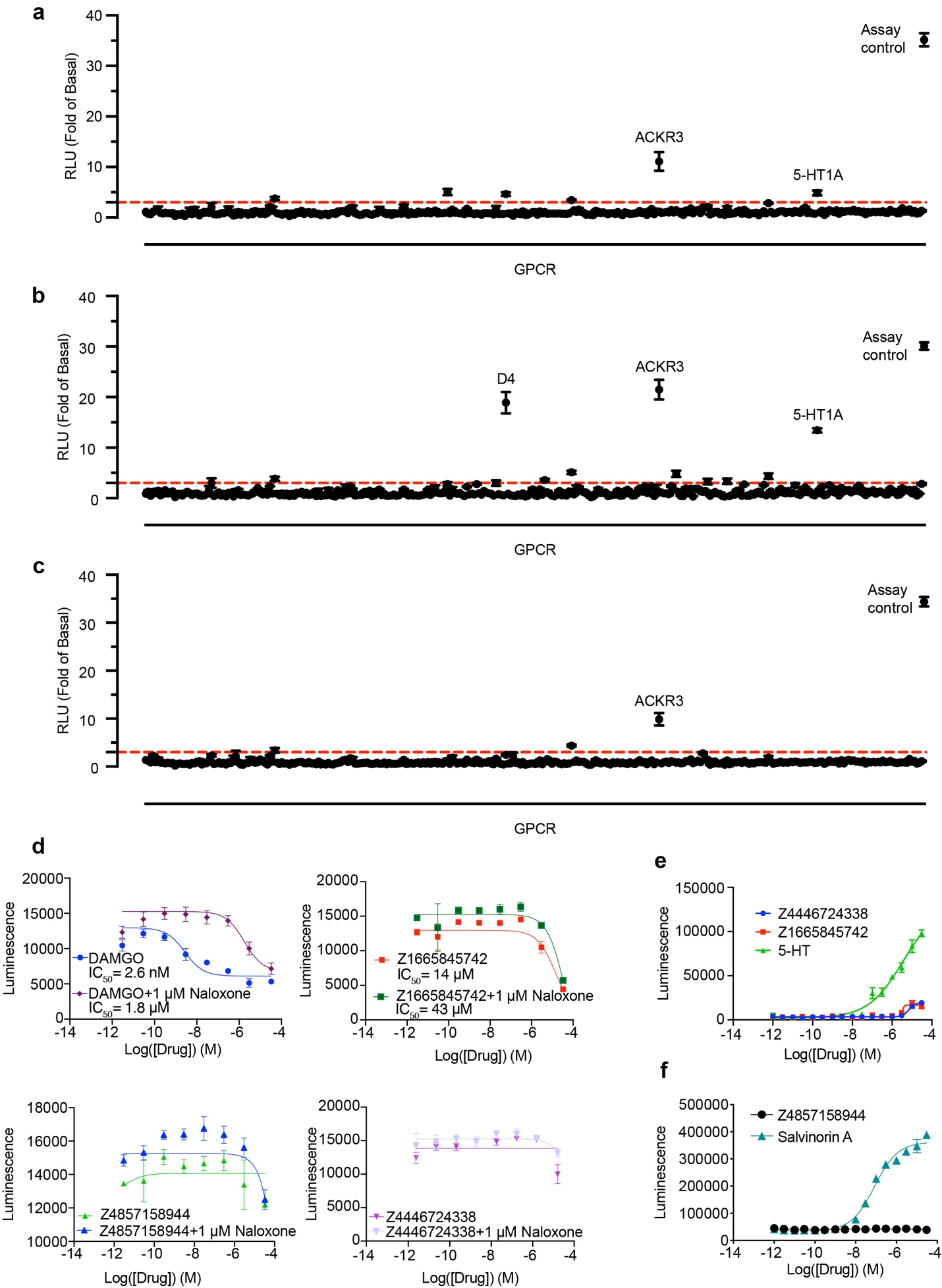

**Supplementary Information Figure 9 | The off-target profiling of Z4446724338, Z1665845742 and Z4857158944.** TANGO screens against a panel of 320 GPCRs for σ_2_ ligands **a**, Z4446724338, **b**, Z1665845742 and **c**, Z4857158944. **d**, GloSensor μOR-mediated cAMP inhibition (G_i_ activation) by DAMGO, Z4446724338, Z1665845742 and Z4857158944. Follow-up does-response curves for pain-related receptors that showed activation in GPCRome: **e**, Z4446724338 and Z1665845742 against 5HT1A; **f**, Z4857158944 against κOR.

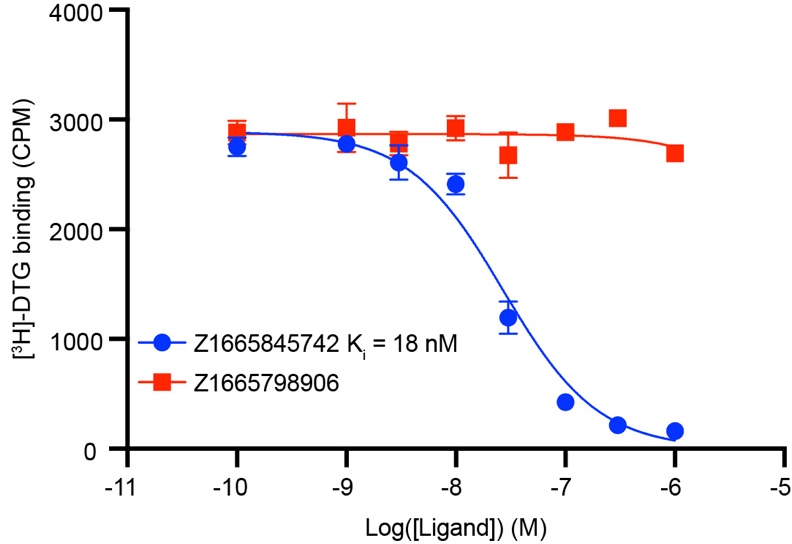

**Supplementary Information Figure 10 | Concentration-response binding curves for the probe pair molecules.** The data are the mean ± SEM from three technical replicates.

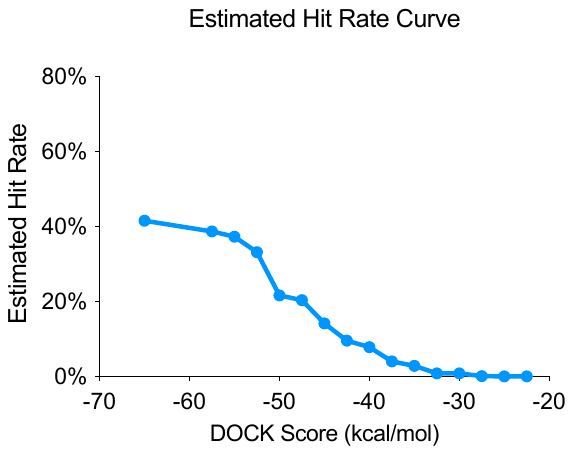

### **Supplementary Information Figure 11 | The estimated hit-rate was plotted against docking energy.**

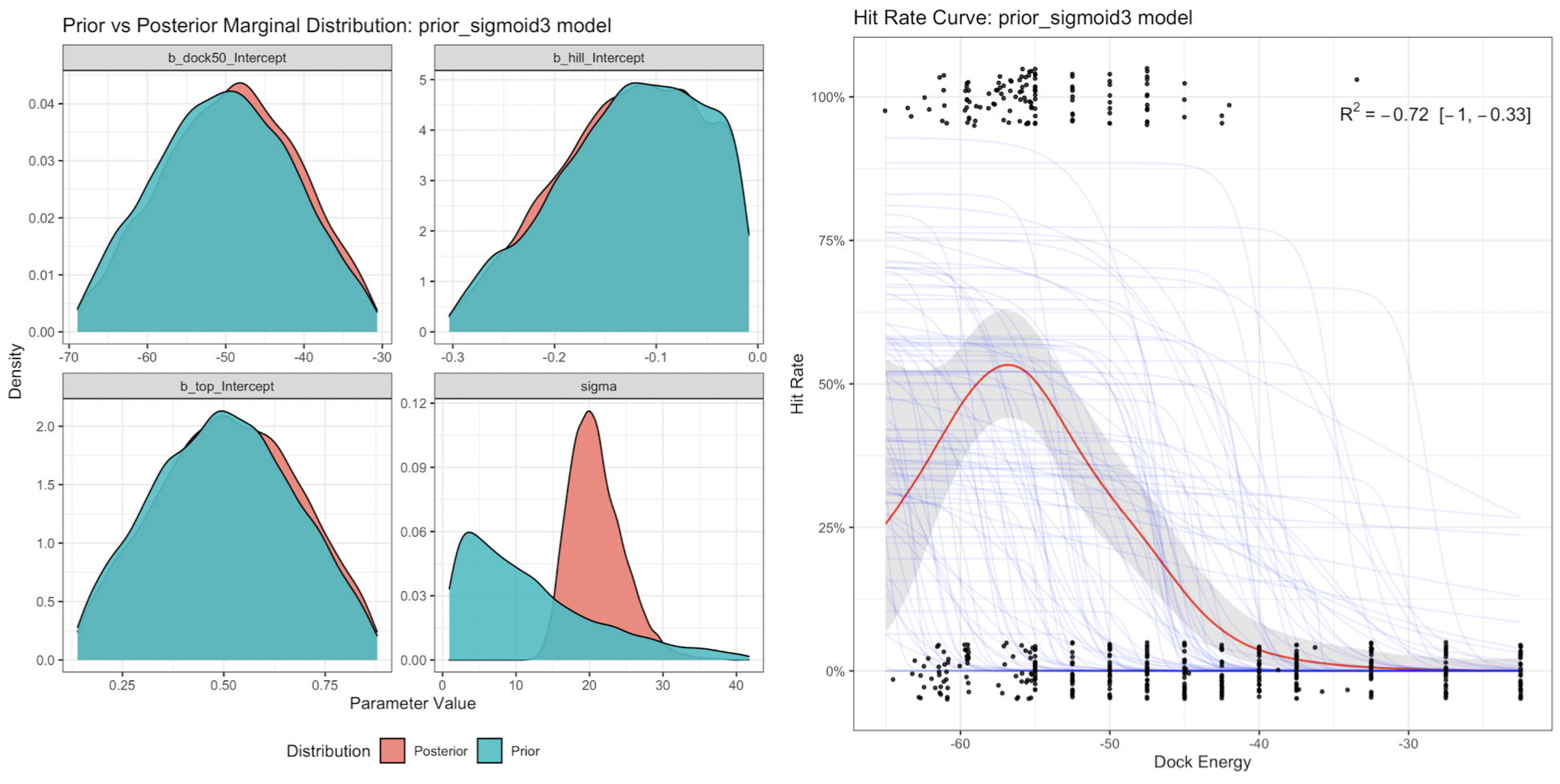

### **Supplementary Information Figure 12 | Prior and posterior parameter distributions for the prior sigmoid model.**
